# Host microRNA-31-5p represses oncogenic herpesvirus lytic reactivation by restricting the RNA-binding protein KHDRBS3-mediated viral gene expression

**DOI:** 10.1101/2025.01.22.634336

**Authors:** Soo Mi Lee, Christopher L. Avalos, Christos Miliotis, Hanna M. Doh, Erica Chan, Kenneth M. Kaye, Frank J. Slack

## Abstract

Oncogenic Kaposi’s sarcoma-associated herpesvirus (KSHV), an etiological agent of Kaposi’s sarcoma and primary effusion lymphoma, employs a biphasic life cycle consisting of latency and lytic replication to achieve lifelong infection. Despite its essential role in KSHV persistence and tumorigenicity, much remains unknown about how KSHV lytic reactivation is regulated. Leveraging high-throughput transcriptomics, we identify microRNA-31-5p (miR-31-5p) as a key regulator of KSHV lytic reactivation capable of restricting KSHV entry into the lytic replication cycle. Ectopic expression of miR-31-5p impairs KSHV lytic gene transcription and production of lytic viral proteins, culminating in dramatic reduction of infectious virion production during KSHV reactivation. miR-31-5p overexpression also markedly reduces the expression of critical viral early genes, including the master regulator of the latent-lytic switch, KSHV replication and transcription activator (RTA) protein. Through mechanistic studies, we demonstrate that miR-31-5p represses KSHV lytic reactivation by directly targeting the KH domain protein KHDRBS3, an RNA-binding protein known to regulate RNA processing including alternative splicing. Our study highlights KHDRBS3 as an essential proviral host factor that is key to the successful completion of KSHV lytic replication and suggests its novel function in viral lytic gene transcription during KSHV reactivation. Taken together, these findings reveal a previously unrecognized role for the miR-31-5p/KHDRBS3 axis in regulating the KSHV latency-lytic replication switch and provide insights into gene expression regulation of lytic KSHV, which may be leveraged for lytic cycle-targeted therapeutic strategies against KSHV-associated malignancies.

## Introduction

The DNA virus Kaposi’s sarcoma-associated herpesvirus (KSHV) is an oncogenic herpesvirus that is linked directly to the development of Kaposi’s sarcoma and primary effusion lymphoma^1–3^. KSHV employs two alternative life cycles, the latent phase and the lytic replication phase, to accomplish lifelong infection. Upon reactivation from latency, KSHV enters the lytic replication phase, which is characterized by an intricate cascade of lytic gene expression that integrates viral DNA replication and structural protein synthesis, resulting in the generation of viable viral particles^4^. Thus, periodic reactivation is essential for viral transmission, tumorigenicity, and for the establishment and maintenance of latent viral reservoirs^4,6^. While environmental cues can induce reactivation, mechanisms that control the KSHV latency-lytic replication switch are not fully understood. Understanding the molecular mechanisms governing successful activation and progression of the KSHV lytic phase is of the utmost importance for developing lytic cycle-targeted therapy as a novel treatment strategy for KSHV-associated diseases.

KSHV lytic genes are classified into three major kinetic classes (immediate-early (IE), early (E), and late (L) genes) which are expressed in a highly coordinated sequential cascade. IE genes encode mainly transcriptional activators that trigger this lytic gene cascade from latency. Notably, the IE protein replication and transcription activator (RTA) encoded by the ORF50 gene acts as a master regulator of the latent-to-lytic switch that can initiate and drive KSHV lytic cycle to completion. As an IE gene, RTA activates a broad spectrum of lytic viral genes, including ORF57, K8, *polyadenylated nuclear (PAN)* RNA, and RTA itself^4,6^. Of note, *PAN* RNA, a key viral long noncoding RNA (lncRNA) required for L gene expression and virion production, becomes the most abundant polyadenylated transcript in the infected host cell during KSHV lytic reactivation^5^. Although the majority of *PAN* RNA is predominantly localized in the nucleus, substantial levels of this lncRNA have been detected in the cytoplasm^5^. IE and E gene products necessary for viral DNA replication are first transcribed, followed by viral DNA replication, and L genes encoding structural proteins required for assembly and release of infectious particles are transcribed, ultimately leading to the production of new KSHV virions^4,6^. During the KSHV lytic phase, innate antiviral immune responses are activated to restrict lytic replication. In contrast, during latency, KSHV exists in a quiescent, immunologically silent state in which no progeny virions are produced and lytic genes are tightly repressed, with only a few viral genes expressed from the latency locus, including LANA (ORF73), vCyclin (ORF72), vFLIP (ORF71), and multiple viral miRNAs^4,6,7^.These latent proteins antagonize and evade host immune surveillance to establish and support a lifelong, persistent KSHV infection in the host^7^. Moreover, KSHV miRNAs have been shown to facilitate viral latency by repressing lytic gene expression^8–10^. Like all herpesviruses, KSHV maintains in the latent phase in the infected host following primary infection, and as a result, most KSHV-associated tumors are latently infected and only a small percentage of cells undergo lytic replication, which is integral to disease progression and dissemination^11,12^. The balance between latent and lytic KSHV replication cycles is stringently controlled to ensure viral persistence and consequently tumor development^4,6^.

MicroRNAs (miRNAs) are a class of highly conserved ∼23 nucleotide (nt) noncoding RNAs that play crucial roles in post-transcriptional gene regulation. They regulate gene expression by directly binding to target mRNAs, typically in the 3’ untranslated regions (UTRs) through a target site complementary to the miRNA seed (miRNA nucleotides 2-7 from the 5’ end), to induce mRNA decay and/or translational repression^13^. miRNAs have emerged as critical regulators of diverse developmental, cellular, and physiological processes, and their dysregulation has been causally linked to many pathological conditions, including virus infections and virus-induced malignancies. There is increasing evidence that host and viral miRNAs control multiple biological processes relevant to infection, ranging from virus replication and propagation to host antiviral responses, through complex regulatory mechanisms affecting various stages of the viral life cycle^14–16^. However, relatively little is known about the roles of host miRNAs in the initiation and regulation of KSHV lytic reactivation.

Here, we sought to investigate functional roles for host miRNAs in regulating the transition from latent infection to lytic replication. Leveraging high-throughput miRNA-mRNA transcriptomics during the KSHV lytic replication phase in KSHV-infected cells, we identify host miR-31-5p as a potent regulator of KSHV lytic reactivation. We show that ectopic expression of miR-31-5p restricts KSHV lytic gene transcription and viral protein expression, ultimately resulting in a significant reduction in infectious virion production during KSHV reactivation. Overexpression of miR-31-5p also results in markedly reduced expression of the master regulator of the latent-lytic switch, KSHV replication and transcription activator (RTA) protein. Through mechanistic studies, we demonstrate that the observed effects of miR-31-5p is mediated through direct repression of KH RNA binding domain containing, signal transduction associated 3 (KHDRBS3), an RNA-binding protein known to control alternative splicing^17–21^ and implicated in other RNA processing events, such as RNA stability^22^ or export^22^. Our results suggest that KHDRBS3 serves as a proviral factor for KSHV lytic reactivation and facilitates oncogenic herpesvirus lytic replication through its function in viral gene transcription. Accordingly, KSHV lytic gene expression and infectious virion production are significantly impaired by KHDRBS3 silencing. This study therefore highlights the RNA-binding protein KHDRBS3 as a critical host factor that is key to the successful completion of KSHV lytic replication and uncovers a previously unrecognized role of the miR-31-5p/KHDRBS3 axis in transcriptional regulation of KSHV lytic gene expression. These findings may be leveraged for developing novel therapeutic strategies targeting the viral latent-lytic switch for the treatment of KSHV-associated malignancies.

## Results

### Cellular miRNA expression landscape during KSHV lytic reactivation from latency

To characterize changes in cellular miRNA expression during KSHV lytic reactivation, we utilized a well-established model of KSHV reactivation, the doxycycline (Dox)-inducible KSHV producer cell line iSLK.219, which contains a Dox-inducible KSHV lytic switch protein RTA to mediate efficient reactivation and entry into the lytic cycle upon Dox treatment. These cells also harbor a recombinant KSHV (rKSHV.219) genome featuring GFP expression driven by the constitutive elongation factor 1 (EF-1) promoter as an indicator for latently infected cells, and inducible RFP expression driven by the KSHV early lytic gene PAN promoter as an indicator for early lytic reactivated cells^24^. Thus, a constitutive GFP marker can be used as a latent infection maker, while an RFP marker can be used as an early lytic reactivation marker. These iSLK.219 cells were subjected to Dox treatment to induce RTA expression, resulting in increased expression of RFP driven by the viral lytic PAN promoter over a 120 h time course of KSHV lytic reactivation (Supplementary Fig. 1a). As expected, protein expression of the IE ORF45 viral lytic protein was robustly induced in lytic iSLK.219 cells (RFP-positive) but was undetectable in latent iSLK.219 and other KSHV-latently infected cell lines, indicating a successful transition from a dormant latent phase to an active lytic replication phase of the virus (Supplementary Fig. 1b).

To identify host miRNAs that play an important role in regulating KSHV reactivation and/or progression of the lytic replication phase, we performed small-RNA sequencing of RNA isolated from latent KSHV-infected iSLK.219 cells at 0 h or lytic iSLK.219 cells at 72 h post Dox-induced KSHV lytic reactivation. To rule out the possibility that Dox itself can affect miRNA expression, we also performed small-RNA sequencing of RNA isolated from KSHV-negative iSLK cells, which lacks the KSHV genome but harbors the Dox-inducible RTA transgene, without Dox treatment at 0 h or at 72 h post-Dox treatment. A list of differentially expressed cellular miRNAs (n = 36) relevant to KSHV lytic reactivation was obtained from this analysis (Supplementary Table 2). The top down-regulated miRNAs included miR-31-5p, miR-29a-3p, miR-181a-3p, miR-194-5p, and miR-449c-5p and the top up-regulated miRNAs included miR-139-5p, miR-7-5p, miR-210-3p and miR-3065-5p (Fig. 1a, b). We confirmed that Dox treatment in iSLK cells did not significantly down-regulate or up-regulate these cellular miRNAs, indicating that Dox itself or RTA transgene expression alone does not affect the levels of these miRNAs (data not shown). In addition, 8 KSHV-encoded miRNAs, including pre-miR-K12-12, were significantly up-regulated upon lytic reactivation in iSLK.219 cells (FC ≥ 2) (Supplementary Fig. 2 and Supplementary Table 3). Notably, four out of the top five down-regulated host miRNAs, miR-29a-3p, miR-181a-3p, miR-194-5p, and 449c-5p have been implicated in restricting the infection and replication of influenza virus (IV)^25^, rotavirus (RV)^26^, herpes simplex virus (HSV)^27^, human immunodeficiency virus (HIV)^28^ and the closely related simian immunodeficiency virus (SIV)^28^, as well as attenuating disease pathogenesis of Epstein-Barr virus (EBV)^29^, human papillomavirus (HPV)^30^, and West Nile virus (WNV)^31^. Moreover, the highly conserved miR-449 family consisting of three miRNAs (miR-449a, miR-449b, and miR-449c) is important for regulation of virus-host interactions by modulating virus replication and miR-449a, which shares the same seed sequence as miR-449c, is down-regulated by hepatitis B virus (HBV) and hepatitis C virus (HCV) infection^32^. However, the role of miR-31-5p (hereafter referred to as miR-31) in the context of viral infections, replication and viral pathogenesis remains largely uncharacterized. Therefore, we sought to investigate the role of this host miRNA in orchestrating KSHV reactivation and lytic replication, which is an essential pathogenic step in multiple human diseases.

**Figure 1.**
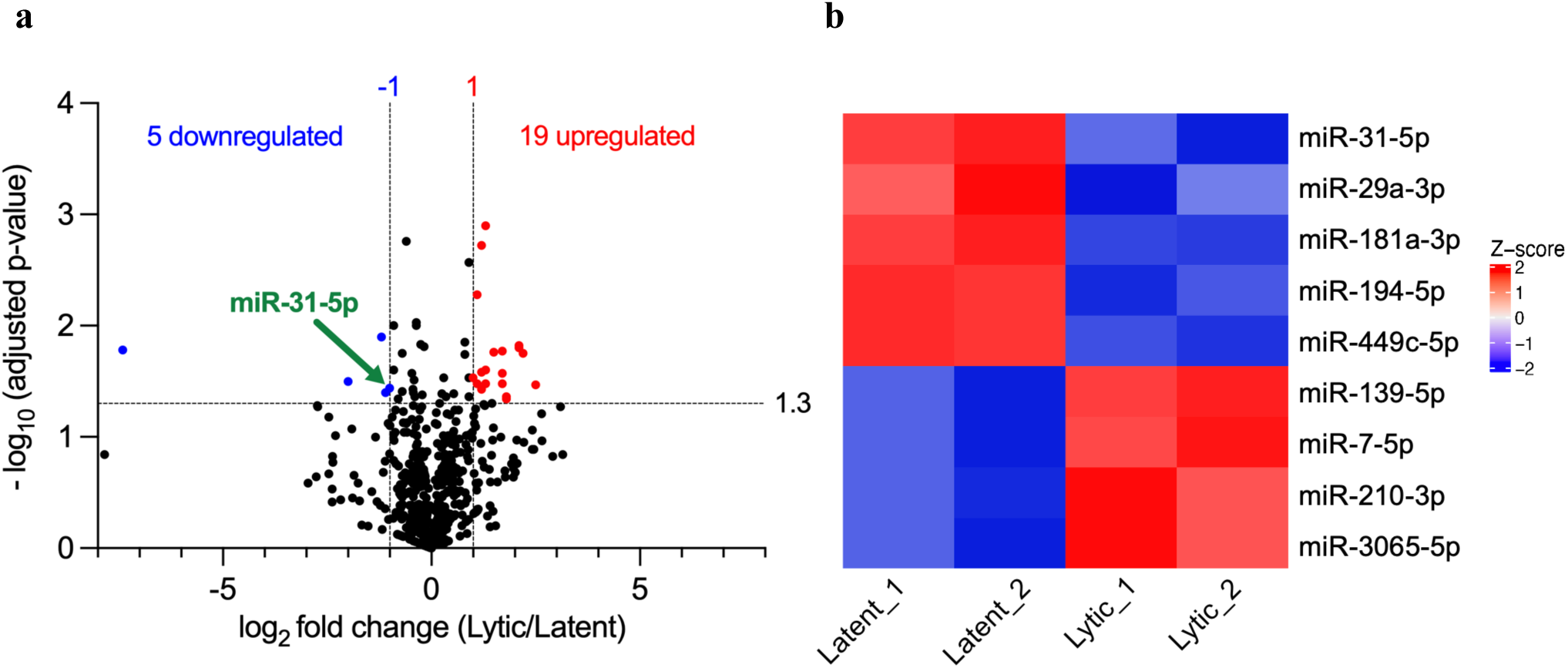
Characterization of cellular miRNA transcriptome during KSHV lytic reactivation by small-RNA sequencing. (**a**) Volcano plot representation of the small-RNA sequencing data depicting the fold change (log2 FC) of differentially expressed miRNAs in lytic iSLK.219 cells (Dox+) compared to latent iSLK.219 cells (Dox-) (two biological replicates) versus statistical significance of differential expression (-log10 [adjusted p-value]) 72 h post Dox-induced lytic reactivation. Up-regulated (red) and down-regulated (blue) genes have an adjusted p-value < 0.05 and log2 FC ≥ |1|. miR-31-5p (green) is labeled. (**b**) Heatmap depicting the expression of top down-regulated and up-regulated miRNAs (two biological replicates), as determined by small-RNA sequencing. Total RNA was isolated from latent iSLK.219 cells at 0 h or lytic iSLK.219 cells at 72 h post Dox-induced KSHV lytic reactivation and subjected to RNA sequencing analysis.

We also performed RNA sequencing analysis of RNA isolated from iSLK.219 in the latent phase or at 72 h post Dox-induced reactivation and analyzed global viral and cellular transcriptome changes between latent and lytic KSHV replication cycles. As expected, expression of viral lytic genes spanning all three kinetic classes, including IE, E, and L genes, was robustly induced during KSHV lytic replication, demonstrating successful induction of the full lytic cascade of viral gene expression (Supplementary Fig. 3a). In addition, gene ontology (GO) analysis demonstrated that cellular genes involved in cell cycle, DNA replication, and nuclear division, were significantly enriched among upregulated genes in reactivated lytic iSLK.219 cells, consistent with lytic infection activating mitogenic signaling to support viral DNA replication^33^ (Supplementary Fig. 3b). Moreover, expression of key viral lytic genes with strong mitogenic activities, such as the vIL-6 (K2), K1, and the KSHV-encoded chemokines, vCCL1 (K6), vCCL2 (K4) and vCCL3 (K4.1)^33^, were, as expected, markedly induced during lytic replication (Supplementary Fig. 3c). These results confirm that the gene expression profile of reactivated lytic iSLK.219 cells exhibit some key hallmark features of KSHV lytic reactivation.

### miR-31 restricts KSHV lytic gene expression and virion production during lytic reactivation

To examine the effect of miR-31 on KSHV lytic reactivation, we transfected iSLK.219 cells with miR-31 or negative control (NC) miRNA mimic and compared Dox-induced lytic reactivation between the two conditions. Subsequent induction of lytic reactivation led to a robust increase in lytically reactivated (RFP-positive) cells in the control sample, whereas miR-31-transfected cells showed a significant decrease in RFP-positive cells at 72 h post Dox-induced reactivation, indicating that overexpression of miR-31 repressed KSHV lytic reactivation (Supplementary Fig. 4a, b). Lytic reactivated (RFP-positive) cells were quantified using image cytometry (Supplementary Fig. 4b). To determine whether the observed effects of miR-31 on KSHV lytic reactivation is the result of its ability to restrict lytic gene transcription, we quantified the RNA levels of multiple KSHV lytic genes, spanning all three kinetic classes (IE, E, and L lytic genes). RT-qPCR analysis revealed that miR-31 transfection led to a significant reduction in the RNA levels of all KSHV lytic genes examined, including *ORF50* and *PAN*, compared to control cells transfected with NC mimic, indicating that miR-31-transfected cells have an impaired ability to induce transcription of these lytic genes (Fig. 2a). Consistent with these results, western blot analysis demonstrated that expression of ORF45 (IE gene), ORF50 (IE gene), K8 alpha (E gene), and ORF26 (L gene) viral proteins was significantly reduced in miR-31-transfected cells compared to control cells at 72 h post lytic reactivation (Fig. 2b).

**Figure 2.**
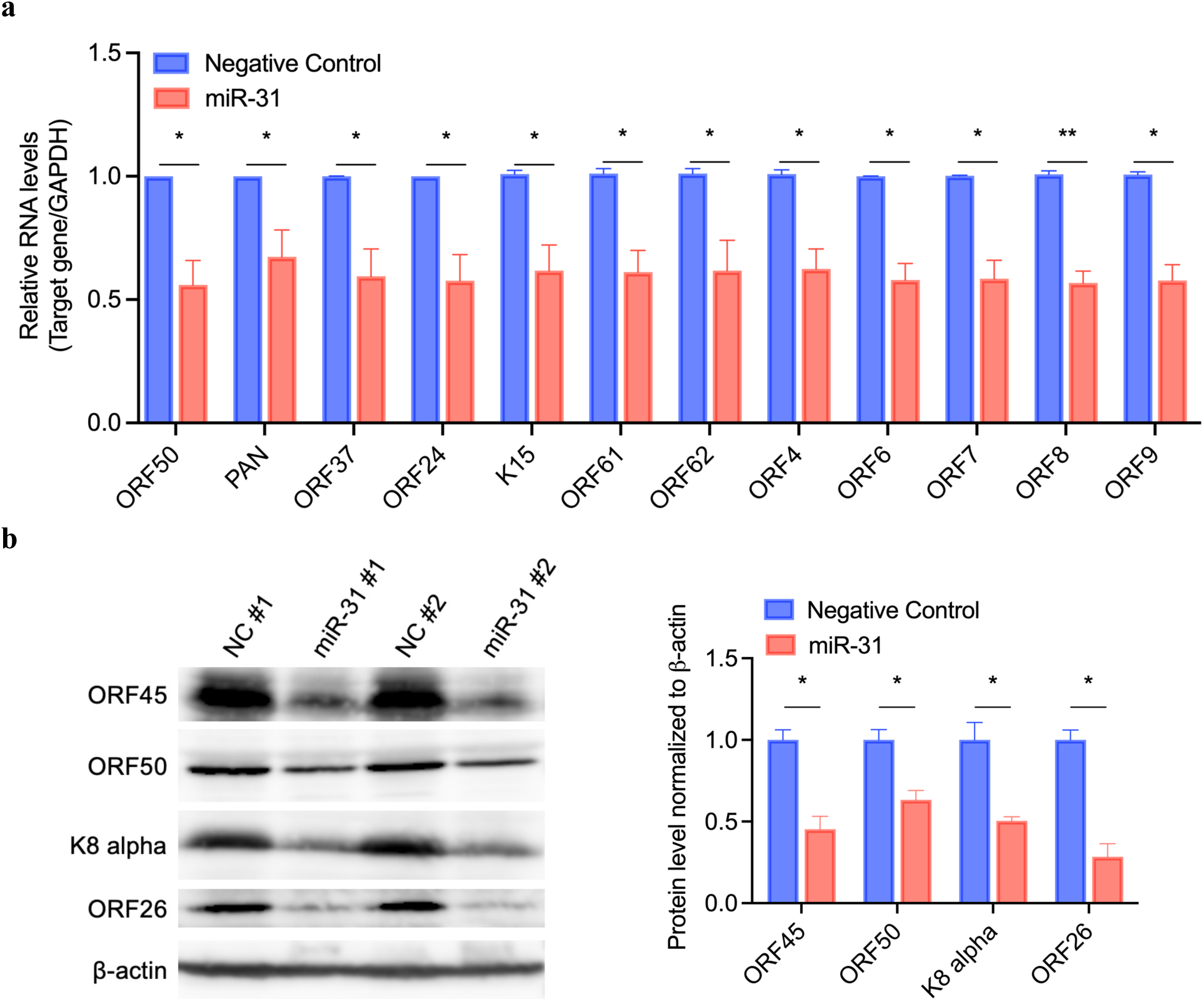
miR-31 restricts KSHV lytic gene transcription and viral protein expression during KSHV reactivation. (**a**) iSLK.219 cells were transfected with miR-31 or negative control (NC) mimic followed by treatment with Dox (0.5 µg/mL) for 72 h to induce KSHV lytic reactivation. Total RNA was isolated at 72 h post-Dox treatment and the expression levels of the indicated KSHV genes were quantified by RT-qPCR. *GAPDH* was used as an endogenous control. Three biological replicates are presented. (**b**) Whole cell lysates were collected from iSLK.219 cells transfected with miR-31 or NC mimic at 72 h post Dox-induced KSHV lytic reactivation and analyzed by western blot probing KSHV lytic proteins ORF45, ORF50, K8 alpha, and ORF26. Representative western blots and densitometric analysis of ORF45, ORF50, K8 alpha, and ORF26. β-actin was used as a loading control. Two biological replicates are presented. Data represent mean ± SEM and *p*-values were determined by two-tailed Student’s *t*-test. \**p* < 0.05, \*\**p* < 0.01, \*\*\**p* < 0.001, \*\*\*\**p* < 0.0001, and ns = not significant.

Next, to examine the impact of miR-31 on extracellular infectious virion production, iSLK.219 cells transfected with either NC or miR-31 mimic were subjected to Dox treatment to initiate KSHV lytic induction and the virus-containing supernatants collected at 96 h post-induction were used to infect naive HEK293T cells. Consistent with the decrease in lytic gene expression, we observed a significant reduction in rKSHV.219-infected (GFP-positive) HEK293T cells when the virus-containing media from miR-31-transfected cells was used for infection (Fig. 3a, b). rKSHV.219-infected (GFP-positive) HEK293T cells were quantified using image cytometry (Fig. 3b). Furthermore, consistent with these results, quantification of viral genomes in the culture supernatant by qPCR confirmed a significant decrease in KSHV genome copies in the media from cells treated with miR-31 compared to those treated with NC mimic, indicating a significant reduction in infectious virion production during KSHV lytic reactivation upon miR-31 overexpression (Fig. 3c).

**Figure 3.**
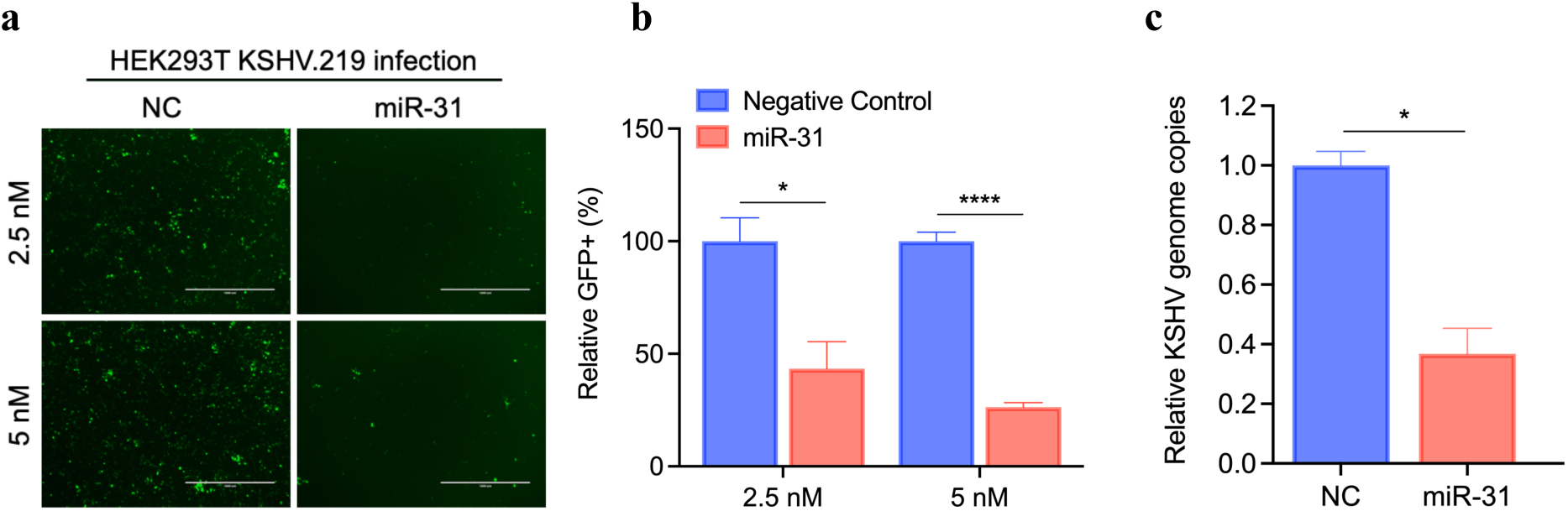
miR-31 impairs the production of infectious progeny virions during KSHV lytic replication. (**a**) iSLK.219 cells were transfected with miR-31 or negative control (NC) mimic followed by treatment with Dox (0.5 µg/mL) for 96 h to induce KSHV lytic reactivation. Virus-containing supernatants were collected at 96 h post-Dox treatment and used to infect naive HEK293T cells. Representative rKSHV.219-infected (GFP-positive) HEK293T cells were imaged at 48 h post infection (Scale bars, 1000 µm). (**b**) Quantification of rKSHV.219-infected (GFP-positive) HEK293T cells in (a) by image cytometry. Three biological replicates are presented. (**c**) Virus-containing supernatants of iSLK.219 cells transfected with miR-31 or NC mimic were collected for DNA extraction at 96 h post Dox-induced KSHV lytic reactivation and KSHV genomes in the supernatants were quantified by RT-qPCR. Two biological replicates are presented. Data represent mean ± SEM and *p*-values were determined by two-tailed Student’s *t*-test. \**p* < 0.05, \*\**p* < 0.01, \*\*\**p* < 0.001, \*\*\*\**p* < 0.0001, and ns = not significant.

Of note, the primers for RT-qPCR used to quantify *ORF50* levels are specific to the virus-encoded RTA and do not detect the exogenous RTA induced by Dox. Since RFP expression in the iSLK.219 cell line indicates lytic reactivation mediated by the Dox-induced expression of the RTA transgene, we further examined whether miR-31 transfection affected the expression of the RTA transgene induced by Dox. To test this, we transfected miR-31 or NC mimic into KSHV-negative iSLK cells, which lacks the KSHV genome but harbors the Dox-inducible RTA transgene, and quantified RTA transgene expression 72 h post-Dox treatment. Importantly, miR-31 transfection did not affect the expression of the Dox-induced RTA transgene in iSLK cells, indicating that the effect of miR-31 overexpression on KSHV lytic reactivation is independent of the RTA transgene (Supplementary Fig. 4c). Taken together, these findings demonstrate that ectopic expression of miR-31 greatly reduces the ability of KSHV to initiate and progress through the early lytic gene cascade, which ultimately results in a significant reduction in the expression of all viral lytic genes including late structural KSHV proteins necessary for viral particle formation, at both the transcriptional and translational level, culminating in a dramatic reduction in the production of new infectious virions.

### miR-31 directly targets the RNA-binding protein KHDRBS3 to repress KSHV lytic reactivation

To identify the cellular targets of miR-31 that may mediate its effects on KSHV reactivation and lytic replication, we integrated small-RNA sequencing analysis and the matched RNA sequencing (RNA-seq) data with the miRNA target prediction algorithm TargetScan^34^. We identified 135 miR-31 predicted targets with conserved and poorly conserved sites that are up-regulated (log2 FC ≥ 1) in iSLK.219 cells after Dox-induced lytic reactivation (72 h) compared to cells in the latent phase without Dox (0 h), indicating that miR-31 expression negatively correlated with these putative miR-31 targets (Supplementary Table 4). It is well known that the KSHV E gene product SOX (shutoff and exonuclease), encoded by ORF37, promotes shutoff of host gene expression during lytic KSHV infection by inducing global degradation of cellular transcripts (∼95%), with only ∼2% of host mRNAs strongly up-regulated^35–37^. Another study further revealed that lytic KSHV infection significantly down-regulates ∼41% of cellular proteins while ∼15% of host proteins are significantly up-regulated by lytic KSHV^38^. Notably, these up-regulated host proteins that escape SOX-mediated host shutoff (“escapees”) were enriched in Gene Ontology (GO) terms related to host cell machineries required for RNA processing, mRNA translation, and protein folding, emphasizing the significance of the host factors during KSHV lytic reactivation^38^. Ten of the 135 TargetScan predicted miR-31 targets mentioned above were escapees of SOX-directed host shutoff, and thus were selected for further analysis. These escapees included KHDRBS3, a key regulator of RNA processing, and nicotinamide N-methyltransferase (NNMT), a major methyltransferase that regulates cellular methylation potential. To further identify functional targets that are critically involved in KSHV lytic reactivation, we cross-referenced the 135 predicted miR-31 targets with a core set of host genes that are rapidly induced by lytic KSHV^39^. Importantly, these host genes are likely to play a pivotal role in driving the KSHV lytic replication cycle^39^. Cross-referencing revealed 11 putative miR-31 targets which are up-regulated during KSHV lytic reactivation, including SLC51A, a major transporter responsible for bile acid secretion. To determine whether miR-31 can target these cellular factors, we examined their mRNA levels upon miR-31 transfection. We observed a significant decrease in the mRNA levels of *KHDRBS3*, *SLC51A*, and *NNMT* in miR-31-transfected iSLK.219 cells compared to control cells 72 h post-Dox treatment, indicating that miR-31 represses the expression of these host genes (Fig. 4a). In contrast, the mRNA levels of other predicted target genes were not affected upon miR-31 transfection, indicating that these genes probably are not targets of miR-31 in this context (Fig. 4a). The TargetScan tool revealed that *KHDRBS3, SLC51A,* and *NNMT* mRNAs bear miR-31 seed-matched sites in their 3’ UTRs (Fig. 4b-d). To investigate whether miR-31 directly regulates these predicted target sites, we cloned the 3’ UTR of each gene into a dual-luciferase reporter and performed 3’ UTR assays. We found that the 3’ UTR luciferase activity of KHDRBS3 and SLC51A was significantly suppressed upon transfection of miR-31, indicating that miR-31 interacts with the 3’ UTR of *KHDRBS3* and *SLC51A* mRNAs to down-regulate their expression (Fig. 4b, c). Moreover, mutating the putative binding sites within the 3’ UTRs of *KHDRBS3* and *SLC51A* significantly reduced this suppressive effect (Fig. 4b, c), demonstrating that miR-31 directly represses KHDRBS3 and SLC51A expression via binding to the predicted miR-31 target sites. In contrast, miR-31 transfection had no effect on the luciferase activity of reporter plasmids harboring the 3’ UTR of *NNMT*, indicating that this host gene is not a direct target of miR-31 here (Fig. 4d).

**Figure 4.**
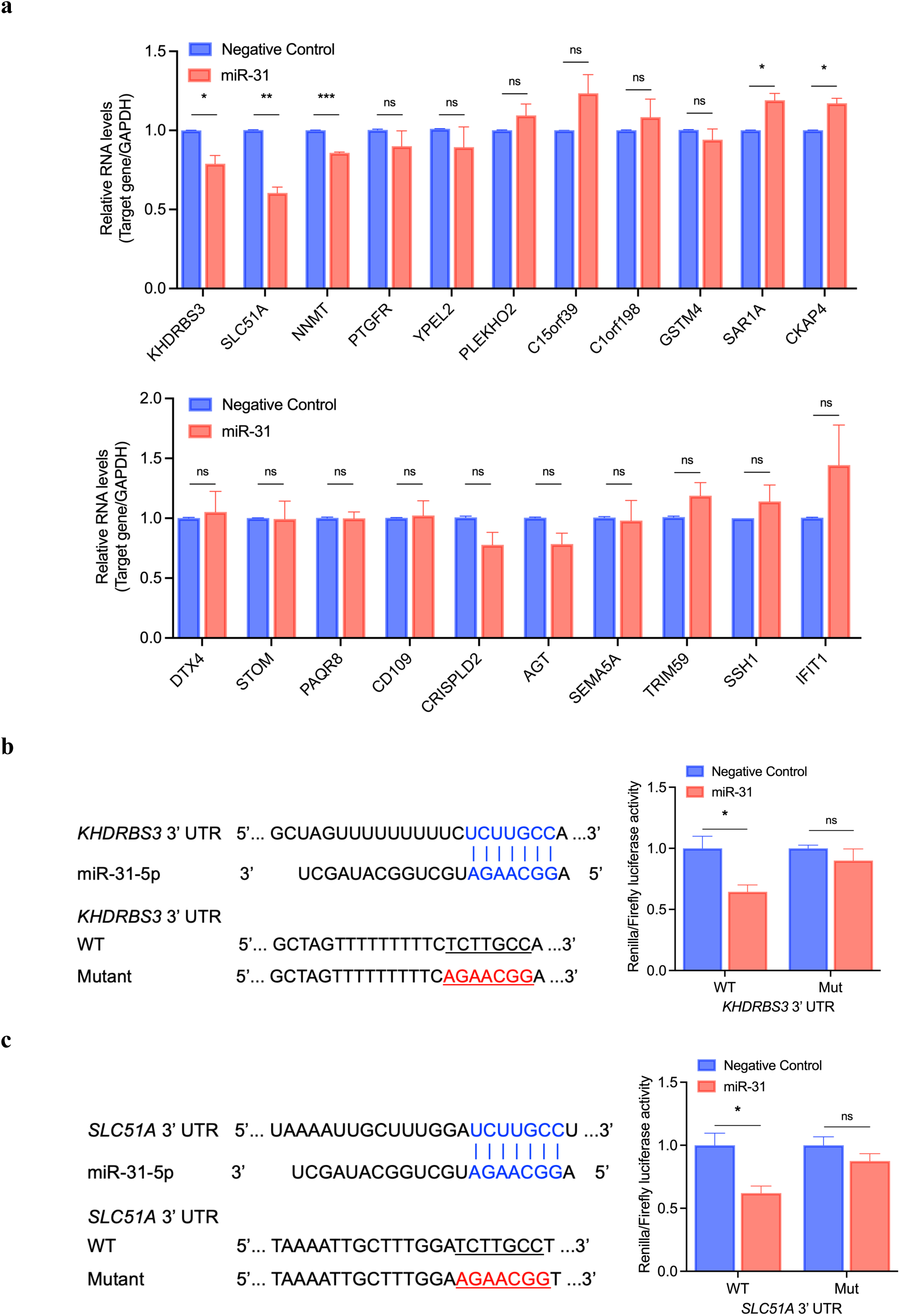
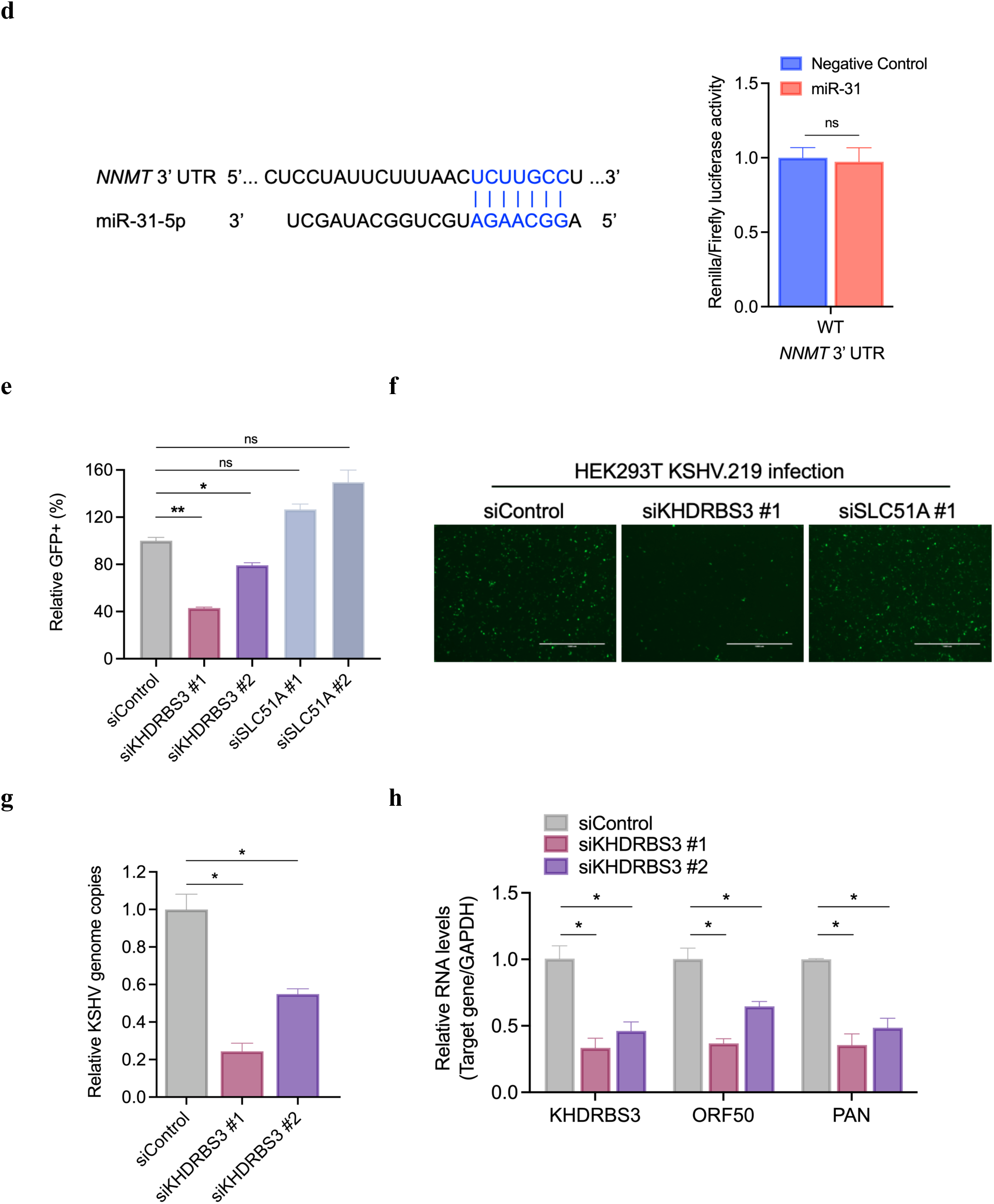
miR-31 directly targets KHDRBS3 to restrict the KSHV lytic gene cascade and infectious virion production during KSHV lytic reactivation. (**a**) iSLK.219 cells were transfected with miR-31 or negative control (NC) mimic followed by treatment with Dox (0.5 µg/mL) for 72 h to induce KSHV lytic reactivation. Total RNA was isolated at 72 h post-Dox treatment and the expression levels of the indicated cellular genes was quantified by RT-qPCR. *GAPDH* was used as an endogenous control. Three biological replicates are presented. (**b and c**) Left, schematic of the TargetScan predicted miR-31 8mer site or 7mer-m8 site in the 3’ UTR of *KHDRBS3* or *SLC51A* mRNA, respectively. The predicted pairing between the miR-31 seed and the wild-type (WT) 3’ UTR of *KHDRBS3* or *SLC51A* is shown in blue. Red letters indicate mutations introduced within the WT 3’ UTR to generate the mutant (Mut) 3’ UTR. Right, dual-luciferase reporter assay in iSLK.219 cells transfected with miR-31 or NC mimic. Firefly and Renilla luciferase activities were measured 72 h after cotransfection with miR-31 or NC mimic and the dual-luciferase reporter construct psiCHECK-2 encoding the WT or Mut 3’ UTRs of *KHDRBS3* or *SLC51A*. Two biological replicates are presented. (**d**) Left, schematic of the TargetScan predicted miR-31 7mer-m8 site in the 3’ UTR of *NNMT* mRNA. The predicted pairing between the miR-31 seed and the WT *NNMT* 3’ UTR is shown in blue. Right, dual-luciferase reporter assay in iSLK.219 cells transfected with miR-31 or NC mimic. Firefly and Renilla luciferase activities were measured 72 h after cotransfection with miR-31 or NC mimic and the dual-luciferase reporter construct psiCHECK-2 encoding the WT or Mut *NNMT* 3’ UTR. Two biological replicates are presented. (**e**) iSLK.219 cells were transfected with control siRNA (siControl) or two different siRNAs targeting KHDRBS3 (siKHDRBS3 #1 and #2) or SLC51A (siSLC51A #1 and #2) followed by treatment with Dox (0.5 µg/mL) for 96 h to induce KSHV lytic reactivation. Virus-containing supernatants were collected at 96 h post-Dox treatment and used to infect naive HEK293T cells. Quantification of rKSHV.219-infected (GFP-positive) HEK293T cells by image cytometry. Two biological replicates are presented. (**f**) Representative rKSHV.219-infected (GFP-positive) HEK293T cells in (**e**) were imaged at 48 h post infection (Scale bars, 1000 µm). (**g**) Virus-containing supernatants of iSLK.219 cells transfected with siControl, siKHDRBS3 #1 or siKHDRBS3 #2 were collected for DNA extraction at 96 h post Dox-induced KSHV lytic reactivation, and KSHV genomes in the supernatants were quantified by RT-qPCR. Two biological replicates are presented. (**h**) Total RNA was isolated from iSLK.219 cells transfected with siControl, siKHDRBS3 #1 or siKHDRBS3 #2 at 72 h post Dox-induced KSHV lytic reactivation and the expression levels of key KSHV lytic genes *ORF50* and *PAN* were analyzed by RT-qPCR. Knockdown efficiency of KHDRBS3 was also verified by RT-qPCR. *GAPDH* was used as an endogenous control. Two biological replicates are presented. Data represent mean ± SEM and *p*-values were determined by two-tailed Student’s *t*-test. \**p* < 0.05, \*\**p* < 0.01, \*\*\**p* < 0.001, \*\*\*\**p* < 0.0001, and ns = not significant.

Furthermore, we investigated viral targets through which miR-31 may impact reactivation and lytic replication of KSHV. To identify potential KSHV RNA targets of miR-31, sequences corresponding to the annotated 3’ UTRs of KSHV transcripts were extracted and processed by the miRanda algorithm^40^. The vast majority of KSHV transcripts utilizes the same polyadenylation site, giving rise to partially overlapping 3’ UTRs with shared regulatory elements and activities^41^. We identified putative seed-matched target sites for miR-31 in the overlapping 3’ UTR region of two KSHV lytic gene clusters: the *ORFs 4, 6, 7, 8, 9* and *10* gene cluster which encodes several lytic proteins involved in viral replication and virion egress; the *ORF61* (ribonucleotide reductase large subunit) and *ORF62* (viral capsid protein) cluster. In addition, we identified putative target sites in the 3’ UTRs of *ORF24* (viral TATA box-binding protein) and *K15* (signal transducing membrane protein). Intriguingly, we found that *PAN* RNA contains three putative binding sites for miR-31. Thus, it is possible that miR-31 binds to *PAN* RNA to down-regulate its expression. The full list of computationally predicted viral targets is presented in Supplementary Table 5. To determine if these viral genes are bona fide miR-31 targets, we attempted to clone the entire *PAN* RNA or the 3’ UTR of each gene downstream of a Renilla luciferase gene within a dual-luciferase reporter construct and performed 3’ UTR assays to assess whether miR-31 targets these predicted binding sites. We managed to successfully clone the 3’ UTRs of *ORF9, ORF24, ORF61* and the entire *PAN* RNA. Ectopic expression of miR-31 had no effect on the luciferase activity of reporter plasmids harboring the 3’ UTRs of *ORF9, ORF24, ORF61* or the *PAN* RNA, indicating that these viral transcripts are likely not direct targets of miR-31 here (data not shown). Therefore, these results suggest that miR-31 indirectly represses these viral genes, at least in part, through down-regulation of KSHV latent-lytic transactivator RTA expression, thereby restricting RTA-mediated transcription and lytic gene expression required for efficient lytic KSHV replication.

Next, to investigate the role of KHDRBS3 and SLC51A during KSHV lytic reactivation and to determine whether depleting KHDRBS3 and SLC51A results in a similar phenotype observed with overexpression of miR-31, we employed a short interfering RNA (siRNA)-based loss-of-function approach whereby iSLK.219 cells were transfected with a scrambled siRNA control or two different siRNAs targeting *KHDRBS3* or *SLC51A*. Notably, depletion of *KHDRBS3* resulted in a significant reduction in infectious virion production, as assessed by rKSHV.219-infected (GFP-positive) HEK293T cells at 48 h post infection (Fig. 4e, f), similar to that observed with the overexpression of miR-31 (Fig. 3a, b), indicating that KHDRBS3 is essential for efficient KSHV lytic replication. In contrast, depletion of SLC51A did not impact infectious virion production compared to scrambled control cells. rKSHV.219-infected (GFP-positive) HEK293T cells were quantified using image cytometry (Fig. 4e). These results demonstrate that KHDRBS3, but not SLC51A, is essential for the effective production of new infectious virions during KSHV lytic reactivation. Consistently, subsequent quantification of viral genomes in the collected virus-containing supernatants by qPCR confirmed a significant decrease in KSHV genome copies in the media from KHDRBS3-depleted cells compared to scrambled control cells (Fig. 4g). We next determined the effect of siRNA-mediated KHDRBS3 knockdown on Dox-induced lytic gene expression in iSLK.219 cells. As expected, RT-qPCR analysis showed a significant reduction in the RNA levels of key IE lytic genes critically involved in early KSHV lytic replication, *ORF50* and *PAN*, in KHDRBS3-depleted cells compared to scrambled control cells at 72 h post Dox-induced lytic reactivation, demonstrating that depletion of KHDRBS3 has a dramatic impact on the early stages of the KSHV lytic gene cascade (Fig. 4h), which would have a direct impact on late lytic cycle progression, viral DNA replication, and viral particle formation (Fig. 4g). Taken together, these results collectively show that KHDRBS3 is required for the optimal expression of viral lytic transcripts and essential for the efficient production of new infectious virions during KSHV lytic reactivation. Our findings therefore demonstrate that miR-31 restricts KSHV lytic reactivation by directly repressing KHDRBS3.

### miR-31-mediated KHDRBS3 repression restricts viral lytic gene transcription, resulting in suppression of KSHV lytic reactivation

KHDRBS3 (KH domain-containing, RNA-binding, signal transduction-associated 3) is a member of the STAR (signal transduction and activation of RNA) family of proteins, which are evolutionarily conserved RNA-binding proteins that regulate gene expression at all stages of RNA metabolism^42–44^. Since KHDRBS3 is known to control alternative splicing through differential selection of splice sites in its target transcripts, influencing the diversity of protein isoforms produced within cells^17–21^, we sought to determine whether KHDRBS3 plays a role in regulating alternative splicing of viral lytic transcripts during KSHV reactivation. We focused on alternative splicing of ORF50 (IE gene), ORF57 (E gene), and K8.1 (L gene) as these are the only lytic genes of which alternatively spliced isoforms have been well-described^45–47^. Analysis of alternatively spliced events by RT-qPCR revealed a significant decrease in the expression of all examined alternatively spliced transcripts of these viral lytic genes in miR-31-transfected cells compared to those transfected with NC mimic at 72 h post Dox-induced reactivation, indicating that miR-31-directed KHDRBS3 repression does not result in notable alterations in splicing patterns of viral lytic genes *ORF50*, *ORF57*, and *K8.1* (Fig. 5a-c and Supplementary Fig. 5a-c); in other words, distinct patterns of alternative splicing are not associated with depletion of KHDRBS3 during KSHV lytic replication. To corroborate these results, we also employed siRNAs to down-regulate KHDRBS3 expression levels in iSLK.219 cells. Accordingly, knockdown of KHDRBS3 led to a significant reduction in the expression levels of all the alternatively spliced transcripts of these lytic genes compared to scrambled control cells, closely mirroring the phenotype observed with the direct repression of KHDRBS3 via miR-31 (Fig. 5d-f). Thus, it is probable that the knockdown of KHDRBS3 resulted in decreased overall abundance of all the spliced mRNA isoforms of *ORF50*, *ORF57*, and *K8.1* genes through its impact on transcription, rather than by influencing specific alternative splicing patterns. These results suggest that KHDRBS3 does not regulate alternative splicing of this subset of KSHV transcripts. Nonetheless, these findings further demonstrate that KHDRBS3 is essential for effective KSHV lytic replication and necessary for the expression of IE, E, and L lytic viral transcripts during the lytic replication phase of KSHV.

**Figure 5.**
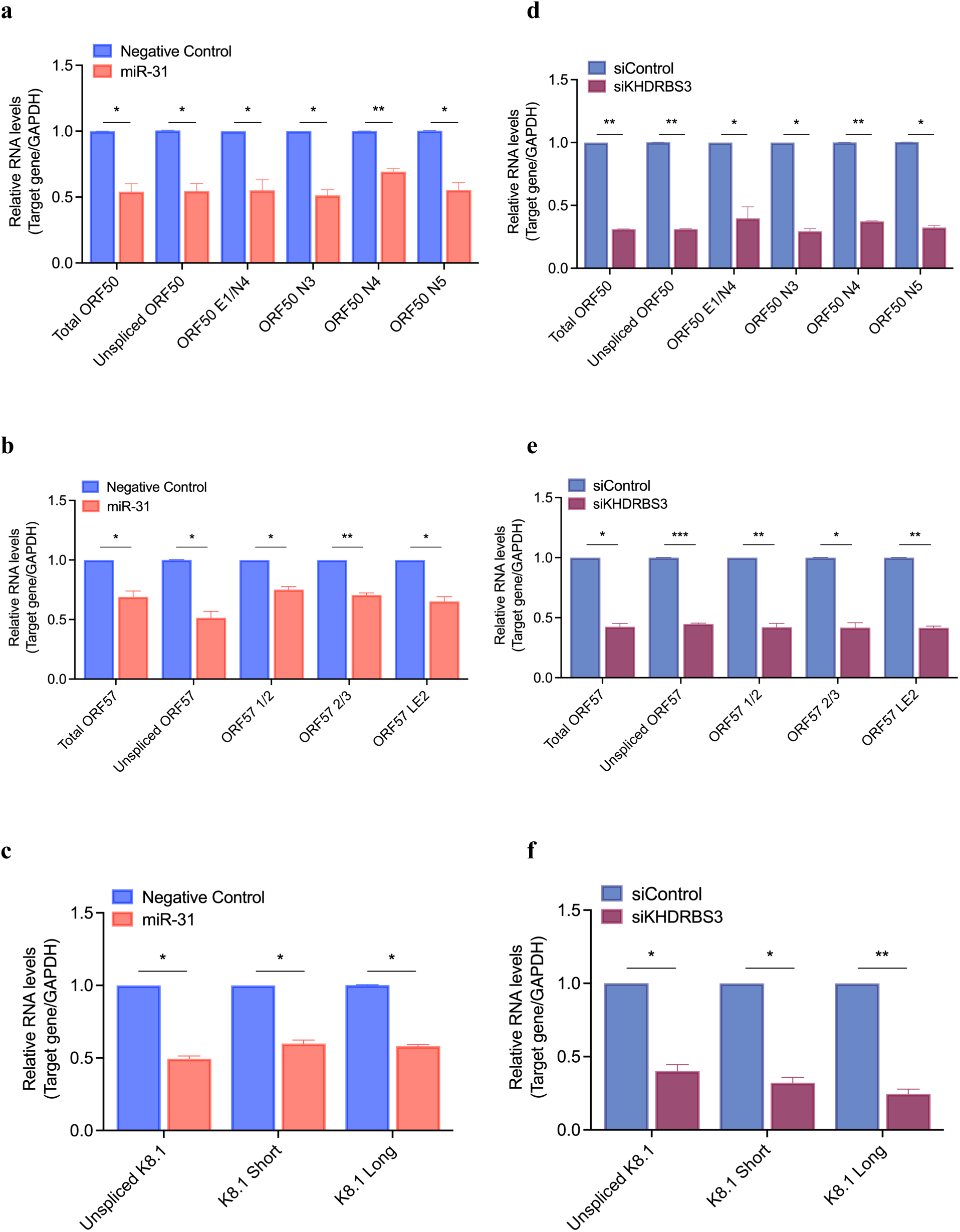
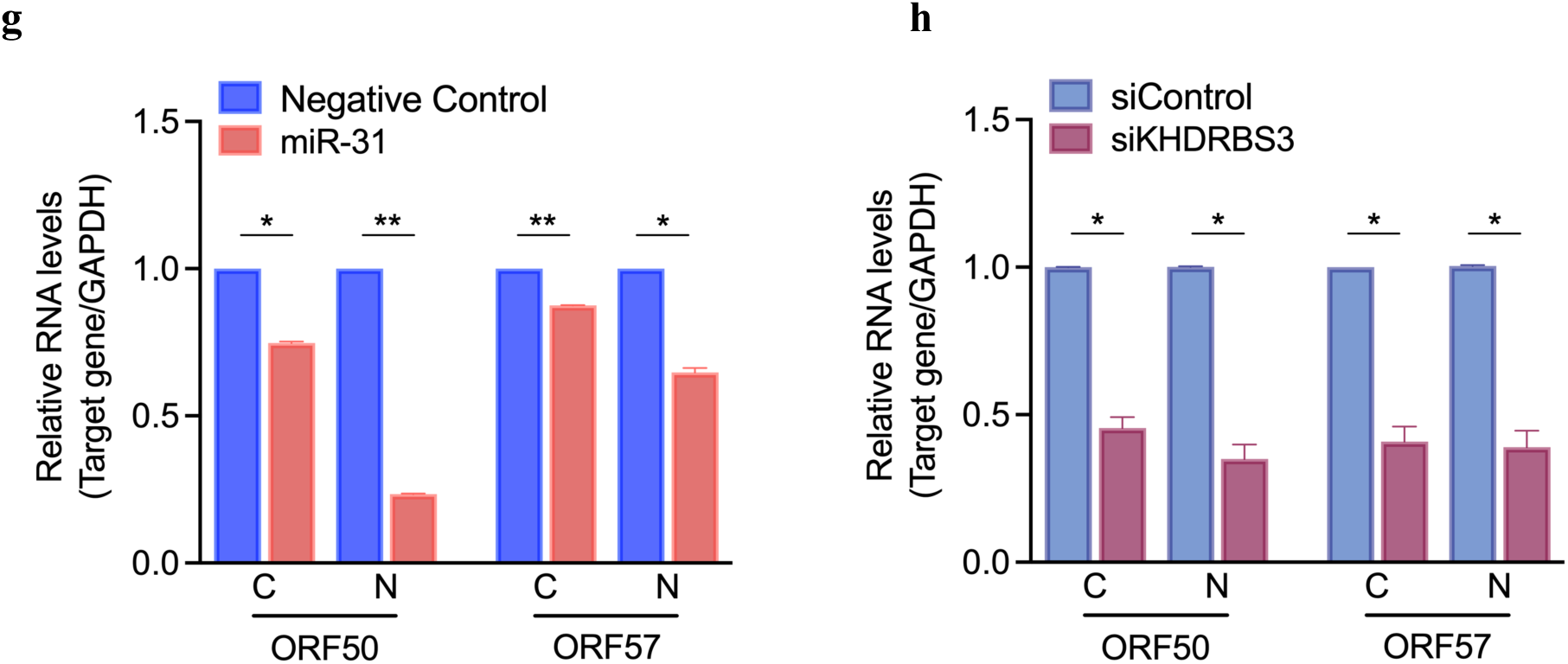
miR-31-mediated viral RNA-processing changes during KSHV lytic reactivation is replicated by KHDRBS3 depletion. (**a-c**) Analysis of alternative splicing patterns of *ORF50* (a), *ORF57* (b), and *K8.1* (c) in iSLK.219 cells transfected with miR-31 or negative control (NC) mimic. Total RNA was isolated from iSLK.219 cells transfected with miR-31 or NC mimic at 72 h post Dox-induced KSHV lytic reactivation and the expression levels of the indicated alternatively spliced transcripts of the immediate-early (IE) lytic KSHV gene *ORF50* (a), early (E) lytic KSHV gene *ORF57* (b), and late (L) lytic KSHV gene *K8.1* (c) were analyzed by RT-qPCR. *GAPDH* was used as an endogenous control. Three biological replicates (a, b) and two biological replicates (c). **(d-f**) Analysis of alternative splicing patterns of *ORF50* (d), *ORF57* (e), and *K8.1* (f) in iSLK.219 cells transfected with control siRNA (siControl) or siRNA targeting KHDRBS3 (siKHDRBS3). Total RNA was isolated from iSLK.219 cells transfected with siControl or siKHDRBS3 at 72 h post Dox-induced KSHV lytic reactivation and the expression levels of the indicated alternatively spliced transcripts of IE *ORF50* (d), E *ORF57* (e), and L *K8.1* (f) were analyzed by RT-qPCR. *GAPDH* was used as an endogenous control. Three biological replicates (e, f) and two biological replicates (d). (**g and h**) Subcellular fractionation assay in iSLK.219 cells transfected with miR-31 or NC mimic (g), and iSLK.219 cells transfected with siControl or siKHDRBS3 (h). iSLK.219 cells transfected with the indicated miRNA mimics or siRNAs were treated with Dox for 72 h to induce KSHV lytic reactivation and then subjected to subcellular fractionation into cytoplasmic and nuclear fractions. Total RNA was isolated from cytoplasmic (C) and nuclear (N) fractions at 72 h post-Dox treatment and the expression levels of KSHV lytic genes *ORF50* and *ORF57* were quantified by RT-qPCR. *GAPDH* was used as an endogenous control. Two biological replicates (g) and three biological replicates (h). Data represent mean ± SEM and *p*-values were determined by two-tailed Student’s *t*-test. \**p* < 0.05, \*\**p* < 0.01, \*\*\**p* < 0.001, \*\*\*\**p* < 0.0001, and ns = not significant.

KHDRBS3 has previously been shown to enhance HIV-1 Rev activation of Rev response element (RRE)-mediated gene expression and nuclear export of unspliced mRNA into the cytoplasm, suggesting that KHDRBS3 may play a crucial role in the export or stability of unspliced and partially spliced HIV-1 RNAs with retained introns^22^. Moreover, considering the multifaceted functions of the KH domain-containing RNA-binding proteins in regulating various aspects of RNA metabolism, from alternative splicing to mRNA trafficking, localization, and translation^48–50^, we conducted subcellular fractionation assays to investigate whether KHDRBS3 may control the nuclear export of KSHV lytic transcripts and therefore their subcellular localization (cytoplasm vs. nucleus distribution). RT-qPCR analysis showed a significant decrease in the mRNA levels of IE *ORF50* and E *ORF57* genes in both the nuclear and cytoplasmic fractions upon miR-31 transfection, at 72 h post Dox-induced reactivation (Fig. 5g). These results show that miR-31-mediated KHDRBS3 knockdown does not result in the nuclear accumulation of these viral transcripts and does not impact their overall subcellular localization, indicating that the export of *ORF50* and *ORF57* mRNAs from the nucleus into the cytoplasm is not dependent on KHDRBS3. In accordance with these results, knockdown of KHDRBS3 by siRNA did not impact export of *ORF50* and *ORF57* mRNAs and reduced overall expression of these viral transcripts in both the nuclear and cytoplasmic compartments, compared to cells transfected with control siRNA, further demonstrating that KHDRBS3 depletion results in the down-regulation of these KSHV transcripts in the nucleus (Fig. 5h), which ultimately leads to the breakdown of the KSHV lytic cascade, preventing the production of new infectious virions (Fig. 4g). This, taken together with the effect of KHDRBS3 knockdown and associated overall decrease in alternatively spliced events suggest that this RNA-binding protein acts upstream of RNA splicing and nuclear export to control the expression of these viral mRNAs in the nucleus, most likely by affecting their transcription.

## Discussion

Small regulatory RNAs, particularly miRNAs, are known to play important roles in host and virus interactions occurring at the post-transcriptional level during viral infections, and the alterations in these small noncoding RNAs could contribute to the pathogenesis of many RNA and DNA viruses^14,15^. However, the role of host miRNAs in the regulation of KSHV lytic reactivation remains largely unknown. In this study, we conducted comprehensive miRNA-mRNA transcriptomics and phenotypic analyses revealing that miR-31 is a key regulator of KSHV lytic reactivation capable of restricting the switch from latent to lytic replication and the formation of infectious progeny virions. We find that ectopic expression of miR-31 impairs KSHV lytic gene transcription and viral protein expression, ultimately resulting in a dramatic reduction in infectious virion production. Mechanistically, this effect is mediated by direct targeting of the host RNA-binding protein KHDRBS3 via miR-31, which is dependent on a highly conserved 8mer seed-matched site for miR-31 in the 3’ UTR of *KHDRBS3* mRNA, resulting in mRNA destabilization and translational repression. Consistently, the reactivation of lytic gene expression and the generation of viable viral particles are significantly suppressed upon depletion of KHDRBS3 by siRNA. miR-31-mediated KHDRBS3 repression also results in significantly reduced expression of critical viral early genes, including ORF50-encoded RTA transcripts and RTA protein expression which, as RTA is the master transcription factor responsible for initiating and driving the KSHV lytic replication cycle, compromises the overall KSHV lytic gene cascade. Consistent with these modes of action, we find that overexpression of miR-31 broadly restricts all kinetic classes of KSHV lytic genes at both the RNA and protein levels during lytic reactivation. Furthermore, KSHV virion production following lytic replication was, as expected, significantly impaired by miR-31 through suppression of viral RNA transcription and viral protein synthesis.

Because an individual miRNA can regulate multiple genes, it is possible that miR-31 has additional biological targets. For example, implicated in the regulation of diverse biological processes, miR-31 has been reported to play a role in immune function, angiogenesis, and cancer by targeting genes involved in apoptosis, cell differentiation, cell motility, and angiogenesis^51,52^. Nevertheless, our findings show that miR-31 regulates the expression of a lytic gene cascade that initiates KSHV lytic replication and consequently the production of new infectious virions by directly repressing KHDRBS3 expression, and that the miR-31/KHDRBS3 axis plays an important regulatory role during KSHV reactivation. The most compelling evidence supporting this comes from the observation that KHDRBS3 depletion by siRNA results in significantly decreased levels of KSHV lytic genes spanning all three kinetic classes (i.e. IE, E, and L lytic genes), and thus subsequently progeny virion production, which phenocopies overexpression of miR-31, demonstrating that these effects are predominantly due to the direct repression of KHDRBS3 via miR-31.

Our findings suggest that the RNA-binding protein KHDRBS3 (also known as Sam68-like mammalian protein 2 (SLM2)) serves as a proviral host factor for KSHV lytic reactivation and is essential for lytic gene expression and subsequent progeny virion production. RNA-binding proteins are a diverse class of proteins that have emerged as central players in gene regulation^48,49,53–57^. They control the maturation and fate of their target RNA substrates by regulating all aspects of RNA metabolism, including mRNA splicing, mRNA polyadenylation, mRNA stability, trafficking, and localization^48,49,53–57^. Although it was originally thought that these steps largely occurred post-transcriptionally, it is now clear that most of these steps can occur co-transcriptionally^58–61^. In addition, recent studies have unveiled widespread, unexpected roles for RNA-binding proteins in controlling gene transcription, which is now thought to be functionally coupled to many co-transcriptional RNA processing events^84^. KHDRBS3 is a member of the hnRNP K-homology (KH) and STAR family of RNA-binding proteins, composed of the KH domain that mediates RNA binding and protein-protein interaction and a nuclear localization sequence in the C-terminal tyrosine-rich region^42–44^. Emerging as a key player in splicing regulation, KHDRBS3 has also been implicated in downstream RNA processing events, such as RNA stability or export^17–23^.

Increasing evidence indicates that RNA-binding proteins are exploited or hijacked by many viruses to enhance viral gene expression in response to various external stimuli that trigger viral replication and propagation^62–67^. Indeed, KHDRBS3 has been reported to play an important role in the regulation of HIV-1 gene expression^22,23^. Specifically, KHDRBS3 synergizes with the HIV-1 Rev protein, an essential regulator of the HIV-1 mRNA expression that facilitates the nuclear export of unspliced and partially spliced viral mRNAs, in RRE-directed HIV-1 gene expression^22,23^. Moreover, KHDRBS3 was also shown to promote HIV-1 Rev-mediated nuclear export of unspliced mRNA into the cytoplasm, suggesting that this RNA-binding factor may play a critical role in modulating the export or stability of unspliced HIV-1 RNAs which serve either as genomic RNA or as mRNAs for essential structural proteins^22^. To ensure the integrity of gene expression, unspliced or incompletely spliced mRNAs containing introns are normally retained and degraded in the nucleus^68,69^. However, the HIV Rev protein and its cognate target sequence, RRE, an essential viral RNA element, drive the nuclear export of partially spliced transcripts as well as the unspliced mRNA, which are responsible for the synthesis of all viral proteins to achieve successful HIV-1 replication and viral particle formation^70,71^. Another member of the KH domain family, KHDRBS1 (also known as Sam68), the closest relative of KHDRBS3, has also been reported to functionally complement for and synergize with HIV-1 Rev in RRE-mediated viral gene expression, nuclear RNA export, and virus production, and is absolutely required for HIV-1 production^23,62,63,72–74^. It has been demonstrated that KHDRBS1 enhances 3’ end processing of unspliced viral RNAs to promote viral protein synthesis, suggesting that KHDRBS1-induced increase in HIV-1 protein expression is achieved through enhancement of HIV-1 RNA 3’ end formation, which likely results in increased translational efficiency^75^. Furthermore, KHDRBS1 is also able to enhance the activities of the Rev-like proteins of other complex retroviruses, namely Human T-cell leukemia virus (HTLV) and equine infectious anemic virus (EIA V)^67^. Intriguingly, it was also observed that herpes simplex virus 1 (HSV-1) RNA-binding protein Us11 can functionally substitute for Rev in mediating RRE-directed gene expression and facilitate Rev-dependent viral RNA export^76,77^.

KSHV lytic reactivation is a highly orchestrated process characterized by coordinately and temporally regulated changes in the expression of hundreds of lytic genes within distinct functional categories which are subject to extensive transcriptional and post-transcriptional regulation. Given the importance of RNA-binding proteins of KHDRBS family in gene regulation, we speculated that KHDRBS3 could be essential for the proper regulation of KSHV lytic gene expression during the viral lytic gene cascade. It is possible, for example, that KHDRBS3 functions as a key splicing factor that promotes alternative splicing of viral lytic pre-mRNAs. Previous studies have shown that alternative splicing contributes significantly to enhance viral transcriptome complexity and thus viral proteome diversity during KSHV lytic reactivation^78–82^. Nearly one-third of all KSHV genes are subject to alternative splicing during lytic replication^83^, including the main KSHV latent-lytic switch protein RTA encoded by ORF50^45^. A recent study reported 372 novel unannotated introns in iSLK.219 cells, while only 27 introns were previously described, revealing a highly complex splicing landscape of the KSHV transcriptome that is still not fully understood^82^. This study refined the existing annotation of KSHV transcriptome by identifying many previously unknown pre-mRNA splicing events^82^.

Considering that the master switch of KSHV lytic reactivation RTA encoded by *ORF50* undergoes highly complex alternative splicing and expresses multiple spliced transcripts^45^, we hypothesized that RTA-induced KSHV lytic cycle activation could be controlled by KHDRBS3-mediated splicing of *RTA* pre-mRNA. To test this hypothesis, we analyzed alternative splicing of the IE lytic gene *ORF50*, E lytic gene *ORF57*, and L lytic gene *K8.1* during KSHV reactivation upon miR-31-directed KHDRBS3 knockdown and found a significant decrease in the expression of all the alternatively spliced transcripts examined in this study, closely mirroring the effect of depleting KHDRBS3 by siRNA, suggesting that KHDRBS3 does not mediate alternative splicing events occurring within these viral lytic genes and thus does not contribute to establishing distinct patterns of their splicing. Furthermore, subcellular fractionation assays showed significantly reduced expression of *ORF50* and *ORF57* mRNAs in both the nuclear and cytoplasmic compartments upon KHDRBS3 depletion. Thus, knockdown of KHDRBS3 did not result in the nuclear accumulation of these viral transcripts but rather led to their down-regulation in the nucleus accompanied by a significant decrease in the cytoplasm, suggesting that KHDRBS3 plays a critical role in transcription of the IE *ORF50* and E *ORF57* viral genes during KSHV reactivation. Collectively, these findings demonstrate that this RNA-binding factor acts upstream of splicing and export, influencing the fate of these viral transcripts in the nucleus, most likely through regulating their transcription - similar to the proposed role of nuclear RNA-binding proteins in transcription control^84^. Alternatively, KHDRBS3 may regulate the expression of a key transcription factor or a chromatin remodeler that is critically involved in selective transcriptional activation of IE viral genes via modulation of local chromatin configuration as KSHV transitions to the lytic phase from its latency. Although our results suggest that KHDRBS3 does not control alternative splicing and nuclear export of *ORF50* and *ORF57* mRNAs, it could be involved in mRNA export or splicing regulation of a specific subset of other viral transcripts, ensuring their proper splicing and selective transit through nuclear pores to the cytoplasm to facilitate rapid induction of viral protein translation during KSHV reactivation.

KSHV SOX-induced RNA decay is a seemingly self-destructive strategy where viral RNAs must successfully navigate around the cellular RNA decay machinery to regulate and maintain the stability of viral transcripts and promote a highly productive lytic infection. Therefore, the lytic reactivation-induced up-regulation of KHDRBS3 might be a compensatory mechanism to adapt to widespread RNA decay triggered by lytic KSHV, thereby maintaining the integrity of the KSHV transcriptome to ensure efficient KSHV lytic replication. In addition, rapid induction of protein translation via KHDRBS3-mediated processing of mRNAs encoding key proteins involved in lytic replication can contribute to the regulation and complexity of lytic gene expression. Intriguingly, we observed elevated KHDRBS3 levels in bulk endemic and epidemic KS^85^ (Supplementary Fig. 6), suggesting an important role for this RNA-binding protein in KS tumorigenesis, perhaps it has oncogenic functions through its engagement with specific oncogenes or tumor suppressor genes and critical targets implicated in tumor growth - consistent with the emerging role for RNA-biding proteins as cancer drivers^86–88^. Many RNA-binding proteins have been found up-regulated in multiple cancer types, including KHDRBS1^87–89^, the closest relative of KHDRBS3, which has been implicated in the oncogenesis and progression of several human cancers^86–88,90^. It is important to note, however, that given the immense phenotypic, temporal, and spatial heterogeneity of KS cells and cells composing the tumor immune microenvironment (TIME), it will be critical to leverage single-cell and spatial technologies to determine KHDRBS3 expression in malignant cells versus other cell types of the TIME to distinguish contributions from the neoplastic compartment versus non-malignant cells in the microenvironment. This approach will provide insights into the cell type- and context-dependent roles of KHDRBS3 in KS. Similarly, given the context dependency of miRNA-target interactions, it will be important to examine intercellular heterogeneity of miR-31 expression and targeting in complex KS tissues at the single-cell level to assess the impact of tumor heterogeneity and give insights into cell type- and cell status-specific miR-31 functions and miR-31-mediated target regulations obscured at the bulk level.

While overexpression of miR-31 resulted in repression of KSHV reactivation and lytic replication, we speculate that endogenous miR-31 may play an important proviral role in regulating the steady-state levels of KSHV lytic mRNAs by modulating KHDRBS3 expression, potentially serving as a molecular buffer against unwanted fluctuations in lytic gene transcription due to aberrant or inappropriate activation of the KSHV lytic replication cycle, which may elicit a damaging inflammatory response to the virus and ultimately lead to its immune elimination. Such ability to tightly control or fine-tune KSHV lytic reactivation, especially in the face of environmental and stochastic perturbations, would help maintain optimal levels of KSHV lytic transcripts and subsequent wave of lytic activity to ensure robust persistence, transmission, and pathogenicity of KSHV. Indeed, previous studies have shown that miRNAs help to confer robustness to various biological processes by buffering against unwanted variations in gene expression^91^. Moreover, because unwanted fluctuations in the KSHV lytic cycle activity can inadvertently trigger host antiviral immune responses and hasten viral clearance, such miR-31-mediated noise buffering would also minimize damaging consequences for the virus and the host. This is particularly critical for herpesviruses due to their long-term maintenance of latent infections, which are completely dependent on the cellular machinery of the infected host cell to successfully complete their life cycles^92^. During millions of years of co-evolution with its human host, KSHV has evolved a plethora of regulatory mechanisms to influence the levels of RTA-mediated transactivation and lytic reactivation, primarily through transcriptional mechanisms that affect RTA abundance and activity^93^. Thus, we speculate that miR-31 adds another layer of post-transcriptional control for the KSHV lytic switch that extends the regulation at other levels, allowing for precise control of the initial wave of lytic cascade and subsequent virion production, by regulating KHDRBS3-mediated viral gene expression, thereby modulating both the magnitude and duration of the KSHV lytic cycle activity and adjusting the levels of reactivation depending on the cellular context and the nature of the stimuli.

Given the versatile roles of RNA-binding proteins in all aspects of RNA metabolism^48,49,53–57,84^, it is plausible that KHDRBS3 has instrumental roles in orchestrating a myriad of RNA processing events, at transcriptional and/or post-transcriptional levels, crucial for remodeling the viral and host transcriptomes and shaping the kinetics of lytic gene induction necessary for optimal KSHV reactivation. For future studies, it will be of interest to characterize global genome-wide changes in RNA processing upon KHDRBS3 silencing to determine its effects on downstream RNA processing activities, such as splicing regulation, in the human and viral genomes, and to elucidate the mechanisms by which it regulates these dynamic alterations in viral and cellular RNA processing during KSHV lytic reactivation. It is possible that the expression of KSHV lytic genes could also be regulated co/post-transcriptionally by other RNA-binding proteins that form RNA-protein complexes of various compositions affecting other key aspects of RNA metabolism during the KSHV lytic gene cascade. Looking beyond KSHV, we further speculate that KHDRBS3 may serve more broadly as a mediator in the switch from latent to lytic replication and facilitate the lytic cascade for other herpesviruses, particularly alpha- and gamma-herpesviruses, which are known to induce global mRNA decay during lytic infection^94^. Thus, future investigation will illuminate how KHDRBS3 and other host RNA-binding proteins modulate the activity and fate of viral RNAs, which have to escape cellular RNA decay pathways to enable the optimal replication and production of infectious viruses. Looking forward, we anticipate that deeper explorations in this area will offer new insights into functional interactions between host RNA-binding proteins and viral RNAs during the viral life cycle and their critical role for efficient reactivation and virion production, which can inform development of innovative antiviral therapies targeting key pathogenic viral RNA-host protein interaction mechanisms.

## Materials and Methods

### Cell lines and cultures

iSLK.219 cells were maintained in DMEM (Gibco) supplemented with 10% FBS, 1% penicillin-streptomycin (Gibco), 1 µg/mL puromycin (Gibco), 250 µg/mL G418 (Gibco), and 1200 µg/mL hygromycin (InvivoGen). iSLK cells were maintained in DMEM (Gibco) supplemented with 10% FBS, 1% penicillin-streptomycin (Gibco), 250 µg/mL G418 (Gibco), and 1200 µg/mL hygromycin (InvivoGen). iSLK.219 and iSLK cells were reactivated with 0.5 µg/mL doxycycline (Sigma-Aldrich). HEK293T cells were cultured in DMEM (Gibco) supplemented with 10% FBS and 1% penicillin-streptomycin (Gibco). For all inducible cell lines, the Tet System approved FBS (Takara Bio) was used. The identity of cell lines was confirmed by high-throughput RNA-seq and analysis of reads mapping to the KSHV genome, and the absence of mycoplasma contamination was verified using MycoAlert Mycoplasma Detection Kit (Lonza).

### RNA isolation and RT-qPCR

Total cell RNA was isolated using mirVana miRNA Isolation Kit (Invitrogen). cDNA was synthesized using SuperScript IV Reverse Transcriptase (Invitrogen) and then qPCR was performed using LightCycler 480 SYBR Green I Master Kit (Roche) with LightCycler 480 System (Roche). mRNA expression levels were calculated by the 2^−ΔΔCt^ method relative to *GAPDH* and normalized to control samples. The ΔCt values were used for the statistical analyses. RT-qPCR primers are listed in Supplementary Table 1 and 6.

### Western blotting

Whole-cell lysates were prepared in radioimmuno-precipitation assay (RIPA) buffer (Boston BioProducts) supplemented with phosphatase inhibitor (Sigma-Aldrich) and protease inhibitor mixtures (Thermo Scientific) and sonicated to shear DNA prior to centrifugation for 30 min at 16,000 rpm at 4 °C. Total protein concentrations were determined using Pierce BCA Protein Assay Kit (Thermo Scientific). Equal amounts of protein samples prepared in 4x Laemmli sample buffer (BIO-RAD) were resolved by 12% sodium dodecyl sulfate-polyacrylamide gel electrophoresis (SDS-PAGE), transferred onto nitrocellulose membranes (BIO-RAD), and blocked with blocking buffer (5% wt/vol milk, TBST) for 1 h at room temperature (RT). Membranes were incubated with primary antibodies with gentle agitation overnight at 4 °C and then incubated with horseradish peroxidase (HRP)-linked secondary antibodies for 1 h at RT. Protein bands were visualized by applying Clarity ECL Western Blotting Substrates (BIO-RAD) and using a gel imaging system or film. Relative protein levels were quantified using the Fiji software compared to β-actin and normalized to control samples in relevant graphs. All antibodies used are listed in Supplementary Table 1.

### Antibodies

The following antibodies and dilutions were used for western blotting: KSHV ORF26 (NOVUS no. NBP1-47357, 1:500), KSHV ORF45 (NOVUS no. NBP2-37685, 1:500), KSHV ORF50 (Abbiotec no. 251345, 1:500), KSHV K8 (NOVUS no. NB100-2189, 1:500), β-actin (Santa Cruz Technology no. sc-47778, 1:1,000), Anti-rabbit IgG, HRP-linked (CST no. 7074, 1:2,000), and Anti-mouse IgG, HRP-linked (CST no. 7076, 1:2,000).

### Transfection

*mir*Vana miRNA mimics (has-miR-31-5p, assay ID: MC11465; Negative Control No. 1) were purchased from Invitrogen. Silencer Select siRNAs (KHDRBS #1, assay ID: s20950; KHDRBS3 #2, assay ID: s20949; SLC51A #1, assay ID: s225837; SLC51A #2, assay ID: s47293; Negative Control No. 1) were purchased from Thermo Scientific. Cells were transfected with the indicated miRNA mimics (2.5 or 5 nM each) or siRNAs (40 nM each) using the Lipofectamine RNAiMAX Transfection Reagent (Invitrogen). At 72 h after transfection, cells were harvested for total RNA and total protein extraction. Downstream functional assays and analyses were performed at 72 h posttransfection.

### Dual-luciferase reporter assay

The full-length 3′ UTRs of *KHDRBS3*, *SLC51A* and *NNMT* were PCR-amplified from human genomic DNA using the primers listed in Supplementary Table 6. This product was cloned into the psiCHECK-2 vector (Promega) using *NotI* and *XhoI* restriction sites and verified by Sanger sequencing. The miRNA target sites in the WT 3’ UTRs of *KHDRBS3* and *SLC51A* were mutated using QuikChange Lightening Site-Directed Mutagenesis Kit (Agilent) and the mutagenic primers listed in Supplementary Table 6. Sanger sequencing confirmed successful mutagenesis of the miRNA target sites. psiCHECK-2 vector (Promega) containing either a WT or mutant 3’ UTRs of the indicated genes was cotransfected with miRNA mimic of interest or with NC mimic (Invitrogen) into iSLK.219 cells using the Lipofectamine 3000 Transfection Reagent (Invitrogen). The Renilla and firefly luciferase activities were measured 72 h posttransfection using the Dual-Luciferase Reporter System (Promega) following the manufacturer’s instructions. Renilla signals were normalized to an internal firefly luciferase as an internal control for transfection efficiency.

### KSHV lytic induction and fluorescence microscopy

For KSHV reactivation, iSLK.219 cells transfected with the indicated miRNA mimics or siRNAs were treated with 0.5 µg/mL doxycycline (Sigma-Aldrich), and then collected 72 h post-Dox treatment for later downstream analyses. Representative GFP and RFP images of the cells were taken 72 h post-Dox treatment using an EVOS FL Cell Imaging System (Life Technologies). RFP-positive iSLK.219 cells lytic reactivated with doxycycline (Sigma-Aldrich) were quantified using the Celigo imaging cytometer (Nexcelom Bioscience) at the indicated time points.

### De novo infection of KSHV in HEK293T cells

For supernatant transfer, iSLK.219 cells transfected with the indicated miRNA mimics or siRNAs were induced with 0.5 µg/mL doxycycline (Sigma-Aldrich) to produce infectious KSHV virions. After 96 h of induction, cell-free virus containing supernatants were collected and used to quantify virions or infect naive HEK293T cells. At 48 h post-infection, GFP-positive HEK293T cells infected with rKSHV.219 were imaged using an EVOS FL Cell Imaging System (Life Technologies) and quantified using the Celigo imaging cytometer (Nexcelom Bioscience).

### Quantification of KSHV genome copy number

Cell-free virus containing supernatants were centrifuged through a 20% sucrose solution at 25,000 rpm for 90 min at 4 °C to obtain concentrated virus, and then the viral DNA was isolated from the supernatants using DNAzol^®^ (Molecular Research Center). KSHV genome copy numbers were quantified by qPCR using LightCycler 480 SYBR Green I Master Kit (Roche) and LANA N-terminus primers with LightCycler 480 system (Roche) based on a standard curve constructed from known amounts of pcDNA3.1 plasmid containing ORF73/LANA.

### Subcellular fractionation

The cells were harvested and lysed in 350 µL ice cold Hypotonic Lysis Buffer supplemented with 100U of RNase OUT RNase inhibitor, by incubating on ice for 10 minutes followed by centrifugation at 1000 g at 4°C for 3 min. The supernatant (cytoplasmic fraction) was transferred to a separate tube. The remaining pellet (nuclear fraction) was washed with ice cold HLB buffer and centrifuged at 300g at 4°C for 2 min; this step was repeated three times and then total RNA was isolated from the cytoplasmic and nuclear fractions using mirVana miRNA Isolation Kit (Invitrogen). The purity of the nuclear fraction was verified by *MALAT1* RNA levels.

### RT-qPCR-based alternative splicing analysis for *ORF50*, *ORF57*, and *K8.1*

Primers used for detection of *ORF50* alternatively spliced transcripts were designed such that at least one primer from a pair spans spliced regions, such as the exon-exon junctions if possible, to ensure specific detection of specific alternatively spliced isoforms and the same reverse primer located in the E2 exon (ORF50 E2 R) shared by all the isoforms was used for RT-qPCR. *N3* and *N5* spliced mRNA isoforms were detected using two isoform-specific primer pairs (See Supplementary Fig. 5a for schematics and the primer pairs). N4 containing isoforms were detected using a forward primer specific to N4 (ORF50 N4 F) and the reverse primer located in the E2 exon (ORF50 E2 R). This primer set detects both the spliced and unspliced *N4* isoforms. The unspliced pre-mRNA transcripts were quantified using the same N4 forward primer (ORF50 N4 F) and a reverse primer located in the intron (ORF50 N4 Intronic R). Since the E1 and N4 exons share an exon-exon junction, a forward primer located in the N4/E1-E2 exon-exon junction (ORF50 N4/E1_E2 F) and the reverse primer (ORF50 E2 R) were used to detect both the spliced *N4* and the spliced *E1* isoforms. Total *ORF57* transcripts were detected with a forward primer located in exon 1 (ORF57 Total F) and a reverse primer located in exon 2 (ORF57 Total R). *ORF57* spliced mRNA isoforms ("ORF57 1/2") were detected with a forward primer located at the exon-exon junction between exon 1 and exon 2 common to both spliced isoforms (ORF57 E1_E2 F) and the same reverse primer used to detect total *ORF57* transcripts (ORF57 Total R), whereas the unspliced transcripts were detected with a forward primer within the intron (ORF57 UnSp F) and a reverse primer at the exon-intron junction of exon 2 (ORF57 UnSp R) as indicated in Supplementary Fig. 5b. The longer *ORF57* isoform with a longer exon 2 ("ORF57 LE2") was detected with a forward primer located in a region of exon 2 shared by both *ORF57* spliced isoforms (ORF57 E2 F) and a reverse primer located near the 3’ end of the longer exon 2 (ORF57 LE2 R), which is downstream of the shorter *ORF57* isoform with a third exon ("ORF57 2/3") and thus unique to the *ORF57 LE2* isoform. This primer pair detects both the unspliced and spliced *ORF57 LE2* isoforms. The shorter *ORF57* isoform with exon 3 ("ORF57 2/3") was quantified using the same forward primer used to detect the *ORF57 LE2* isoform (ORF57 E2 F) and a reverse primer located within the junction of exon 2 and exon 3 (ORF57 E2_E3 R). Unspliced *K8.1* transcripts were detected using a forward primer located within the intron (K8.1 UnSp F) and a reverse primer in exon 2 (K8.1 E2 R). The long and short *K8.1* spliced mRNA isoforms were detected using two isoform-specific forward primers that span their unique exon-exon junctions (K8.1 Short F and K8.1 Long F) and the same reverse primer in exon 2 (K8.1 E2 R) (See Supplementary Fig. 5c for schematics and the primer pairs). The specificity of the PCR primers used was verified by their qPCR melting curve and their production of specific single amplicons. Of note, *K8.1* is located within another transcript, *K8*, and a previous study reported novel bicistronic *K8/K8.1* mRNAs with little or no protein-coding capacity^47^. Thus, the primers used for detection of *K8.1* would detect both transcripts. PCR primer sequences are provided in Supplementary Table 6.

### High-throughput sequencing

Total RNA was isolated using *mir*Vana miRNA Isolation Kit (Invitrogen) following the manufacturer’s protocol. All RNA samples were analyzed for RNA integrity using the Agilent 2100 Bioanalyzer. For miRNA-seq, small RNA library preparation was performed using Illumina TruSeq small RNA sample preparation protocol and sequenced as 50-nucleotide (nt) single-end runs on the HiSeq 2500 platform. Raw reads were subjected to an in-house program, ACGT101-miR (LC Sciences) to remove adapter dimers, junk, low complexity, common RNA families (rRNA, tRNA, snRNA, snoRNA) and repeats. Subsequently, the remaining 18 to 26 nucleotide reads were mapped to the microRNA database (miRBase) version 22.0 (https://www.mirbase.org) by basic local alignment search tool (BLAST) to identify known and novel miRNAs. The differential expression analysis was performed using DESeq2 version 1.26.0^95^. For digital gene expression (DGE)-seq, libraries were prepared following Illumina small RNA sample preparation protocol and sequenced as 50-nt single-end runs on the HiSeq 2500 platform. Single-end reads were mapped to the KSHV reference genome (NC_009333) or the human genome (Ensembl GRCh37) using Bowtie2 version 2.2.9^96^. The abundance of gene expression was estimated using RNA-Seq by Expectation Maximization (RSEM) version 1.3.0^97^, and the differential expression analysis was performed using edgeR version 3.12.1^98^. Genes showing significant differences (log2 CPM >0 and |log2 FC| >1) were selected for enrichment analysis using GAGE version 2.20.1^99^.

### Statistical analysis

Statistical significance was determined by two-tailed Student’s *t*-test and *p*-value *<* 0.05 was considered statistically significant. All results represent mean ± SEM and were analyzed by statistical tests as indicated in the figure legends. *p*-values are: \**p* < 0.05, \*\**p* < 0.01, \*\*\**p* < 0.001, \*\*\*\**p* < 0.0001, and ns = not significant. The details on the statistical analysis of the small-RNA sequencing and RNA-seq data can be found in the *Materials and Methods* section. For analysis of KHDRBS3 expression in KS patient tissues, we used the published RNA-seq data (GSE147704)^85^ and the raw counts for KHDRBS3 gene were normalized according to a scaling normalization method^100^ with statistical significance of differential expression determined by two-tailed Student’s *t*-test.

### Data availability

The small-RNA sequencing and RNA sequencing data generated during this study are uploaded at Gene Expression Omnibus. All software used in this study is listed in Supplementary Table 1.

## Acknowledgements

We thank Jonathan Dreyfuss, of the Harvard Catalyst Biostatistical Consulting Program for providing biostatistics consultation. We acknowledge the Ludwig Center at Harvard and the NCI Outstanding Investigator Award (R35CA232105) to F.J.S. K.M.K acknowledges support from the following NIH grants (AI165382, AI150575, DE025208).

## Author Contributions

Conceptualization: S.M.L., K.M.K., F.J.S.

Investigation: S.M.L., C.L.A., C.M., H.M.D., E.C., F.J.S.

Resources: K.M.K, F.J.S.

Supervision: K.M.K., F.J.S.

Writing - review & editing: S.M.L., F.J.S.

## Supplementary Information

**Supplementary Figure 1.**
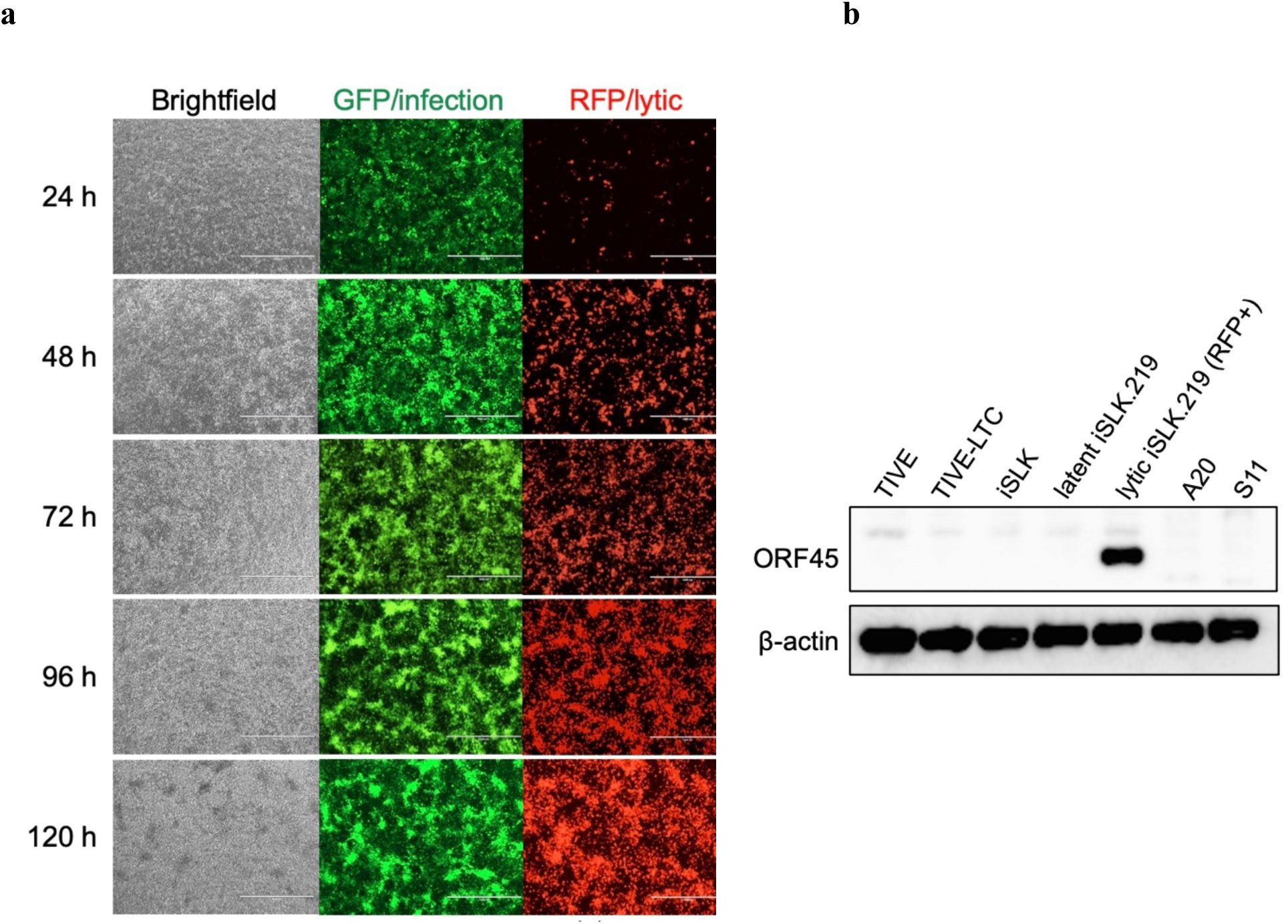
iSLK.219 cells with dox-inducible RTA expression undergo efficient lytic reactivation upon Dox treatment. (**a**) Representative images showing phase, GFP and RFP fluorescence of iSLK.219 cells over a 120 h Dox-induced lytic reactivation time course, with samples collected at 24, 48, 72, 96, and 120 h. iSLK.219 cells harbor a recombinant KSHV (rKSHV.219) genome with constitutive GFP expression to indicate latent infection, and induced RFP expression upon lytic reactivation to indicate lytic infection. (**b**) Western blot analysis of immediate-early KSHV lytic protein ORF45 expression in the indicated cell lines, including latent iSLK.219 cells at 0 h and lytic iSLK.219 cells at 72 h post Dox-induced KSHV lytic reactivation. β-actin was used as a loading control.

**Supplementary Figure 2.**
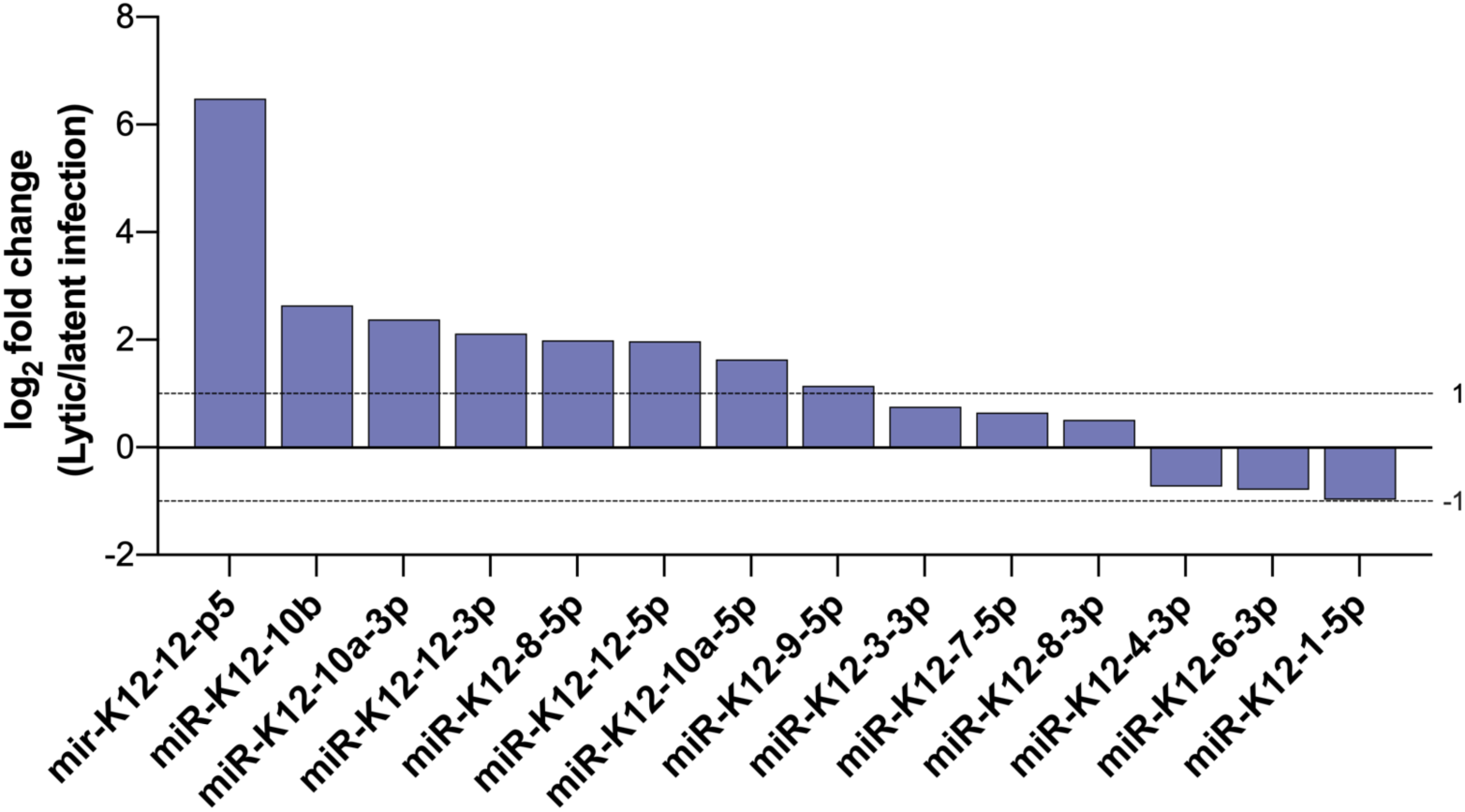
Changes in KSHV miRNA expression in iSLK.219 cells during KSHV lytic reactivation, as determined by small-RNA sequencing. Waterfall plot depicting KSHV-encoded miRNAs with a log_2_ fold change value ≥ |0.5| in iSLK.219 cells with lytic (Dox+) compared to latent (Dox-) KSHV infection. Total RNA isolated from latent iSLK.219 cells at 0 h or lytic iSLK.219 cells at 72 h post Dox-induced KSHV lytic reactivation was subjected to small-RNA sequencing analysis. mir-K12-12-p5 is the miR-K12-12 precursor (pre-miR-K12-12).

**Supplementary Figure 3.**
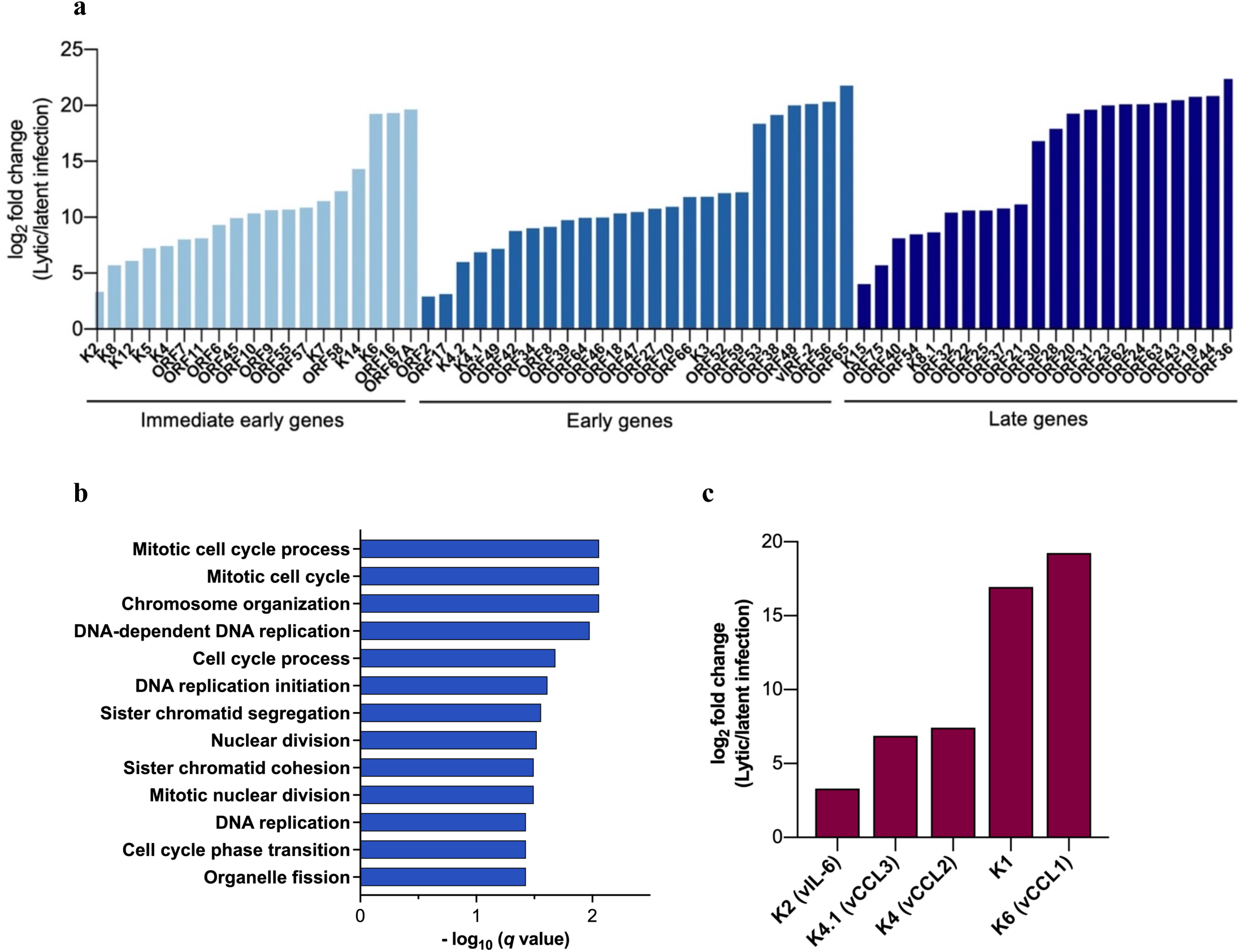
Changes in viral and cellular mRNA expression in iSLK.219 cells during KSHV lytic reactivation, as determined by RNA sequencing. (a) RNA sequencing analysis of 66 KSHV open reading frames (ORFs) in iSLK.219 cells with lytic (Dox+) compared to latent (Dox-) KSHV infection. (**b**) Gene ontology (GO) analysis was performed to analyze all genes significantly up-regulated in lytic iSLK.219 cells (Dox+) compared to latent iSLK.219 cells (Dox-) at 72 h post Dox-induced lytic reactivation, as determined by RNA-seq, identifying significantly enriched GO terms. (**c**) Expression of KSHV lytic genes with mitogenic signaling activities, K2 (vIL-6), K4.1 (vCCL3), K4 (vCCL2), K1, and K6 (vCCL1), in iSLK.219 cells with lytic (Dox+) compared to latent (Dox-) KSHV infection. Total RNA was isolated from latent iSLK.219 cells at 0 h or lytic iSLK.219 cells at 72 h post Dox-induced KSHV lytic reactivation was subjected to RNA sequencing analysis.

**Supplementary Figure 4.**
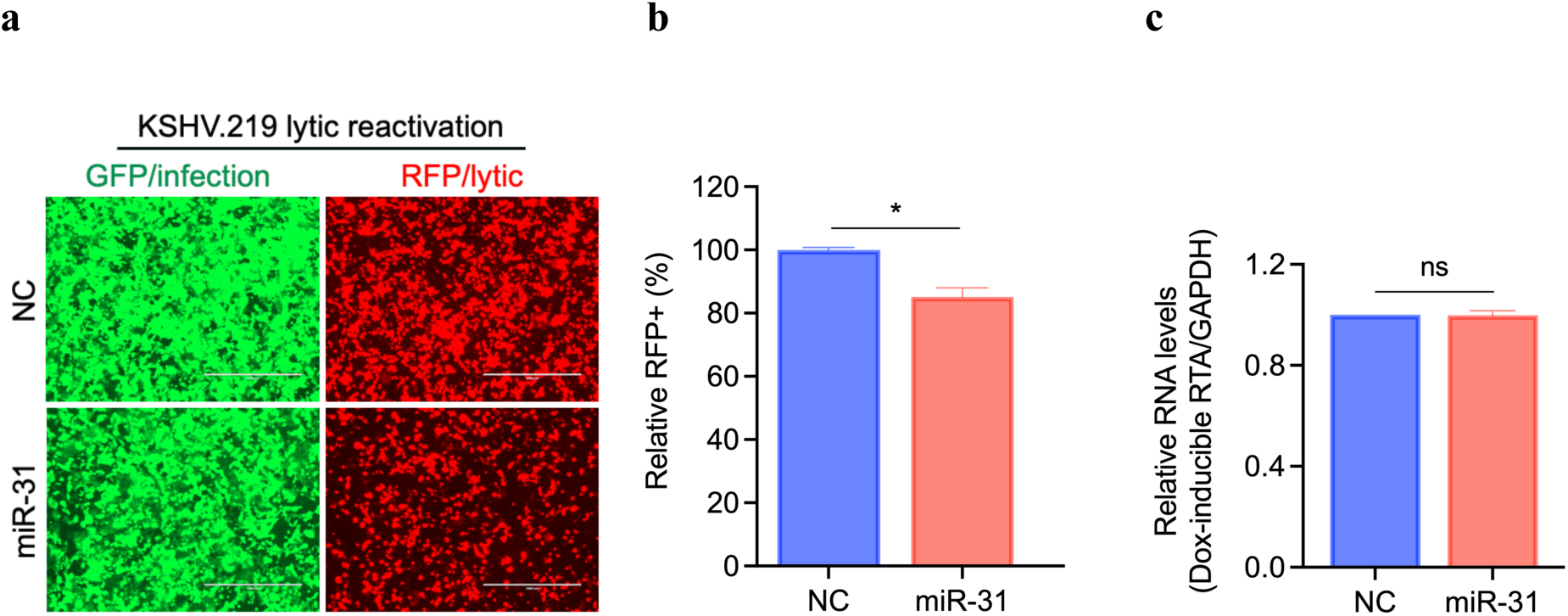
Ectopic miR-31 expression suppresses KSHV lytic reactivation. **(a)** iSLK.219 cells were transfected with negative control (NC) or miR-31 mimic followed by treatment with Dox (0.5 µg/mL) for 72 h to induce KHSV lytic reactivation, and representative images were photographed for GFP and RFP fluorescence at 72 h post Dox-induced lytic reactivation (Scale bars, 1000 µm). (**b**) Quantification of RFP-positive cells in (A) using image cytometry. Two biological replicates are presented. (**c**) iSLK cells were transfected with negative control (NC) or miR-31 mimic followed by treatment with Dox (0.5 µg/mL) for 72 h. Total RNA was isolated at 72 h post-Dox treatment and expression of Dox-induced *RTA* was quantified by RT-qPCR. *GAPDH* was used as an endogenous control. Two biological replicates are presented. Data represent mean ± SEM and *p*-values were determined by two-tailed Student’s *t*-test. \**p* < 0.05, \*\**p* < 0.01, \*\*\**p* < 0.001, \*\*\*\**p* < 0.0001, and ns = not significant.

**Supplementary Figure 5.**
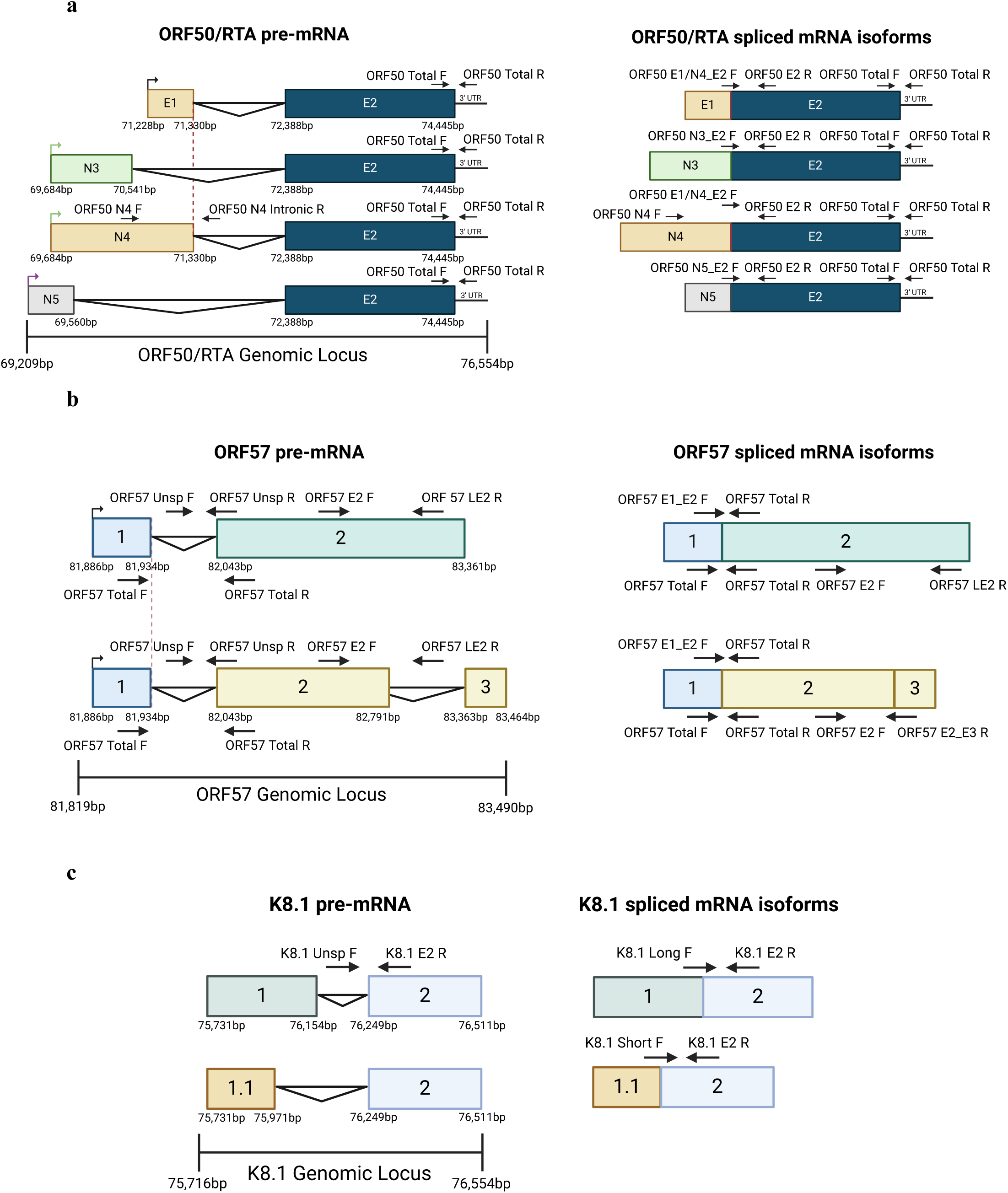
Analysis of alternative splicing in the immediate-early lytic ORF50, early lytic ORF57, and late lytic K8.1 KSHV genes. (**a-c**) Schematic representation of alternatively spliced transcripts of *ORF50* (**a**), *ORF57* (**b**), and *K8.1* (**c**) in the context of the corresponding gene, focusing on alternatively spliced exons. The region of mRNA that demonstrates the alternative splicing events is depicted in greater detail. Colored boxes represent exons and the horizontal black lines represent introns. The arrowheads depict the approximate RT-qPCR primer location. PCR primer sequences are provided in Supplementary Table 6.

**Supplementary Figure 6.**
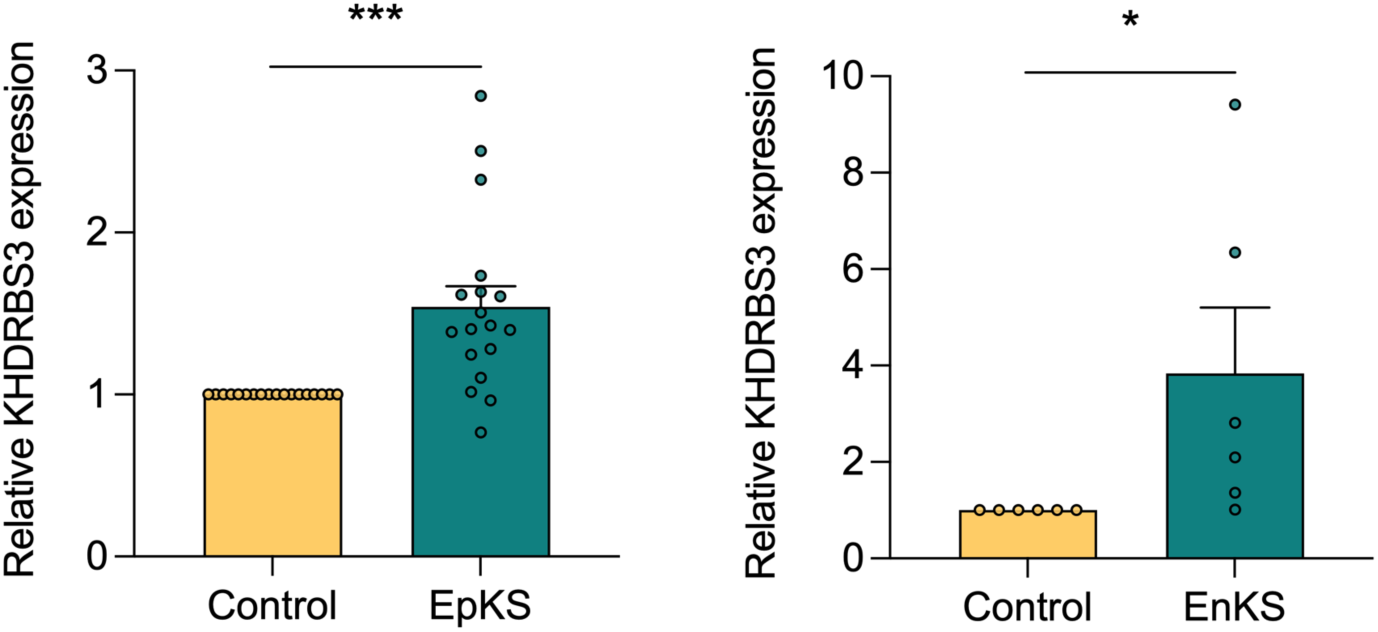
RNA-seq analysis of KHDRBS3 and miR-31-5p expression in KS lesions. *KHDRBS3* expression in 18 pairs of AIDS-associated/epidemic KS (EpKS) (Left) or 6 pairs of non-AIDS associated/endemic KS (EnKS) (Right), relative to matched uninvolved control tissues from the published RNA-seq data^1^. Data represent mean ± SEM and *p*-values were determined by two-tailed Student’s *t*-test. \**p* < 0.05, \*\**p* < 0.01, \*\*\**p* < 0.001, \*\*\*\**p* < 0.0001, and ns = not significant.

**Supplementary Table 1.**
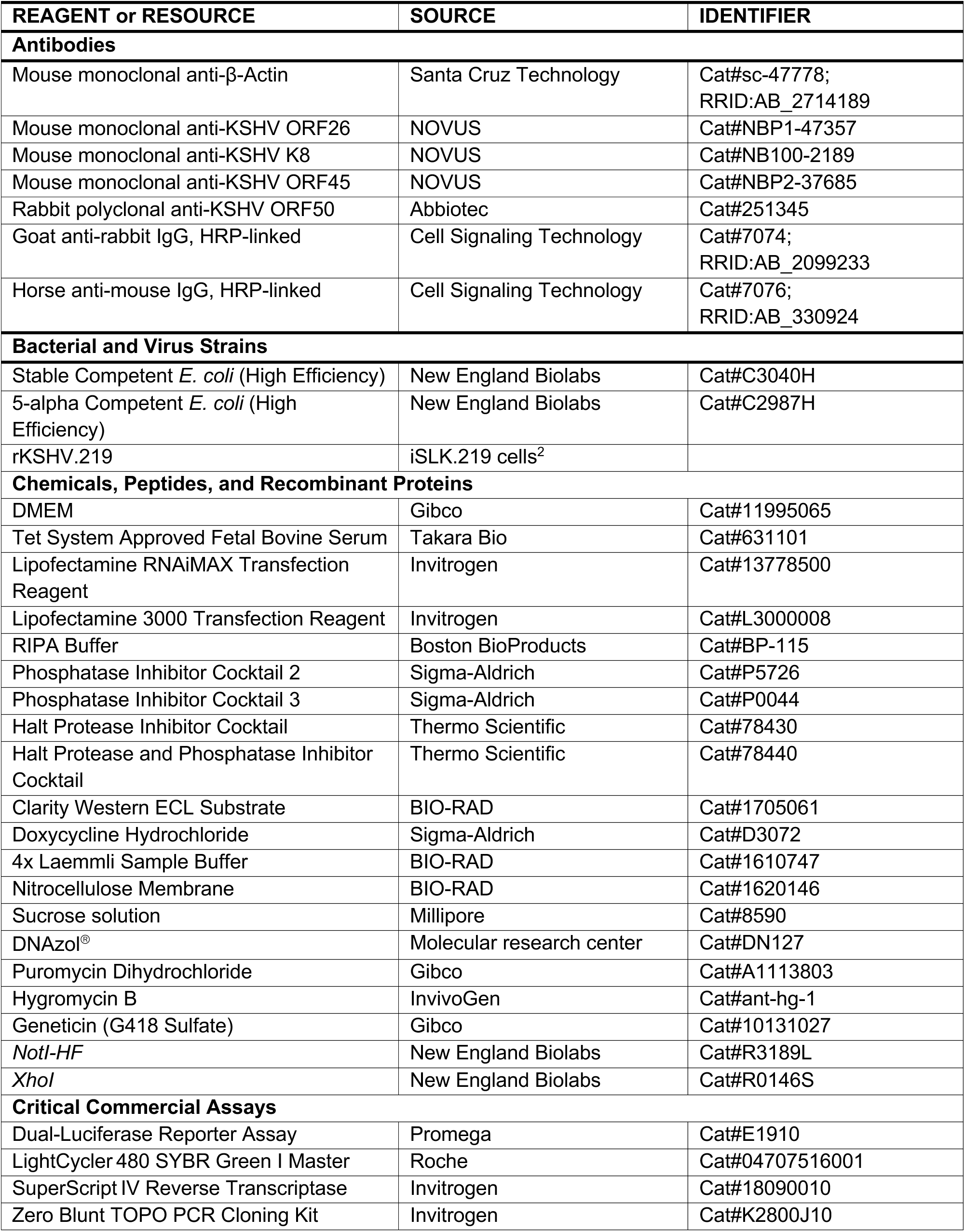
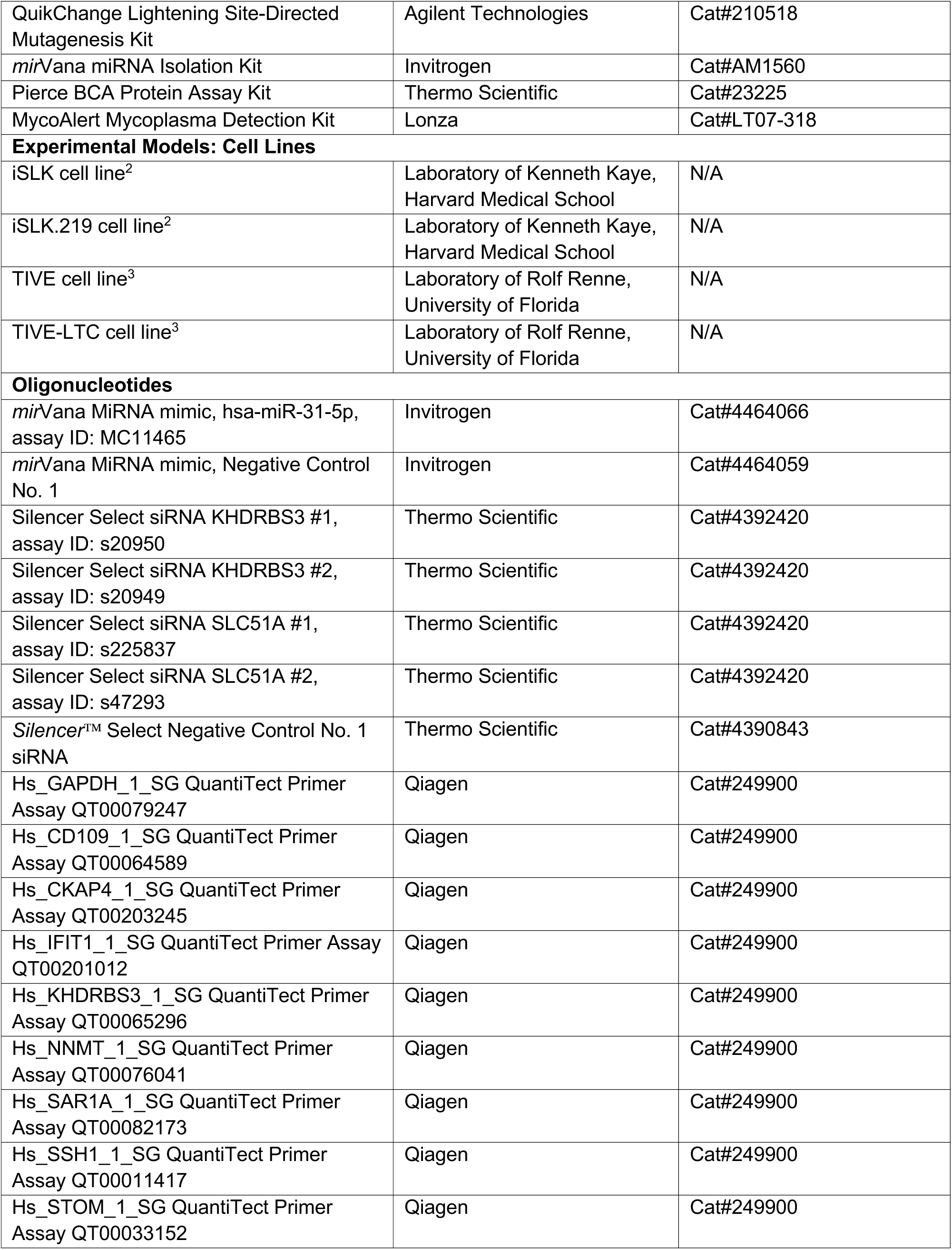
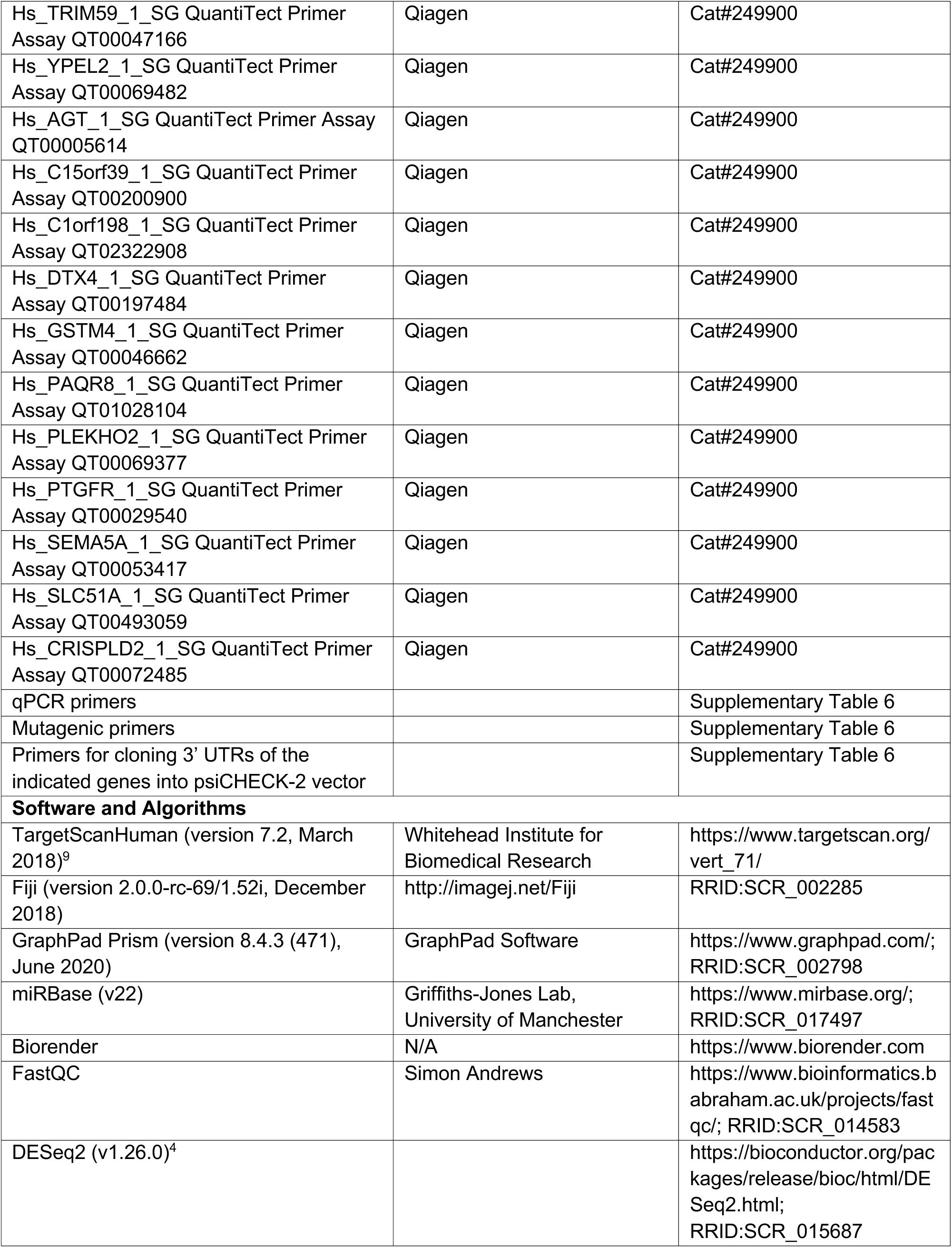
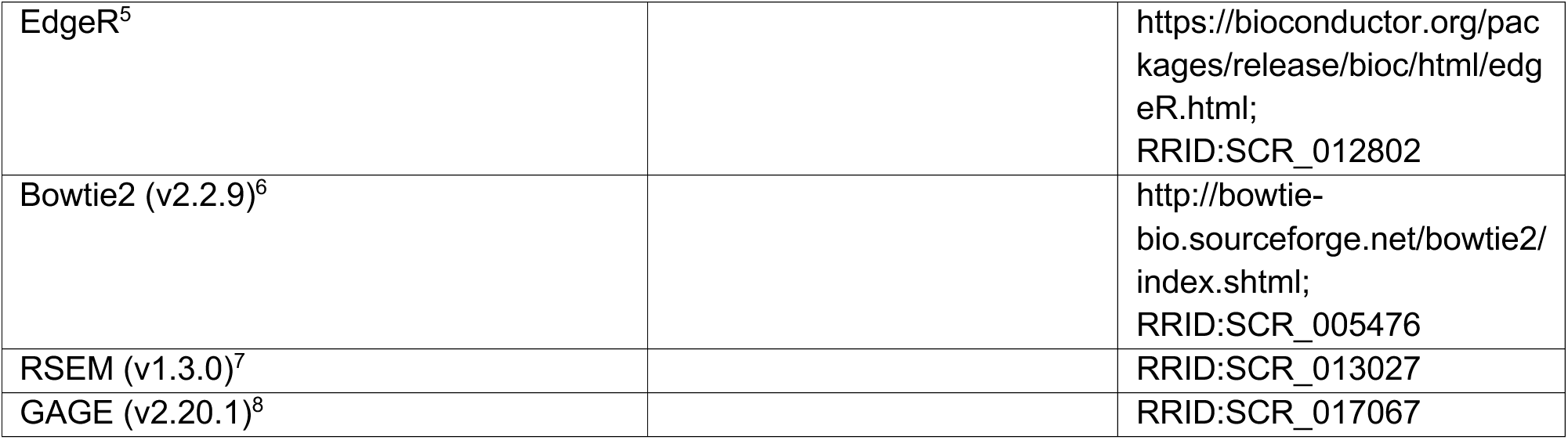
Reagent sources.

**Supplementary Table 2.**
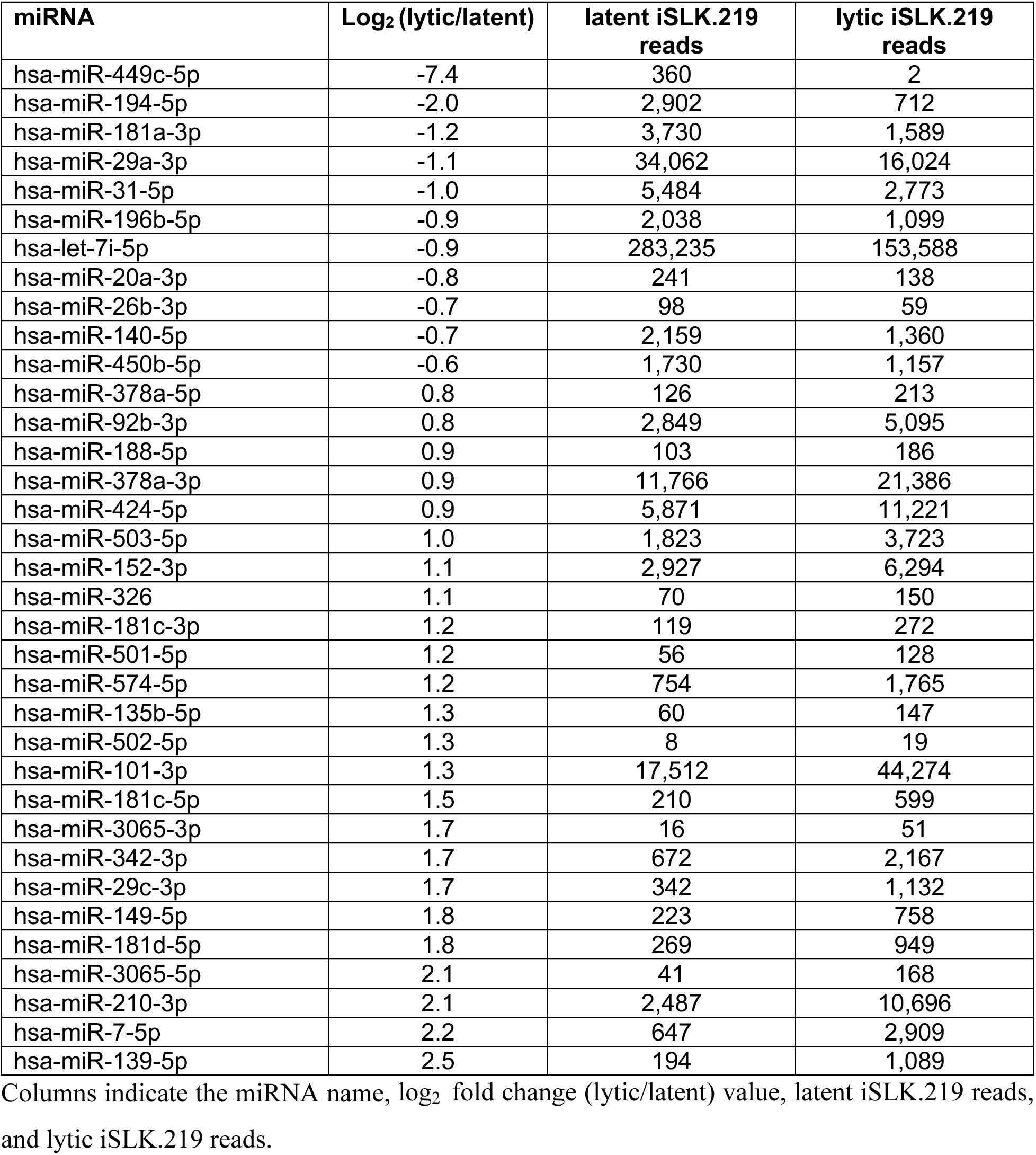
A list of differentially expressed cellular miRNAs identified by small-RNA sequencing analysis of iSLK.219 cells with lytic (Dox+) compared to latent (Dox-) KSHV infection (log_2_ fold change ≥ |0.6| and adjusted p-value < 0.05). These miRNAs are ranked according to the log2 fold change (lytic/latent) value.

**Supplementary Table 3.**
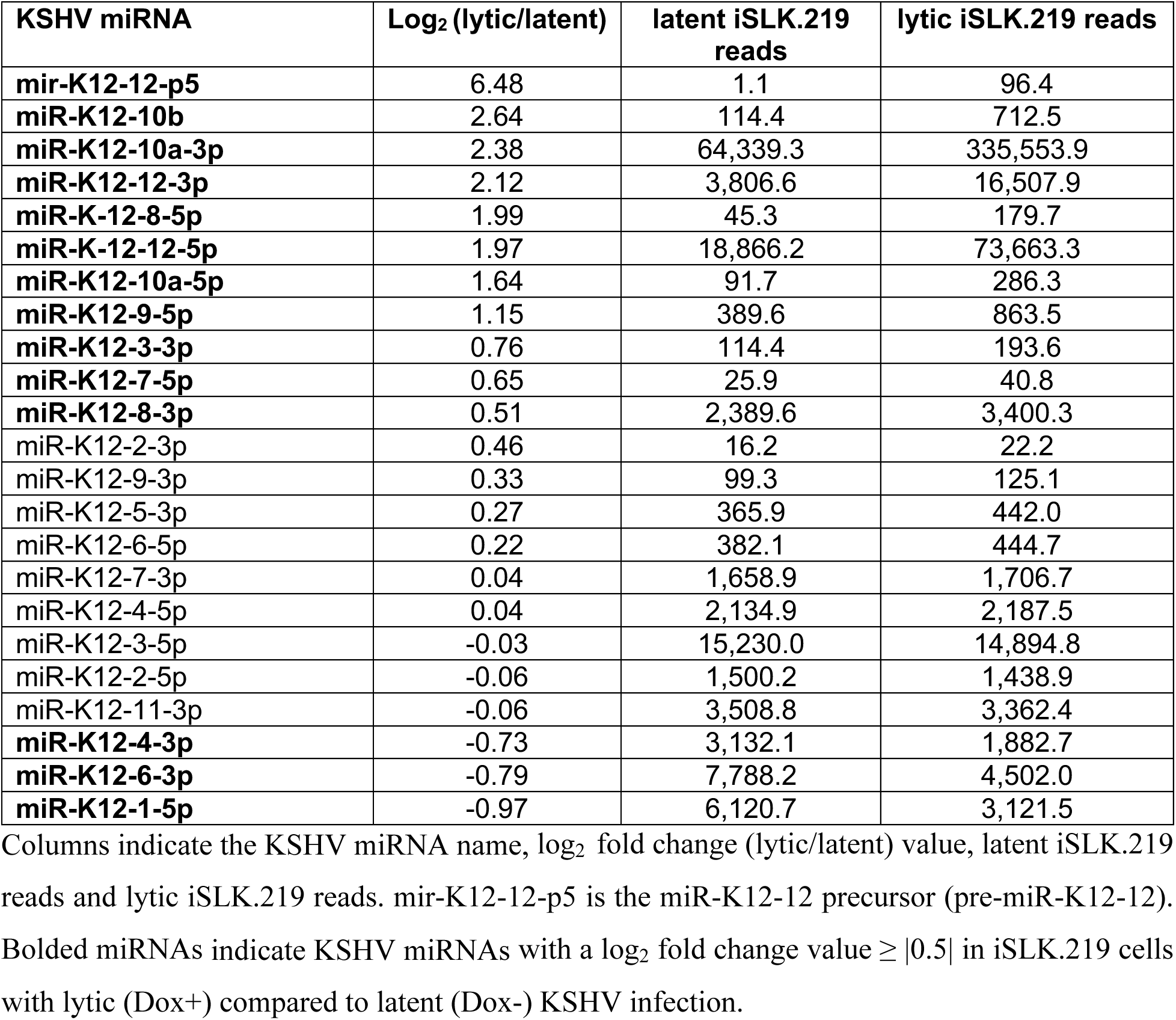
A list of differentially expressed KSHV-encoded miRNAs identified by small-RNA sequencing analysis of iSLK.219 cells with lytic (Dox+) compared to latent (Dox-) KSHV infection. These KSHV-encoded miRNAs are ranked according to the log_2_ fold change (lytic/latent) value. KSHV miRNAs in the small-RNA seq data that have less than 10 reads are removed to avoid uncertain low counts.

**Supplementary Table 4.**
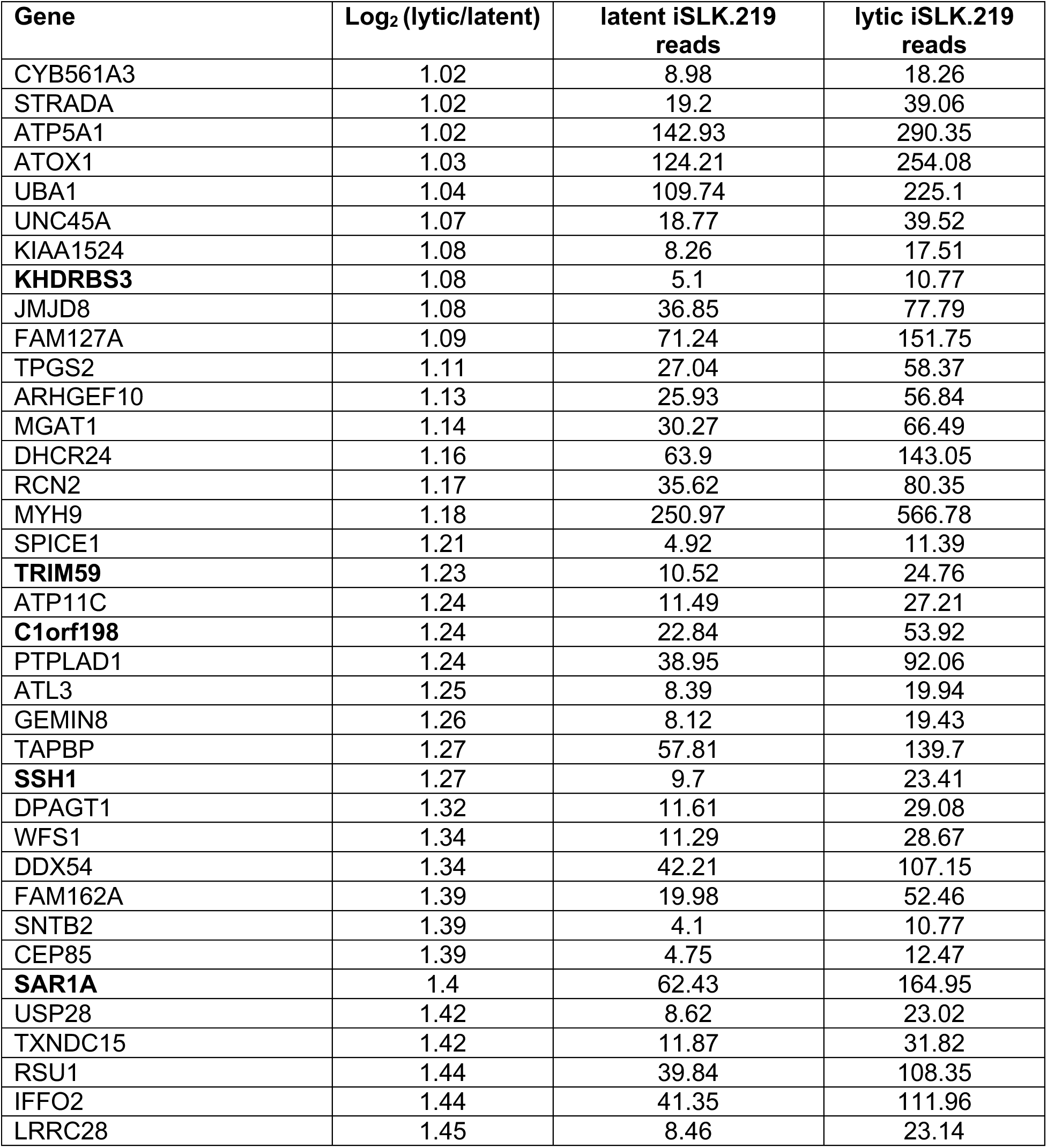
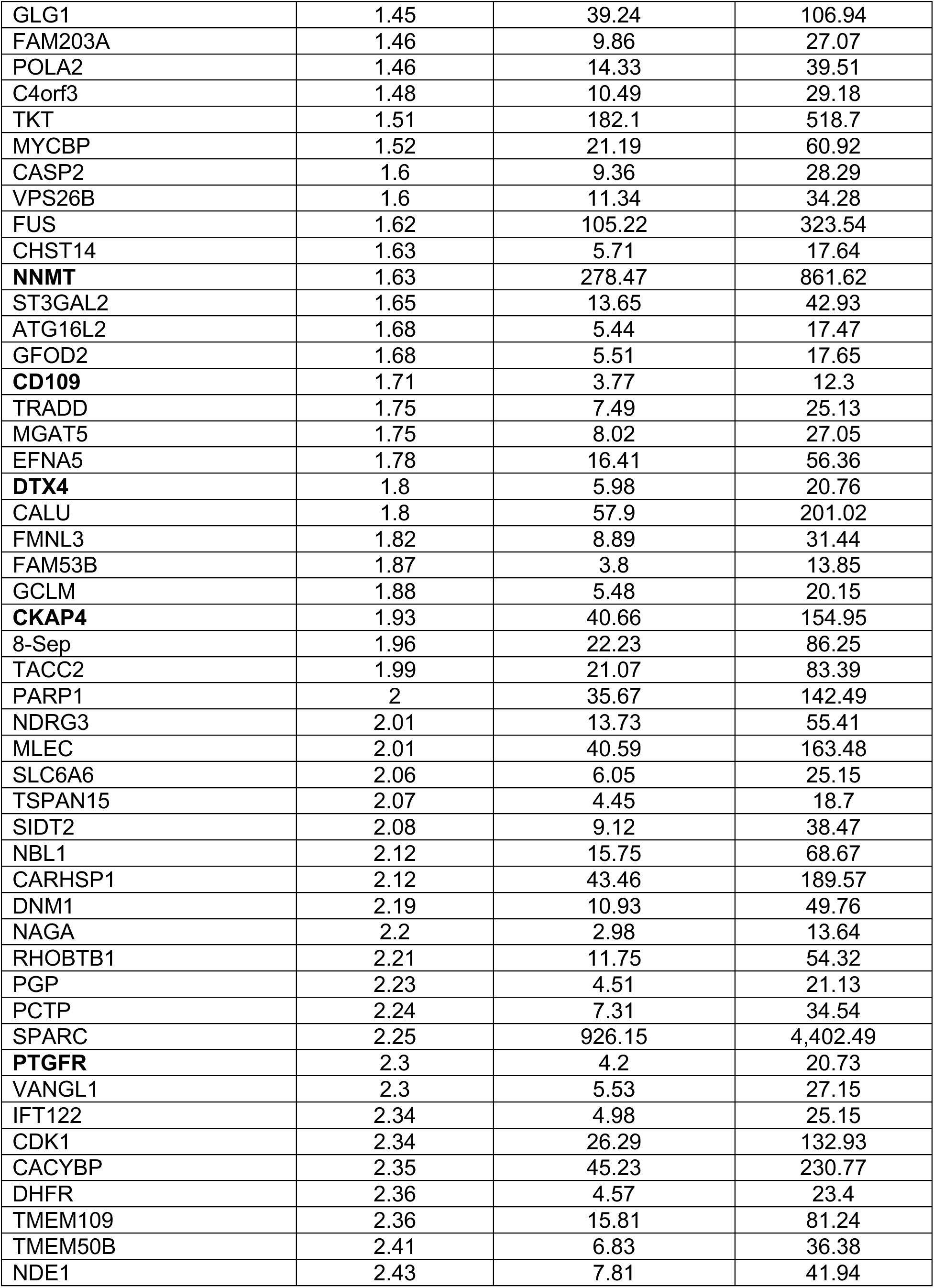
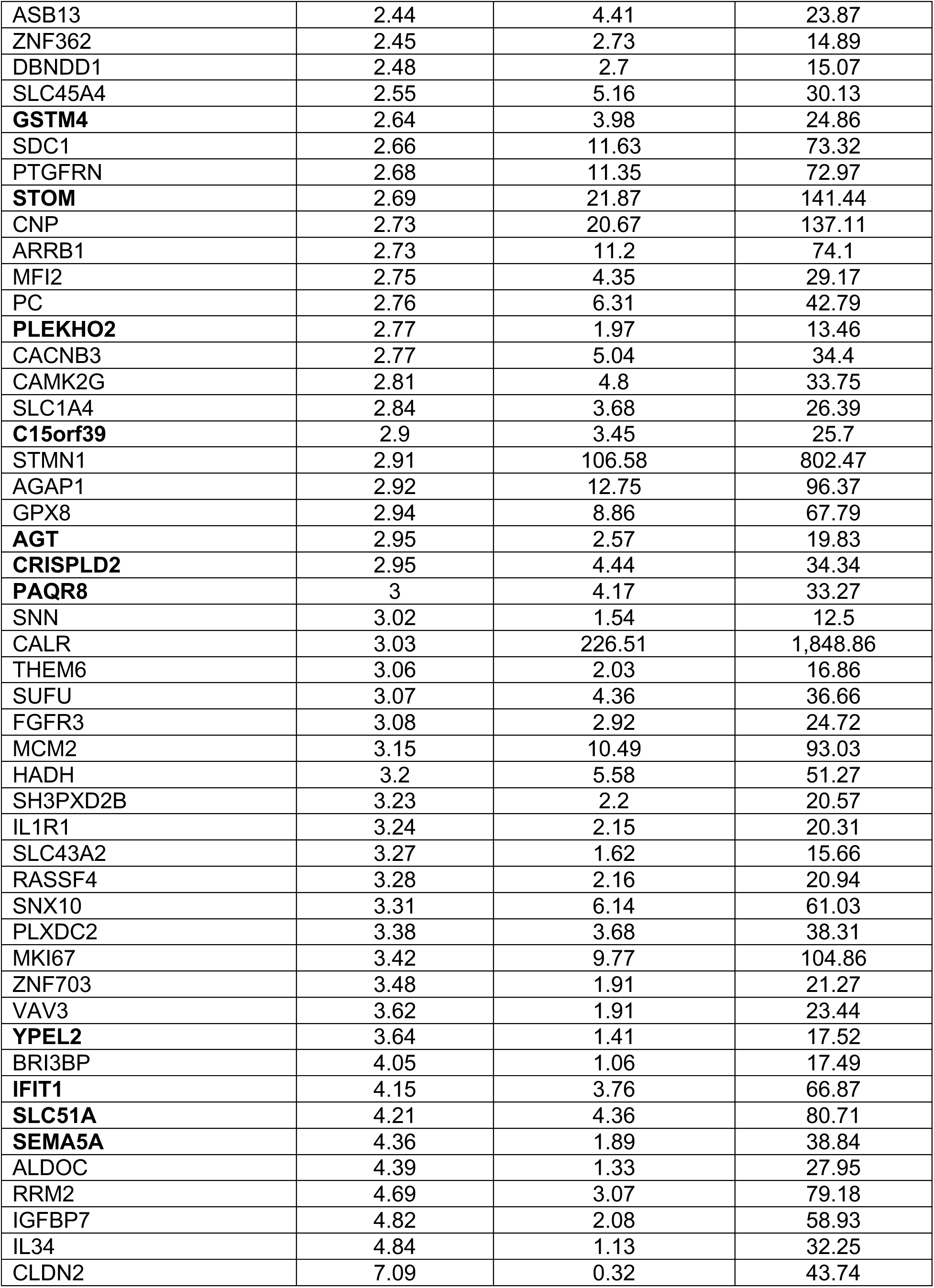
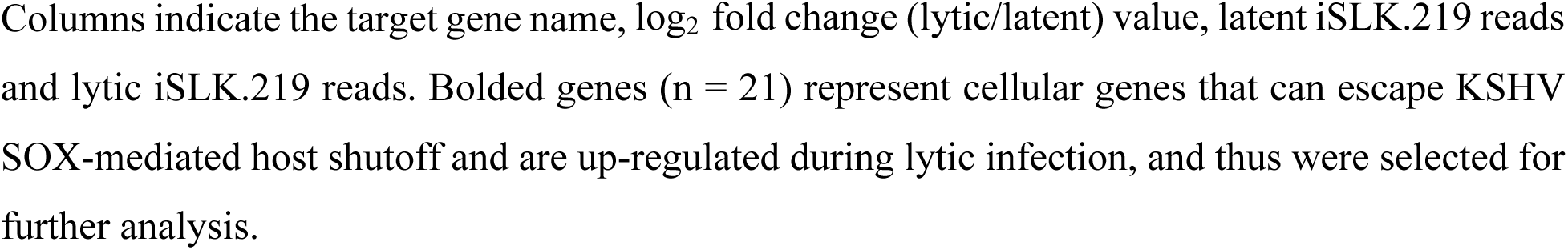
A list of TargetScan predicted miR-31-5p targets with conserved and poorly conserved sites and the log2 fold change in mRNA expression of these predicted targets in iSLK.219 cells with lytic (Dox+) compared to latent (Dox-) KSHV infection, as determined by RNA-seq. These predicted targets are ranked according to the log_2_ fold change (lytic/latent) value. Genes in the RNA-seq data that have less than 10 reads in lytic samples are removed to avoid uncertain low counts.

**Supplementary Table 5.**
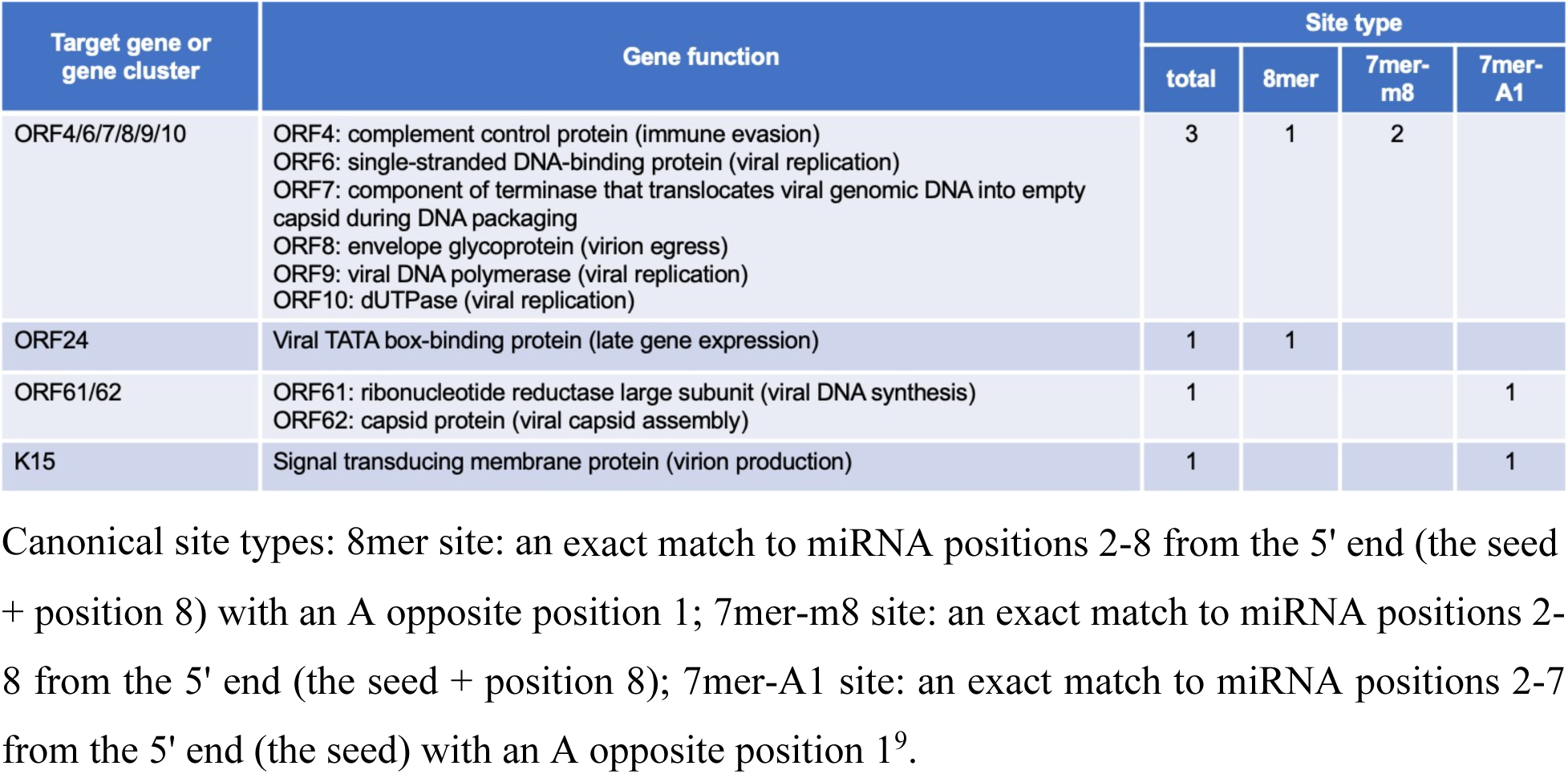
A list of predicted KSHV targets of miR-31-5p.

**Supplementary Table 6.**
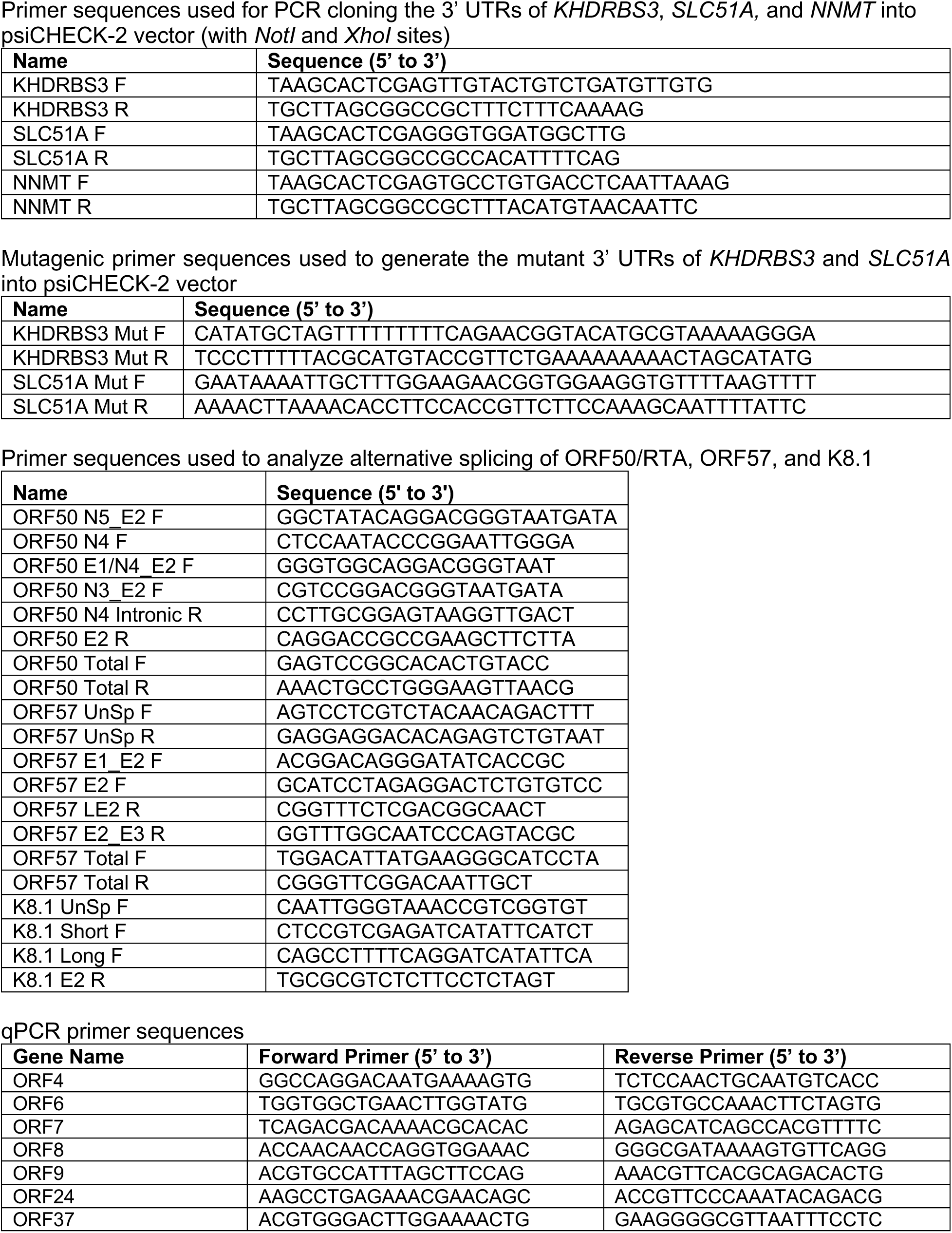
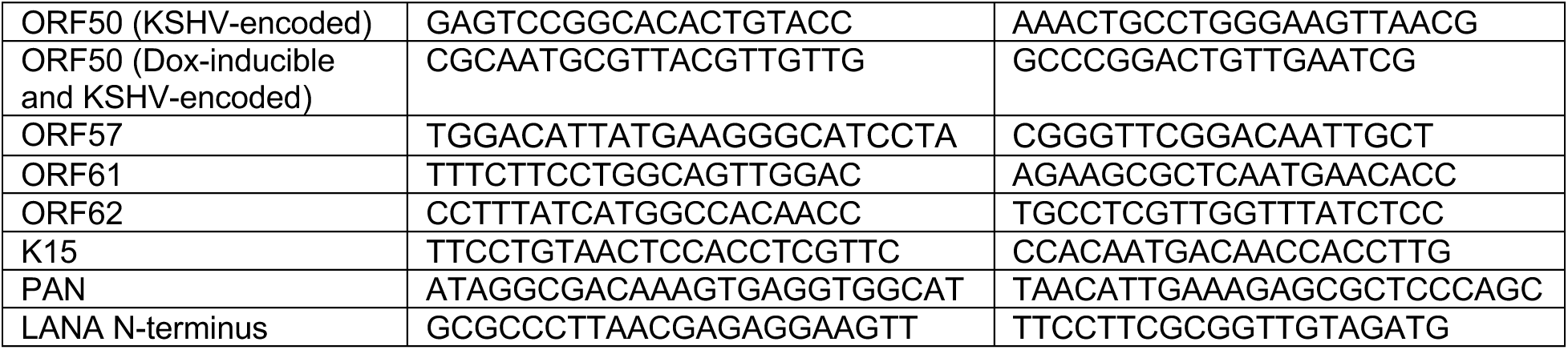
Oligonucleotides.

## References

1. Cesarman, E., Chang, Y., Moore, P. S., Said, J. W. & Knowles, D. M. Kaposi’s sarcoma-associated herpesvirus-like DNA sequences in AIDS-related body-cavity-based lymphomas. The New England Journal of Medicine, 332, 1186–1191 (1995).

2. Soulier, J., Grollet, L., Oksenhendler, E., Cacoub, P., Cazals-Hatem, D., Babinet, P., d’Agay, M. F., Clauvel, J. P., Raphael, M. & Degos, L. Kaposi’s sarcoma-associated herpesvirus-like DNA sequences in multicentric Castleman’s disease. Blood, 86, 1276–1280 (1995).

3. Ganem, D. KSHV and Kaposi’s sarcoma: the end of the beginning? Cell, 91, 157–160 (1997).

4. Yan, L., Majerciak, V., Zheng, Z. M. & Lan, K. Towards Better Understanding of KSHV Life Cycle: from Transcription and Posttranscriptional Regulations to Pathogenesis. Virologica Sinica, 34, 135–161 (2019).

5. Sztuba-Solinska, J. & Le Grice, S. F. J. The "Topological Train Ride" of a viral long non-coding RNA. RNA Biology, 15, 13–16 (2018).

6. Aneja, K. K. & Yuan, Y. Reactivation and Lytic Replication of Kaposi’s Sarcoma-Associated Herpesvirus: An Update. Frontiers in Microbiology, 8, 613 (2017).

7. Ma, Z., Ni, G. & Damania, B. Innate Sensing of DNA Virus Genomes. Annual Review of Virology, 5, 341–362 (2018).

8. Bellare, P. & Ganem, D. Regulation of KSHV lytic switch protein expression by a virus-encoded microRNA: an evolutionary adaptation that fine-tunes lytic reactivation. Cell Host & Microbe, 6, 570–575 (2009).

9. Gottwein E. Kaposi’s Sarcoma-Associated Herpesvirus microRNAs Frontiers in Microbiology, 3, 165 (2012).

10. Qin, J., Li, W., Gao, S. J. & Lu, C. KSHV microRNAs: Tricks of the Devil. Trends in Microbiology, 25, 648–661 (2017).

11. Giffin, L. & Damania, B. KSHV: pathways to tumorigenesis and persistent infection. Advances in Virus Research, 88, 111–159 (2014).

12. Mesri, E. A., Feitelson, M. A. & Munger, K. Human viral oncogenesis: a cancer hallmarks analysis. Cell Host & Microbe, 15, 266–282 (2014).

13. Bartel, D. P. Metazoan MicroRNAs. Cell, 173, 20–51 (2018).

14. Guo, Y. E. & Steitz, J. A. (2014). Virus meets host microRNA: the destroyer, the booster, the hijacker. Molecular and Cellular Biology, 34, 3780–3787 (2014).

15. Bruscella, P., Bottini, S., Baudesson, C., Pawlotsky, J. M., Feray, C. & Trabucchi, M. Viruses and miRNAs: More Friends than Foes. Frontiers in Microbiology, 8, 824 (2017).

16. Lee, S. M., Kaye, K. M. & Slack, F. J. Cellular microRNA-127-3p suppresses oncogenic herpesvirus-induced transformation and tumorigenesis via down-regulation of SKP2. Proceedings of the National Academy of Sciences of the United States of America, 118, e2105428118 (2021).

17. Fruscio, M. D., Chen, T. & Richard, S. Characterization of Sam68-like mammalian proteins SLM-1 and SLM-2: SLM-1 is a Src substrate during mitosis. Proceedings of the National Academy of Sciences of the United States of America, 96, 2710–2715 (1999).

18. Stoss, O., Olbrich, M., Hartmann, A. M., König, H., Memmott, J., Andreadis, A. & Stamm, S. The STAR/GSG family protein rSLM-2 regulates the selection of alternative splice sites. The Journal of Biological Chemistry, 276, 8665–8673 (2001).

19. Traunmüller, L., Gomez, A. M., Nguyen, T. M. & Scheiffele, P. Control of neuronal synapse specification by a highly dedicated alternative splicing program. Science, 352, 982–986 (2016).

20. Boeckel, J. N., Möbius-Winkler, M., Müller, M., Rebs, S., Eger, N., Schoppe, L., Tappu, R., Kokot, K. E., Kneuer, J. M., Gaul, S., Bordalo, D. M., Lai, A., Haas, J., Ghanbari, M., Drewe-Boss, P., Liss, M., Katus, H. A., Ohler, U., Gotthardt, M., Laufs, U., Streckfuss-Bömeke, K. & Meder, B. SLM2 is a novel cardiac splicing factor involved in heart failure due to dilated cardiomyopathy. Genomics, Proteomics & Bioinformatics, 20, 129–146 (2022).

21. Traunmüller, L., Schulz, J., Ortiz, R., Feng, H., Furlanis, E., Gomez, A. M., Schreiner, D., Bischofberger, J., Zhang, C. & Scheiffele, P. A cell-type-specific alternative splicing regulator shapes synapse properties in a trans-synaptic manner. Cell Reports, 42, 112173 (2013).

22. Reddy, T. R., Suhasini, M., Xu, W., Yeh, L. Y., Yang, J. P., Wu, J., Artzt, K. & Wong-Staal, F. A role for KH domain proteins (Sam68-like mammalian proteins and quaking proteins) in the post-transcriptional regulation of HIV replication. The Journal of Biological Chemistry, 277, 5778–5784 (2002).

23. Soros, V. B., Carvajal, H. V., Richard, S. & Cochrane, A. W. Inhibition of human immunodeficiency virus type 1 Rev function by a dominant-negative mutant of Sam68 through sequestration of unspliced RNA at perinuclear bundles. Journal of Virology, 75, 8203–8215 (2001).

24. Myoung, J. & Ganem, D. Generation of a doxycycline-inducible KSHV producer cell line of endothelial origin: maintenance of tight latency with efficient reactivation upon induction. Journal of Virological Methods, 174, 12–21 (2011).

25. Yang, X., Liang, Y., Bamunuarachchi, G., Xu, Y., Vaddadi, K., Pushparaj, S., Xu, D., Zhu, Z., Blaha, R., Huang, C. & Liu, L. miR-29a is a negative regulator of influenza virus infection through targeting of the frizzled 5 receptor. Archives of Virology, 166, 363–373 (2021).

26. Banerjee, A., Chawla-Sarkar, M. & Mukherjee, A. Rotavirus-Mediated Suppression of miRNA-192 Family and miRNA-181a Activates Wnt/β-Catenin Signaling Pathway: An In Vitro Study. Viruses, 14, 558 (2022).

27. Kim, Y. J., Yeon, Y., Lee, W. J., Shin, Y. U., Cho, H., Lim, H. W. & Kang, M. H. Analysis of MicroRNA Expression in Tears of Patients with Herpes Epithelial Keratitis: A Preliminary Study. Investigative ophthalmology & visual science, 63, 21 (2022).

28. Sisk, J. M., Witwer, K. W., Tarwater, P. M. & Clements, J. E. SIV replication is directly downregulated by four antiviral miRNAs. Retrovirology, 10, 95 (2013).

29. Harris-Arnold, A., Arnold, C. P., Schaffert, S., Hatton, O., Krams, S. M., Esquivel, C. O. & Martinez, O. M. Epstein-Barr virus modulates host cell microRNA-194 to promote IL-10 production and B lymphoma cell survival. American Journal of Transplantation, 15, 2814–2824 (2015).

30. Dehghani, A., Khajepour, F., Dehghani, M., Razmara, E., Zangouey, M., Abadi, M. F. S., Nezhad, R. B. A., Dabiri, S. & Garshasbi, M. Hsa-miR-194-5p and hsa-miR-195-5p are down-regulated expressed in high dysplasia HPV-positive Pap smear samples compared to normal cytology HPV-positive Pap smear samples. BMC Infectious Diseases, 24, 182 (2024).

31. Kumar, M. & Nerurkar, V. R. Integrated analysis of microRNAs and their disease related targets in the brain of mice infected with West Nile virus. Virology, 452**-**453, 143-151 (2014).

32. Lv, J., Zhang, Z., Pan, L. & Zhang, Y. MicroRNA-34/449 family and viral infections. Virus Research, 260, 1–6 (2019).

33. Arvin, A., Campadelli-Fiume, G., Mocarski, E., Moore, P. S., Roizman, B., Whitley, R. & Yamanishi, K. (Eds.). Human Herpesviruses: Biology, Therapy, and Immunoprophylaxis. Cambridge University Pres (2007).

34. Agarwal, V., Bell, G. W., Nam, J. W. & Bartel, D. P. Predicting effective microRNA target sites in mammalian mRNAs. eLife, 4, e05005 (2015).

35. Chandriani, S. & Ganem, D. Host transcript accumulation during lytic KSHV infection reveals several classes of host responses. PLoS One, 2, e811 (2007).

36. Glaunsinger, B. & Ganem, D. Highly selective escape from KSHV-mediated host mRNA shutoff and its implications for viral pathogenesis. The Journal of Experimental Medicine, 200, 391–398 (2004).

37. Glaunsinger, B. & Ganem, D. Lytic KSHV infection inhibits host gene expression by accelerating global mRNA turnover. Molecular Cell, 13, 713–723 (2004).

38. Gabaev, I., Williamson, J. C., Crozier, T. W. M., Schulz, T. F. & Lehner, P. J. Quantitative Proteomics Analysis of Lytic KSHV Infection in Human Endothelial Cells Reveals Targets of Viral Immune Modulation. Cell Reports, 33, 108249 (2020).

39. Papp, B., Motlagh, N., Smindak, R. J., Jin Jang, S., Sharma, A., Alonso, J. D. & Toth, Z. Genome-Wide Identification of Direct RTA Targets Reveals Key Host Factors for Kaposi’s Sarcoma-Associated Herpesvirus Lytic Reactivation. Journal of Virology, 93, e01978–18 (2019).

40. Enright, A. J., John, B., Gaul, U., Tuschl, T., Sander, C. & Marks, D. S. MicroRNA targets in Drosophila. Genome Biology, 5, R1 (2003).

41. McClure, L. V., Kincaid, R. P., Burke, J. M., Grundhoff, A. & Sullivan, C. S. Comprehensive mapping and analysis of Kaposi’s sarcoma-associated herpesvirus 3’ UTRs identify differential posttranscriptional control of gene expression in lytic versus latent infection. Journal of Virology, 87, 12838–12849 (2013).

42. Vernet, C. & Artzt, K. STAR, a gene family involved in signal transduction and activation of RNA. Trends in Genetics, 13, 479–484 (1997).

43. Lukong, K. E. & Richard, S. Sam68, the KH domain-containing superSTAR. Biochimica et Biophysica Acta (BBA) - Reviews on Cancer, 1653, 73–86 (2003).

44. Feracci, M., Foot, J. N., Grellscheid, S. N., Danilenko, M., Stehle, R., Gonchar, O., Kang, H. S., Dalgliesh, C., Meyer, N. H., Liu, Y., Lahat, A., Sattler, M., Eperon, I. C., Elliott, D. & Dominguez, C. Structural basis of RNA recognition and dimerization by the STAR proteins T-STAR and Sam68. Nature Communications, 7, 10355 (2016).

45. Wakeman, B. S., Izumiya, Y. & Speck, S. H. Identification of Novel Kaposi’s Sarcoma Associated Herpesvirus Orf50 Transcripts: Discovery of New RTA Isoforms with Variable Transactivation Potential. Journal of Virology, 91, e01434–16 (2017).

46. Majerciak, V. & Zheng, Z. M. Alternative RNA splicing of KSHV ORF57 produces two different RNA isoforms. Virology, 488, 81–87 (2016).

47. Tang, S. & Zheng, Z. M. Kaposi’s sarcoma-associated herpesvirus K8 exon 3 contains three 5’-splice sites and harbors a K8.1 transcription start site. Journal of Biological Chemistry, 277, 14547–14556 (2002).

48. Burd, C. G. & Dreyfuss, G. Conserved structures and diversity of functions of RNA-binding proteins. Science, 265, 615–621 (1994).

49. Dreyfuss, G., Kim, V. N. & Kataoka, N. Messenger-RNA-binding proteins and the messages they carry. Nature Reviews Molecular Cell Biology, 3, 195–205 (2002).

50. Geuens, T., Bouhy, D. & Timmerman, V. The hnRNP family: insights into their role in health and disease. Human Genetics, 135, 851–867 (2016).

51. Stepicheva, N. A. & Song, J. L. Function and regulation of microRNA-31 in development and disease. Molecular Reproduction & Development, 83, 654–674 (2016).

52. Yu, T., Ma, P., Wu, D., Shu, Y. & Gao, W. Functions and mechanisms of microRNA-31 in human cancers. Biomedicine & Pharmacotherapy, 108, 1162–1169 (2018).

53. Fu, X. D. & Ares, Jr., M. Context-dependent control of alternative splicing by RNA-binding proteins. Nature Reviews Genetics, 15, 689–701 (2014).

54. Martin, K. C. & Ephrussi, A. mRNA localization: gene expression in the spatial dimension. Cell, 136, 719–730 (2009).

55. Moore, M. J. & Proudfoot, N. J. Pre-mRNA processing reaches back to transcription and ahead of translation. Cell, 136, 688–700 (2009).

56. Sonenberg, N. & Hinnebusch, A. G. Regulation of translation initiation in eukaryotes: mechanisms and biological targets. Cell, 136, 731–745 (2009).

57. Dassi, E. Handshakes and Fights: The Regulatory Interplay of RNA-Binding Proteins. Frontiers in Molecular Biosciences, 4, 67 (2017).

58. Hirose, Y. & Manley, J. L. RNA polymerase II and the integration of nuclear events. Genes & Development, 14, 1415–1429 (2000).

59. Maniatis, T. & Reed, R. An extensive network of coupling among gene expression machines. Nature, 416, 499–506 (2002).

60. Proudfoot, N. J., Furger, A. & Dye, M. J. Integrating mRNA processing with transcription. Cell, 108, 501–512 (2002).

61. Kornblihtt, A. R., Mata, M. D. L., Fededa, J. P., Munoz, M. J. & Nogues, G. Multiple links between transcription and splicing. RNA, 10, 1489–1498 (2004).

62. Modem, S., Badri, K. R., Holland, T. C. & Reddy, T. R. Sam68 is absolutely required for Rev function and HIV-1 production. Nucleic Acids Research, 33, 873–879 (2005).

63. Suhasini, M. & Reddy, T. R. Cellular proteins and HIV-1 Rev function. Current HIV Research, 7, 91–100 (2009).

64. Batra, R., Stark, T. J., Clark, A. E., Belzile, J. P., Wheeler, E. C., Yee, B. A., Huang, H., Gelboin Burkhart, C., Huelga, S. C., Aigner, S., Roberts, B. T., Bos, T. J., Sathe, S., Donohue, J. P., Rigo, F., Ares, M., Jr, Spector, D. H. & Yeo, G. W. RNA-binding protein CPEB1 remodels host and viral RNA landscapes. Nature Structural & Molecular Biology, 23, 1101–1110 (2016).

65. Kajitani, N. & Schwartz, S. The role of RNA-binding proteins in the processing of mRNAs produced by carcinogenic papillomaviruses. Seminars in Cancer Biology, 86, 482–496 (2022).

66. Jayabalan, A. K., Griffin, D. E. & Leung, A. K. L. Pro-Viral and Anti-Viral Roles of the RNA-Binding Protein G3BP1. Viruses, 15, 449 (2023).

67. Reddy, T. R., Xu, W. D. & Wong-Staal, F. General effect of Sam68 on Rev/Rex regulated expression of complex retroviruses. Oncogene, 19, 4071–4074 (2000).

68. Wegener, M. & Müller-McNicoll, M. Nuclear retention of mRNAs - quality control, gene regulation and human disease. Seminars in Cell & Developmental Biology, 79, 131–142 (2018).

69. Gordon, J. M., Phizicky, D. V. & Neugebauer, K. M. Nuclear mechanisms of gene expression control: pre-mRNA splicing as a life or death decision. Current Opinion in Genetics & Development, 67, 67–76 (2021).

70. Emery, A. & Swanstrom, R. HIV-1: To Splice or Not to Splice, That Is the Question. Viruses, 13, 181 (2021).

71. Toro-Ascuy, D., Rojas-Araya, B., Valiente-Echeverría, F. & Soto-Rifo, R. Interactions between the HIV-1 Unspliced mRNA and Host mRNA Decay Machineries. Viruses, 8, 320 (2016).

72. Reddy, T. R., Xu, W., Mau, J. K., Goodwin, C. D., Suhasini, M., Tang, H., Frimpong, K., Rose, D. W. & Wong-Staal, F. Inhibition of HIV replication by dominant negative mutants of Sam68, a functional homolog of HIV-1 Rev. Nature Medicine, 5, 635–642 (1999).

73. Derry, J. J., Richard, S., Carvajal, H. V., Ye, X., Vasioukhin, V., Cochrane, A. W., Chen, T. & Tyner, A. L. Sik (BRK) phosphorylates Sam68 in the nucleus and negatively regulates its RNA binding activity. Molecular and Cellular Biology, 20, 6114–6126 (2000).

74. Chen, T., Côté, J., Carvajal, H. V. & Richard, S. Identification of Sam68 arginine glycine-rich sequences capable of conferring nonspecific RNA binding to the GSG domain. Journal of Biological Chemistry, 276, 30803–30811 (2001).

75. McLaren, M., Asai, K. & Cochrane, A. A novel function for sam68: enhancement of HIV-1 RNA 3’ end processing. RNA, 10, 1119–1129 (2004).

76. Dodon, M. D., Mikaélian, I., Sergeant, A. & Gazzolo, L. The herpes simplex virus 1 Us11 protein cooperates with suboptimal amounts of human immunodeficiency virus type 1 (HIV-1) Rev protein to rescue HIV-1 production. Virology, 270, 43–53 (2000).

77. Diaz, J. J., Dodon, M. D., Schaerer-Uthurralt, N., Simonin, D., Kindbeiter, K., Gazzolo, L. & Madjar, J. J. Post-transcriptional transactivation of human retroviral envelope glycoprotein expression by herpes simplex virus Us11 protein. Nature, 379, 273–277 (1996).

78. Zheng, Z. M. Split genes and their expression in Kaposi’s sarcoma-associated herpesvirus. Reviews in Medical Virology, 13, 173–184 (2003).

79. Ajiro, M. & Zheng, Z. M. Oncogenes and RNA splicing of human tumor viruses. Emerging Microbes & Infections, 3, e63 (2014).

80. Arias, C., Weisburd, B., Stern-Ginossar, N., Mercier, A., Madrid, A. S., Bellare, P., Holdorf, M., Weissman, J. S. & Ganem, D. KSHV 2.0: a comprehensive annotation of the Kaposi’s sarcoma-associated herpesvirus genome using next-generation sequencing reveals novel genomic and functional features. PLoS Pathogens, 10, e1003847 (2014).

81. Bruce, A. G., Barcy, S., DiMaio, T., Gan, E., Garrigues, H. J., Lagunoff, M. & Rose, T. M. Quantitative Analysis of the KSHV Transcriptome Following Primary Infection of Blood and Lymphatic Endothelial Cells. Pathogens, 6, 11 (2017).

82. Zhao, Y., Ye, X., Shehata, M., Dunker, W., Xie, Z. & Karijolich, J. The RNA quality control pathway nonsense-mediated mRNA decay targets cellular and viral RNAs to restrict KSHV. Nature Communications, 11, 3345> (2020).

83. Majerciak, V., Alvarado-Hernandez, B., Lobanov, A., Cam, M. & Zheng, Z. M. Genome-wide regulation of KSHV RNA splicing by viral RNA-binding protein ORF57. PLoS Pathogens, 18, e1010311 (2022).

84. Xiao, R., Chen, J. Y., Liang, Z., Luo, D., Chen, G., Lu, Z. J., Chen, Y., Zhou, B., Li, H., Du, X., Yang, Y., San, M., Wei, X., Liu, W., Lécuyer, E., Graveley, B. R., Yeo, G. W., Burge, C. B., Zhang, M. Q., Zhou, Y. & Fu, X. D. Pervasive Chromatin-RNA Binding Protein Interactions Enable RNA-Based Regulation of Transcription. Cell, 178, 107–121.e18 (2019).

85. Lidenge, S. J., Kossenkov, A. V., Tso, F. Y., Wickramasinghe, J., Privatt, S. R., Ngalamika, O., Ngowi, J. R., Mwaiselage, J., Lieberman, P. M., West, J. T. & Wood, C. Comparative transcriptome analysis of endemic and epidemic Kaposi’s sarcoma (KS) lesions and the secondary role of HIV-1 in KS pathogenesis. PLoS Pathogens, 16, e1008681 (2020).

86. Qin, H., Ni, H., Liu, Y., Yuan, Y., Xi, T., Li, X. & Zheng, L. RNA-binding proteins in tumor progression. Journal of Hematology & Oncology, 13, 90 (2020).

87. Pereira, B., Billaud, M. & Almeida, R. RNA-Binding Proteins in Cancer: Old Players and New Actors. Trends in Cancer, 3, 506–528 (2017).

88. Kang, D., Lee, Y. & Lee, J. S. RNA-Binding Proteins in Cancer: Functional and Therapeutic Perspectives. Cancers (Basel), 12, 2699 (2020).

89. Kechavarzi, B. & Janga, S. C. Dissecting the expression landscape of RNA-binding proteins in human cancers. Genome Biology, 15, R14 (2014).

90. Frisone, P., Pradella, D., Matteo, A. D., Belloni, E., Ghigna, C. & Paronetto, M. P. SAM68: Signal Transduction and RNA Metabolism in Human Cancer. BioMed Research International, 2015, 528954 (2015).

91. Vidigal, J. A. & Ventura, A. The biological functions of miRNAs: lessons from in vivo studies. Trends in Cell Biology, 25, 137–147 (2015).

92. Speck, S. H. & Ganem, D. Viral latency and its regulation: lessons from the gamma-herpesviruses. Cell Host & Microbe, 8, 100–115 (2010).

93. Broussard, G. & Damania, B. Regulation of KSHV Latency and Lytic Reactivation. Viruses, 12, 1034 (2020).

94. Gaucherand, L. & Gaglia, M. M. The Role of Viral RNA Degrading Factors in Shutoff of Host Gene Expression. Annual Review of Virology, 9, 213–238 (2022).

95. Love, M. I., Huber, W. & Anders, S. Moderated estimation of fold change and dispersion for RNA-seq data with DESeq2. Genome Biology, 15, 550 (2014).

96. Langmead, B. & Salzberg, S. L. Fast gapped-read alignment with Bowtie 2. Nature Methods, 9, 357–359 (2012).

97. Li, B. & Dewey, C. N. RSEM: accurate transcript quantification from RNA-Seq data with or without a reference genome. BMC Bioinformatics, 12, 323 (2011).

98. Robinson, M. D., McCarthy, D. J. & Smyth, G. K. edgeR: a Bioconductor package for differential expression analysis of digital gene expression data. Bioinformatics, 26, 139–140 (2010).

99. Luo, W., Friedman, M. S., Shedden, K., Hankenson, K. D. & Woolf, P. J. GAGE: generally applicable gene set enrichment for pathway analysis. BMC Bioinformatics, 10, 161 (2009).

100. Robinson, M. D. & Oshlack, A. A scaling normalization method for differential expression analysis of RNA-seq data. Genome Biology, 11, R25 (2010).

## SI References

1. Lidenge, S. J., Kossenkov, A. V., Tso, F. Y., Wickramasinghe, J., Privatt, S. R., Ngalamika, O., Ngowi, J. R., Mwaiselage, J., Lieberman, P. M., West, J. T. & Wood, C. Comparative transcriptome analysis of endemic and epidemic Kaposi’s sarcoma (KS) lesions and the secondary role of HIV-1 in KS pathogenesis. PLoS Pathogens, 16, e1008681 (2020).

2. Myoung, J. & Ganem, D. Generation of a doxycycline-inducible KSHV producer cell line of endothelial origin: maintenance of tight latency with efficient reactivation upon induction. Journal of Virological Methods, 174, 12–21 (2011).

3. An, F. Q., Folarin, H. M., Compitello, N., Roth, J., Gerson, S. L., McCrae K.R., Fakhari, F. D., Dittmer, D. P. & Renne, R. Long-Term-Infected Telomerase-Immortalized Endothelial Cells: a Model for Kaposi’s Sarcoma-Associated Herpesvirus Latency In Vitro and In Vivo. Journal of Virology, 80, 4833–4846 (2006).

4. Love, M. I., Huber, W. & Anders, S. Moderated estimation of fold change and dispersion for RNA-seq data with DESeq2. Genome Biology, 15, 550 (2014).

5. Robinson, M. D., McCarthy, D. J. & Smyth, G. K. edgeR: a Bioconductor package for differential expression analysis of digital gene expression data. Bioinformatics, 26, 139–140 (2010).

6. Langmead, B. & Salzberg, S. L. Fast gapped-read alignment with Bowtie 2. Nature Methods, 9, 357–359 (2012).

7. Li, B. & Dewey, C. N. RSEM: accurate transcript quantification from RNA-Seq data with or without a reference genome. BMC Bioinformatics, 12, 323 (2011).

8. Luo, W., Friedman, M. S., Shedden, K., Hankenson, K. D. & Woolf, P. J. GAGE: generally applicable gene set enrichment for pathway analysis. BMC Bioinformatics, 10, 161 (2009).

9. Agarwal, V., Bell, G. W., Nam, J. W. & Bartel, D. P. Predicting effective microRNA target sites in mammalian mRNAs. eLife, 4, e05005 (2015).

