## Supplemental Data for "Host microRNA-31-5p represses oncogenic herpesvirus lytic reactivation by restricting the RNA-binding protein KHDRBS3-mediated viral gene expression"

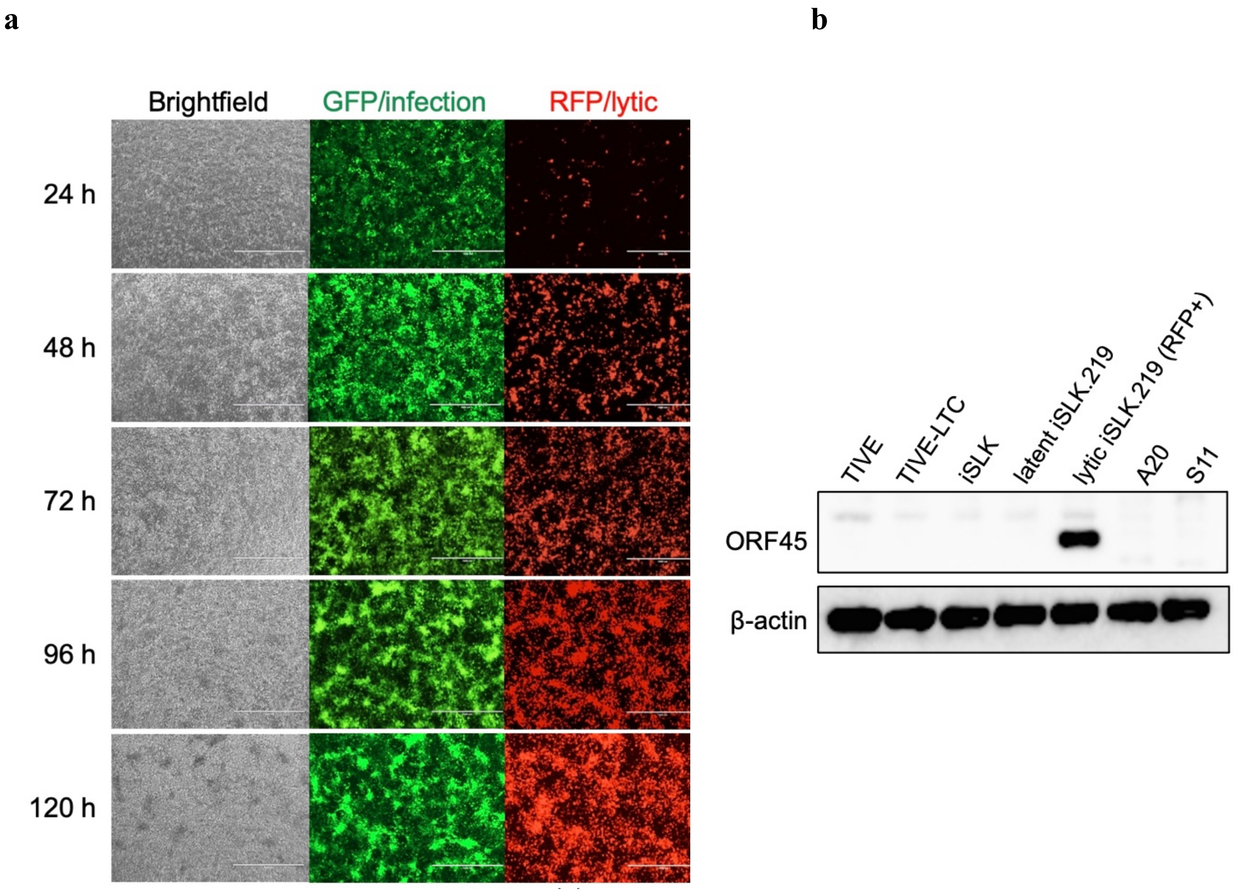


**Supplementary Figure 1. iSLK.219 cells with dox-inducible RTA expression undergo efficient lytic reactivation upon Dox treatment.** (**a**) Representative images showing phase, GFP and RFP fluorescence of iSLK.219 cells over a 120 h Dox-induced lytic reactivation time course, with samples collected at 24, 48, 72, 96, and 120 h. iSLK.219 cells harbor a recombinant KSHV (rKSHV.219) genome with constitutive GFP expression to indicate latent infection, and induced RFP expression upon lytic reactivation to indicate lytic infection. (**b**) Western blot analysis of immediate-early KSHV lytic protein ORF45 expression in the indicated cell lines, including latent iSLK.219 cells at 0 h and lytic iSLK.219 cells at 72 h post Dox-induced KSHV lytic reactivation. β-actin was used as a loading control.

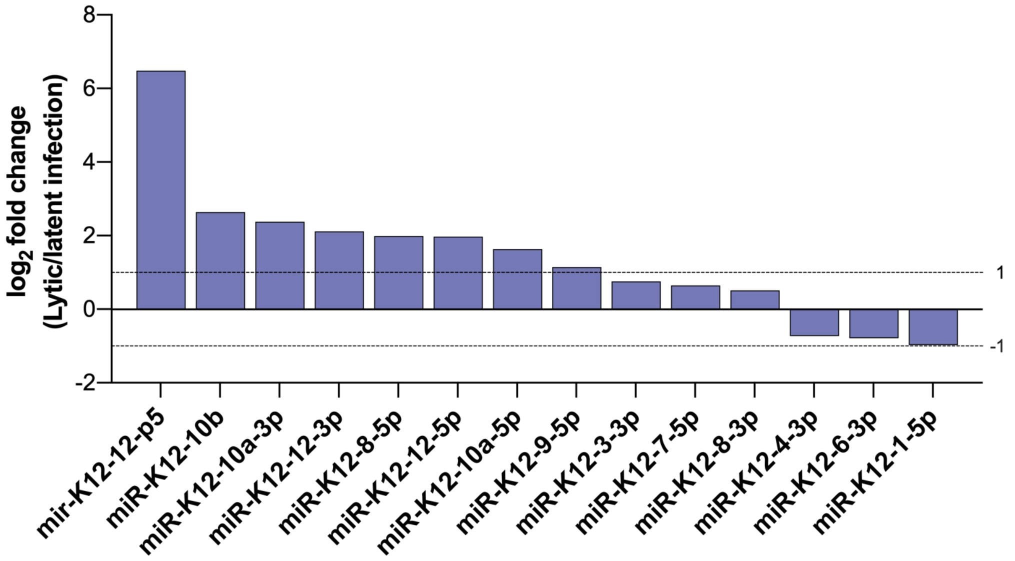


**Supplementary Figure 2. Changes in KSHV miRNA expression in iSLK.219 cells during KSHV lytic reactivation, as determined by small-RNA sequencing**. Waterfall plot depicting KSHV-encoded miRNAs with a $\log_2$ fold change value $\geq |0.5|$ in iSLK.219 cells with lytic (Dox+) compared to latent (Dox-) KSHV infection. Total RNA isolated from latent iSLK.219 cells at 0 h or lytic iSLK.219 cells at 72 h post Dox-induced KSHV lytic reactivation was subjected to small-RNA sequencing analysis. mir-K12-12-p5 is the miR-K12-12 precursor (pre-miR-K12-12).

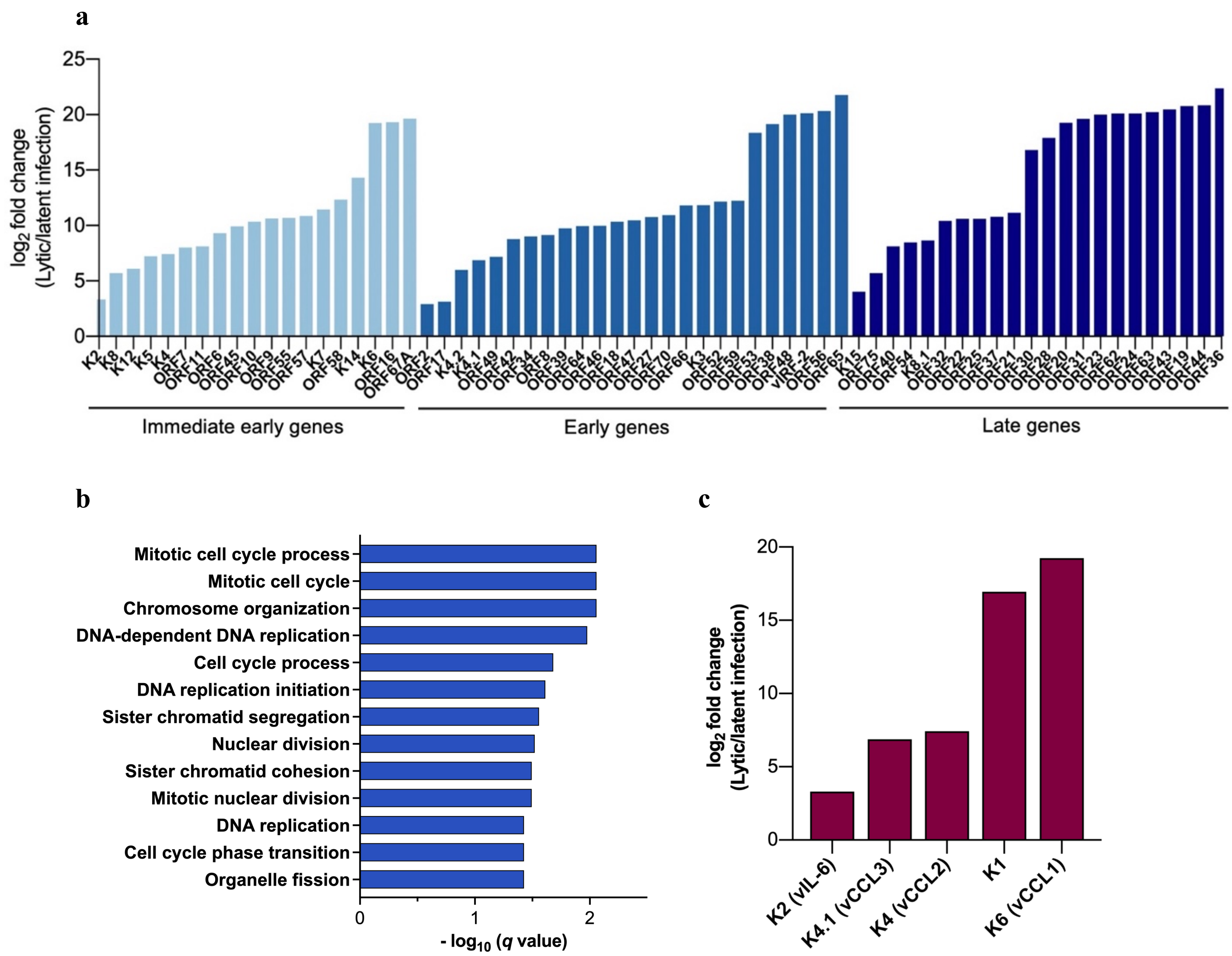


**Supplementary Figure 3. Changes in viral and cellular mRNA expression in iSLK.219 cells during KSHV lytic reactivation, as determined by RNA sequencing.** (a) RNA sequencing analysis of 66 KSHV open reading frames (ORFs) in iSLK.219 cells with lytic (Dox+) compared to latent (Dox-) KSHV infection. (**b**) Gene ontology (GO) analysis was performed to analyze all genes significantly up-regulated in lytic iSLK.219 cells (Dox+) compared to latent iSLK.219 cells (Dox-) at 72 h post Dox-induced lytic reactivation, as determined by RNA-seq, identifying significantly enriched GO terms. (**c**) Expression of KSHV lytic genes with mitogenic signaling activities, K2 (vIL-6), K4.1 (vCCL3), K4 (vCCL2), K1, and K6 (vCCL1), in iSLK.219 cells with lytic (Dox+) compared to latent (Dox-) KSHV infection. Total RNA was isolated from latent iSLK.219 cells at 0 h or lytic iSLK.219 cells at 72 h post Dox-induced KSHV lytic reactivation was subjected to RNA sequencing analysis.

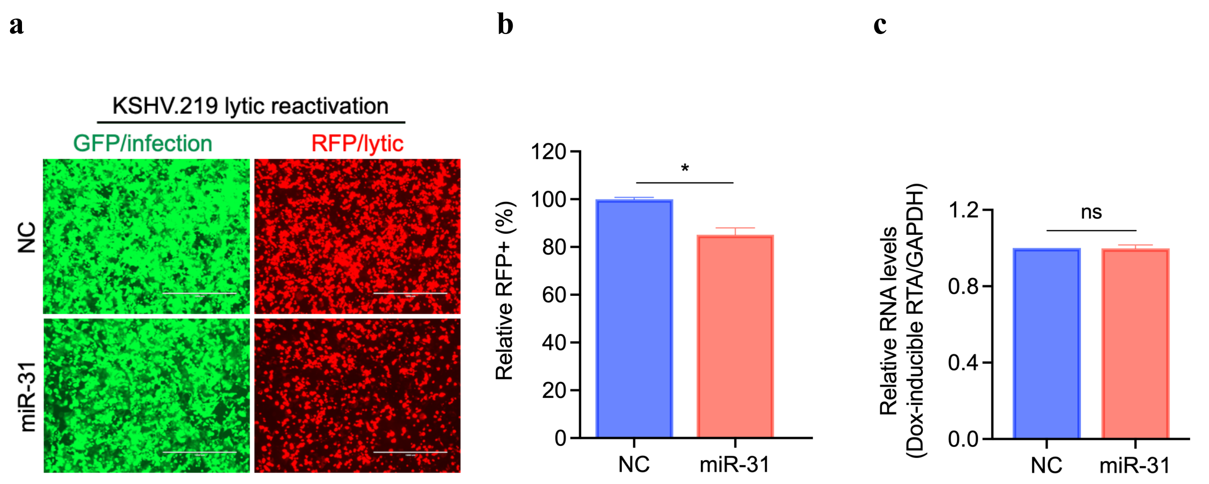


**Supplementary Figure 4. Ectopic miR-31 expression suppresses KSHV lytic reactivation.** (**a**) iSLK.219 cells were transfected with negative control (NC) or miR-31 mimic followed by treatment with Dox (0.5 μg/mL) for 72 h to induce KHSV lytic reactivation, and representative images were photographed for GFP and RFP fluorescence at 72 h post Dox-induced lytic reactivation (Scale bars, 1000 μm). (**b**) Quantification of RFP-positive cells in (A) using image cytometry. Two biological replicates are presented. (**c**) iSLK cells were transfected with negative control (NC) or miR-31 mimic followed by treatment with Dox (0.5 μg/mL) for 72 h. Total RNA was isolated at 72 h post-Dox treatment and expression of Dox-induced *RTA* was quantified by RT-qPCR. *GAPDH* was used as an endogenous control. Two biological replicates are presented. Data represent mean ± SEM and *p*-values were determined by two-tailed Student's *t*-test. $^{*}p < 0.05$, $^{**}p < 0.01$, $^{***}p < 0.001$, $^{****}p < 0.0001$, and ns = not significant.

a

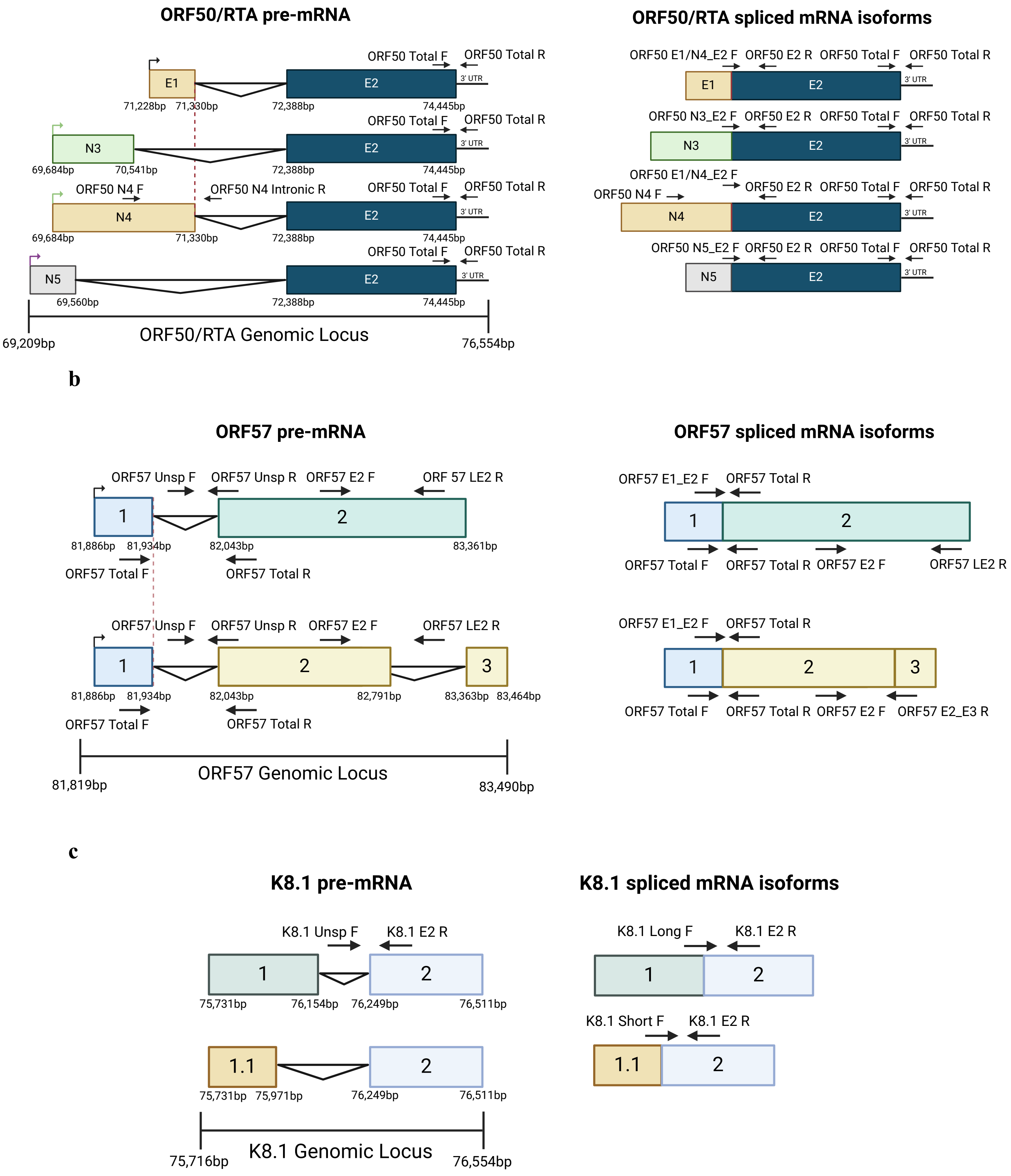
ORF50/RTA pre-mRNA
ORF50 Total F
ORF50 Total R
E1
E2
3' UTR
71,228bp
71,330bp
72,388bp
74,445bp
N3
69,684bp
70,541bp
ORF50 N4 F
ORF50 N4 Intronic R
N4
71,330bp
N5
69,560bp
ORF50/RTA Genomic Locus
69,209bp
76,554bp
ORF50/RTA spliced mRNA isoforms
ORF50 E1/N4_E2 F
ORF50 E2 R
ORF50 N3_E2 F
ORF50 N5_E2 F
b
ORF57 pre-mRNA
ORF57 Unsp F
ORF57 Unsp R
ORF57 E2 F
ORF 57 LE2 R
ORF57 LE2 R
81,886bp
81,934bp
82,043bp
83,361bp
ORF57 Total F
ORF57 Total R
82,791bp
83,363bp
83,464bp
ORF57 Genomic Locus
81,819bp
83,490bp
ORF57 spliced mRNA isoforms
ORF57 E1_E2 F
ORF57 E2_E3 R
c
K8.1 pre-mRNA
K8.1 Unsp F
K8.1 E2 R
75,731bp
76,154bp
76,249bp
76,511bp
1.1
75,971bp
K8.1 Genomic Locus
75,716bp
76,554bp
K8.1 spliced mRNA isoforms
K8.1 Long F
K8.1 Short F

**Supplementary Figure 5. Analysis of alternative splicing in the immediate-early lytic ORF50, early lytic ORF57, and late lytic K8.1 KSHV genes.** (**a-c**) Schematic representation of alternatively spliced transcripts of *ORF50* (**a**), *ORF57* (**b**), and *K8.1* (**c**) in the context of the corresponding gene, focusing on alternatively spliced exons. The region of mRNA that demonstrates the alternative splicing events is depicted in greater detail. Colored boxes represent exons and the horizontal black lines represent introns. The arrowheads depict the approximate RT-qPCR primer location. PCR primer sequences are provided in Supplementary Table 6.

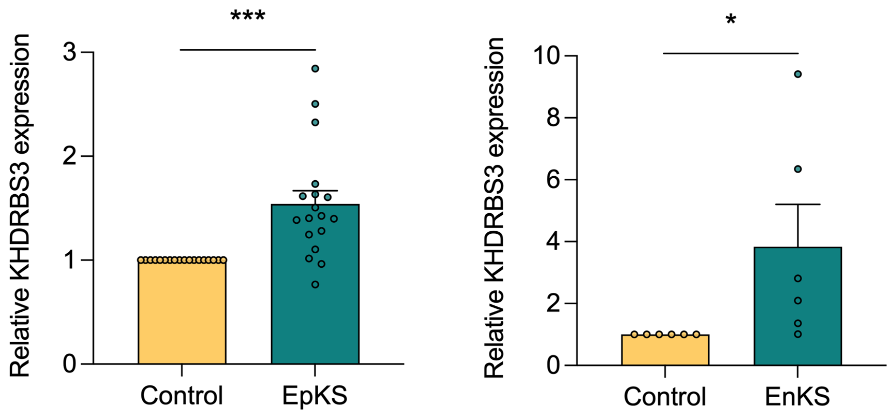


**Supplementary Figure 6. RNA-seq analysis of KHDRBS3 and miR-31-5p expression in KS lesions.** *KHDRBS3* expression in 18 pairs of AIDS-associated/epidemic KS (EpKS) (Left) or 6 pairs of non-AIDS associated/endemic KS (EnKS) (Right), relative to matched uninvolved control tissues from the published RNA-seq data[1]. Data represent mean ± SEM and $p$-values were determined by two-tailed Student's $t$-test. *$p < 0.05$, **$p < 0.01$, ***$p < 0.001$, ****$p < 0.0001$, and ns = not significant.

**Supplementary Table 1.** Reagent sources

| REAGENT or RESOURCE | SOURCE | IDENTIFIER |
|---|---|---|
| **Antibodies** | | |
| Mouse monoclonal anti-β-Actin | Santa Cruz Technology | Cat#sc-47778; RRID:AB_2714189 |
| Mouse monoclonal anti-KSHV ORF26 | NOVUS | Cat#NBP1-47357 |
| Mouse monoclonal anti-KSHV K8 | NOVUS | Cat#NB100-2189 |
| Mouse monoclonal anti-KSHV ORF45 | NOVUS | Cat#NBP2-37685 |
| Rabbit polyclonal anti-KSHV ORF50 | Abbiotec | Cat#251345 |
| Goat anti-rabbit IgG, HRP-linked | Cell Signaling Technology | Cat#7074; RRID:AB_2099233 |
| Horse anti-mouse IgG, HRP-linked | Cell Signaling Technology | Cat#7076; RRID:AB_330924 |
| **Bacterial and Virus Strains** | | |
| Stable Competent *E. coli* (High Efficiency) | New England Biolabs | Cat#C3040H |
| 5-alpha Competent *E. coli* (High Efficiency) | New England Biolabs | Cat#C2987H |
| rKSHV.219 | iSLK.219 cells[2] | |
| **Chemicals, Peptides, and Recombinant Proteins** | | |
| DMEM | Gibco | Cat#11995065 |
| Tet System Approved Fetal Bovine Serum | Takara Bio | Cat#631101 |
| Lipofectamine RNAiMAX Transfection Reagent | Invitrogen | Cat#13778500 |
| Lipofectamine 3000 Transfection Reagent | Invitrogen | Cat#L3000008 |
| RIPA Buffer | Boston BioProducts | Cat#BP-115 |
| Phosphatase Inhibitor Cocktail 2 | Sigma-Aldrich | Cat#P5726 |
| Phosphatase Inhibitor Cocktail 3 | Sigma-Aldrich | Cat#P0044 |
| Halt Protease Inhibitor Cocktail | Thermo Scientific | Cat#78430 |
| Halt Protease and Phosphatase Inhibitor Cocktail | Thermo Scientific | Cat#78440 |
| Clarity Western ECL Substrate | BIO-RAD | Cat#1705061 |
| Doxycycline Hydrochloride | Sigma-Aldrich | Cat#D3072 |
| 4x Laemmli Sample Buffer | BIO-RAD | Cat#1610747 |
| Nitrocellulose Membrane | BIO-RAD | Cat#1620146 |
| Sucrose solution | Millipore | Cat#8590 |
| DNAzol® | Molecular research center | Cat#DN127 |
| Puromycin Dihydrochloride | Gibco | Cat#A1113803 |
| Hygromycin B | InvivoGen | Cat#ant-hg-1 |
| Geneticin (G418 Sulfate) | Gibco | Cat#10131027 |
| *NotI-HF* | New England Biolabs | Cat#R3189L |
| *XhoI* | New England Biolabs | Cat#R0146S |
| **Critical Commercial Assays** | | |
| Dual-Luciferase Reporter Assay | Promega | Cat#E1910 |
| LightCycler 480 SYBR Green I Master | Roche | Cat#04707516001 |
| SuperScript IV Reverse Transcriptase | Invitrogen | Cat#18090010 |
| Zero Blunt TOPO PCR Cloning Kit | Invitrogen | Cat#K2800J10 |

| | | |
|---|---|---|
| QuikChange Lightening Site-Directed Mutagenesis Kit | Agilent Technologies | Cat#210518 |
| *mir*Vana miRNA Isolation Kit | Invitrogen | Cat#AM1560 |
| Pierce BCA Protein Assay Kit | Thermo Scientific | Cat#23225 |
| MycoAlert Mycoplasma Detection Kit | Lonza | Cat#LT07-318 |
| **Experimental Models: Cell Lines** | | |
| iSLK cell line[2] | Laboratory of Kenneth Kaye, Harvard Medical School | N/A |
| iSLK.219 cell line[2] | Laboratory of Kenneth Kaye, Harvard Medical School | N/A |
| TIVE cell line[3] | Laboratory of Rolf Renne, University of Florida | N/A |
| TIVE-LTC cell line[3] | Laboratory of Rolf Renne, University of Florida | N/A |
| **Oligonucleotides** | | |
| *mir*Vana MiRNA mimic, hsa-miR-31-5p, assay ID: MC11465 | Invitrogen | Cat#4464066 |
| *mir*Vana MiRNA mimic, Negative Control No. 1 | Invitrogen | Cat#4464059 |
| Silencer Select siRNA KHDRBS3 #1, assay ID: s20950 | Thermo Scientific | Cat#4392420 |
| Silencer Select siRNA KHDRBS3 #2, assay ID: s20949 | Thermo Scientific | Cat#4392420 |
| Silencer Select siRNA SLC51A #1, assay ID: s225837 | Thermo Scientific | Cat#4392420 |
| Silencer Select siRNA SLC51A #2, assay ID: s47293 | Thermo Scientific | Cat#4392420 |
| *Silencer*™ Select Negative Control No. 1 siRNA | Thermo Scientific | Cat#4390843 |
| Hs_GAPDH_1_SG QuantiTect Primer Assay QT00079247 | Qiagen | Cat#249900 |
| Hs_CD109_1_SG QuantiTect Primer Assay QT00064589 | Qiagen | Cat#249900 |
| Hs_CKAP4_1_SG QuantiTect Primer Assay QT00203245 | Qiagen | Cat#249900 |
| Hs_IFIT1_1_SG QuantiTect Primer Assay QT00201012 | Qiagen | Cat#249900 |
| Hs_KHDRBS3_1_SG QuantiTect Primer Assay QT00065296 | Qiagen | Cat#249900 |
| Hs_NNMT_1_SG QuantiTect Primer Assay QT00076041 | Qiagen | Cat#249900 |
| Hs_SAR1A_1_SG QuantiTect Primer Assay QT00082173 | Qiagen | Cat#249900 |
| Hs_SSH1_1_SG QuantiTect Primer Assay QT00011417 | Qiagen | Cat#249900 |
| Hs_STOM_1_SG QuantiTect Primer Assay QT00033152 | Qiagen | Cat#249900 |

| | | |
|---|---|---|
| Hs_TRIM59_1_SG QuantiTect Primer Assay QT00047166 | Qiagen | Cat#249900 |
| Hs_YPEL2_1_SG QuantiTect Primer Assay QT00069482 | Qiagen | Cat#249900 |
| Hs_AGT_1_SG QuantiTect Primer Assay QT00005614 | Qiagen | Cat#249900 |
| Hs_C15orf39_1_SG QuantiTect Primer Assay QT00200900 | Qiagen | Cat#249900 |
| Hs_C1orf198_1_SG QuantiTect Primer Assay QT02322908 | Qiagen | Cat#249900 |
| Hs_DTX4_1_SG QuantiTect Primer Assay QT00197484 | Qiagen | Cat#249900 |
| Hs_GSTM4_1_SG QuantiTect Primer Assay QT00046662 | Qiagen | Cat#249900 |
| Hs_PAQR8_1_SG QuantiTect Primer Assay QT01028104 | Qiagen | Cat#249900 |
| Hs_PLEKHO2_1_SG QuantiTect Primer Assay QT00069377 | Qiagen | Cat#249900 |
| Hs_PTGFR_1_SG QuantiTect Primer Assay QT00029540 | Qiagen | Cat#249900 |
| Hs_SEMA5A_1_SG QuantiTect Primer Assay QT00053417 | Qiagen | Cat#249900 |
| Hs_SLC51A_1_SG QuantiTect Primer Assay QT00493059 | Qiagen | Cat#249900 |
| Hs_CRISPLD2_1_SG QuantiTect Primer Assay QT00072485 | Qiagen | Cat#249900 |
| qPCR primers | | Supplementary Table 6 |
| Mutagenic primers | | Supplementary Table 6 |
| Primers for cloning 3' UTRs of the indicated genes into psiCHECK-2 vector | | Supplementary Table 6 |
| **Software and Algorithms** | | |
| TargetScanHuman (version 7.2, March 2018)[9] | Whitehead Institute for Biomedical Research | https://www.targetscan.org/vert_71/ |
| Fiji (version 2.0.0-rc-69/1.52i, December 2018) | http://imagej.net/Fiji | RRID:SCR_002285 |
| GraphPad Prism (version 8.4.3 (471), June 2020) | GraphPad Software | https://www.graphpad.com/; RRID:SCR_002798 |
| miRBase (v22) | Griffiths-Jones Lab, University of Manchester | https://www.mirbase.org/; RRID:SCR_017497 |
| Biorender | N/A | https://www.biorender.com |
| FastQC | Simon Andrews | https://www.bioinformatics.babraham.ac.uk/projects/fastqc/; RRID:SCR_014583 |
| DESeq2 (v1.26.0)[4] | | https://bioconductor.org/packages/release/bioc/html/DESeq2.html; RRID:SCR_015687 |

| | | |
|---|---|---|
| EdgeR[5] | | https://bioconductor.org/packages/release/bioc/html/edgeR.html; RRID:SCR_012802 |
| Bowtie2 (v2.2.9)[6] | | http://bowtie-bio.sourceforge.net/bowtie2/index.shtml; RRID:SCR_005476 |
| RSEM (v1.3.0)[7] | | RRID:SCR_013027 |
| GAGE (v2.20.1)[8] | | RRID:SCR_017067 |

**Supplementary Table 2.** A list of differentially expressed cellular miRNAs identified by small-RNA sequencing analysis of iSLK.219 cells with lytic (Dox+) compared to latent (Dox-) KSHV infection ($log_2$ fold change $\geq |0.6|$ and adjusted p-value < 0.05). These miRNAs are ranked according to the $log_2$ fold change (lytic/latent) value.

| miRNA | $Log_2$ (lytic/latent) | latent iSLK.219 reads | lytic iSLK.219 reads |
|---|---|---|---|
| hsa-miR-449c-5p | -7.4 | 360 | 2 |
| hsa-miR-194-5p | -2.0 | 2,902 | 712 |
| hsa-miR-181a-3p | -1.2 | 3,730 | 1,589 |
| hsa-miR-29a-3p | -1.1 | 34,062 | 16,024 |
| hsa-miR-31-5p | -1.0 | 5,484 | 2,773 |
| hsa-miR-196b-5p | -0.9 | 2,038 | 1,099 |
| hsa-let-7i-5p | -0.9 | 283,235 | 153,588 |
| hsa-miR-20a-3p | -0.8 | 241 | 138 |
| hsa-miR-26b-3p | -0.7 | 98 | 59 |
| hsa-miR-140-5p | -0.7 | 2,159 | 1,360 |
| hsa-miR-450b-5p | -0.6 | 1,730 | 1,157 |
| hsa-miR-378a-5p | 0.8 | 126 | 213 |
| hsa-miR-92b-3p | 0.8 | 2,849 | 5,095 |
| hsa-miR-188-5p | 0.9 | 103 | 186 |
| hsa-miR-378a-3p | 0.9 | 11,766 | 21,386 |
| hsa-miR-424-5p | 0.9 | 5,871 | 11,221 |
| hsa-miR-503-5p | 1.0 | 1,823 | 3,723 |
| hsa-miR-152-3p | 1.1 | 2,927 | 6,294 |
| hsa-miR-326 | 1.1 | 70 | 150 |
| hsa-miR-181c-3p | 1.2 | 119 | 272 |
| hsa-miR-501-5p | 1.2 | 56 | 128 |
| hsa-miR-574-5p | 1.2 | 754 | 1,765 |
| hsa-miR-135b-5p | 1.3 | 60 | 147 |
| hsa-miR-502-5p | 1.3 | 8 | 19 |
| hsa-miR-101-3p | 1.3 | 17,512 | 44,274 |
| hsa-miR-181c-5p | 1.5 | 210 | 599 |
| hsa-miR-3065-3p | 1.7 | 16 | 51 |
| hsa-miR-342-3p | 1.7 | 672 | 2,167 |
| hsa-miR-29c-3p | 1.7 | 342 | 1,132 |
| hsa-miR-149-5p | 1.8 | 223 | 758 |
| hsa-miR-181d-5p | 1.8 | 269 | 949 |
| hsa-miR-3065-5p | 2.1 | 41 | 168 |
| hsa-miR-210-3p | 2.1 | 2,487 | 10,696 |
| hsa-miR-7-5p | 2.2 | 647 | 2,909 |
| hsa-miR-139-5p | 2.5 | 194 | 1,089 |

Columns indicate the miRNA name, $log_2$ fold change (lytic/latent) value, latent iSLK.219 reads, and lytic iSLK.219 reads.

**Supplementary Table 3.** A list of differentially expressed KSHV-encoded miRNAs identified by small-RNA sequencing analysis of iSLK.219 cells with lytic (Dox+) compared to latent (Dox-) KSHV infection. These KSHV-encoded miRNAs are ranked according to the $log_2$ fold change (lytic/latent) value. KSHV miRNAs in the small-RNA seq data that have less than 10 reads are removed to avoid uncertain low counts.

| **KSHV miRNA** | **$Log_2$ (lytic/latent)** | **latent iSLK.219 reads** | **lytic iSLK.219 reads** |
|---|---|---|---|
| **mir-K12-12-p5** | 6.48 | 1.1 | 96.4 |
| **miR-K12-10b** | 2.64 | 114.4 | 712.5 |
| **miR-K12-10a-3p** | 2.38 | 64,339.3 | 335,553.9 |
| **miR-K12-12-3p** | 2.12 | 3,806.6 | 16,507.9 |
| **miR-K-12-8-5p** | 1.99 | 45.3 | 179.7 |
| **miR-K-12-12-5p** | 1.97 | 18,866.2 | 73,663.3 |
| **miR-K12-10a-5p** | 1.64 | 91.7 | 286.3 |
| **miR-K12-9-5p** | 1.15 | 389.6 | 863.5 |
| **miR-K12-3-3p** | 0.76 | 114.4 | 193.6 |
| **miR-K12-7-5p** | 0.65 | 25.9 | 40.8 |
| **miR-K12-8-3p** | 0.51 | 2,389.6 | 3,400.3 |
| miR-K12-2-3p | 0.46 | 16.2 | 22.2 |
| miR-K12-9-3p | 0.33 | 99.3 | 125.1 |
| miR-K12-5-3p | 0.27 | 365.9 | 442.0 |
| miR-K12-6-5p | 0.22 | 382.1 | 444.7 |
| miR-K12-7-3p | 0.04 | 1,658.9 | 1,706.7 |
| miR-K12-4-5p | 0.04 | 2,134.9 | 2,187.5 |
| miR-K12-3-5p | -0.03 | 15,230.0 | 14,894.8 |
| miR-K12-2-5p | -0.06 | 1,500.2 | 1,438.9 |
| miR-K12-11-3p | -0.06 | 3,508.8 | 3,362.4 |
| **miR-K12-4-3p** | -0.73 | 3,132.1 | 1,882.7 |
| **miR-K12-6-3p** | -0.79 | 7,788.2 | 4,502.0 |
| **miR-K12-1-5p** | -0.97 | 6,120.7 | 3,121.5 |

Columns indicate the KSHV miRNA name, $log_2$ fold change (lytic/latent) value, latent iSLK.219 reads and lytic iSLK.219 reads. mir-K12-12-p5 is the miR-K12-12 precursor (pre-miR-K12-12). Bolded miRNAs indicate KSHV miRNAs with a $log_2$ fold change value $\geq |0.5|$ in iSLK.219 cells with lytic (Dox+) compared to latent (Dox-) KSHV infection.

**Supplementary Table 4.** A list of TargetScan predicted miR-31-5p targets with conserved and poorly conserved sites and the $log_2$ fold change in mRNA expression of these predicted targets in iSLK.219 cells with lytic (Dox+) compared to latent (Dox-) KSHV infection, as determined by RNA-seq. These predicted targets are ranked according to the $log_2$ fold change (lytic/latent) value. Genes in the RNA-seq data that have less than 10 reads in lytic samples are removed to avoid uncertain low counts.

| Gene | $Log_2$ (lytic/latent) | latent iSLK.219 reads | lytic iSLK.219 reads |
|---|---|---|---|
| CYB561A3 | 1.02 | 8.98 | 18.26 |
| STRADA | 1.02 | 19.2 | 39.06 |
| ATP5A1 | 1.02 | 142.93 | 290.35 |
| ATOX1 | 1.03 | 124.21 | 254.08 |
| UBA1 | 1.04 | 109.74 | 225.1 |
| UNC45A | 1.07 | 18.77 | 39.52 |
| KIAA1524 | 1.08 | 8.26 | 17.51 |
| **KHDRBS3** | 1.08 | 5.1 | 10.77 |
| JMJD8 | 1.08 | 36.85 | 77.79 |
| FAM127A | 1.09 | 71.24 | 151.75 |
| TPGS2 | 1.11 | 27.04 | 58.37 |
| ARHGEF10 | 1.13 | 25.93 | 56.84 |
| MGAT1 | 1.14 | 30.27 | 66.49 |
| DHCR24 | 1.16 | 63.9 | 143.05 |
| RCN2 | 1.17 | 35.62 | 80.35 |
| MYH9 | 1.18 | 250.97 | 566.78 |
| SPICE1 | 1.21 | 4.92 | 11.39 |
| **TRIM59** | 1.23 | 10.52 | 24.76 |
| ATP11C | 1.24 | 11.49 | 27.21 |
| **C1orf198** | 1.24 | 22.84 | 53.92 |
| PTPLAD1 | 1.24 | 38.95 | 92.06 |
| ATL3 | 1.25 | 8.39 | 19.94 |
| GEMIN8 | 1.26 | 8.12 | 19.43 |
| TAPBP | 1.27 | 57.81 | 139.7 |
| **SSH1** | 1.27 | 9.7 | 23.41 |
| DPAGT1 | 1.32 | 11.61 | 29.08 |
| WFS1 | 1.34 | 11.29 | 28.67 |
| DDX54 | 1.34 | 42.21 | 107.15 |
| FAM162A | 1.39 | 19.98 | 52.46 |
| SNTB2 | 1.39 | 4.1 | 10.77 |
| CEP85 | 1.39 | 4.75 | 12.47 |
| **SAR1A** | 1.4 | 62.43 | 164.95 |
| USP28 | 1.42 | 8.62 | 23.02 |
| TXNDC15 | 1.42 | 11.87 | 31.82 |
| RSU1 | 1.44 | 39.84 | 108.35 |
| IFFO2 | 1.44 | 41.35 | 111.96 |
| LRRC28 | 1.45 | 8.46 | 23.14 |

| | | | |
|---|---|---|---|
| GLG1 | 1.45 | 39.24 | 106.94 |
| FAM203A | 1.46 | 9.86 | 27.07 |
| POLA2 | 1.46 | 14.33 | 39.51 |
| C4orf3 | 1.48 | 10.49 | 29.18 |
| TKT | 1.51 | 182.1 | 518.7 |
| MYCBP | 1.52 | 21.19 | 60.92 |
| CASP2 | 1.6 | 9.36 | 28.29 |
| VPS26B | 1.6 | 11.34 | 34.28 |
| FUS | 1.62 | 105.22 | 323.54 |
| CHST14 | 1.63 | 5.71 | 17.64 |
| **NNMT** | 1.63 | 278.47 | 861.62 |
| ST3GAL2 | 1.65 | 13.65 | 42.93 |
| ATG16L2 | 1.68 | 5.44 | 17.47 |
| GFOD2 | 1.68 | 5.51 | 17.65 |
| **CD109** | 1.71 | 3.77 | 12.3 |
| TRADD | 1.75 | 7.49 | 25.13 |
| MGAT5 | 1.75 | 8.02 | 27.05 |
| EFNA5 | 1.78 | 16.41 | 56.36 |
| **DTX4** | 1.8 | 5.98 | 20.76 |
| CALU | 1.8 | 57.9 | 201.02 |
| FMNL3 | 1.82 | 8.89 | 31.44 |
| FAM53B | 1.87 | 3.8 | 13.85 |
| GCLM | 1.88 | 5.48 | 20.15 |
| **CKAP4** | 1.93 | 40.66 | 154.95 |
| 8-Sep | 1.96 | 22.23 | 86.25 |
| TACC2 | 1.99 | 21.07 | 83.39 |
| PARP1 | 2 | 35.67 | 142.49 |
| NDRG3 | 2.01 | 13.73 | 55.41 |
| MLEC | 2.01 | 40.59 | 163.48 |
| SLC6A6 | 2.06 | 6.05 | 25.15 |
| TSPAN15 | 2.07 | 4.45 | 18.7 |
| SIDT2 | 2.08 | 9.12 | 38.47 |
| NBL1 | 2.12 | 15.75 | 68.67 |
| CARHSP1 | 2.12 | 43.46 | 189.57 |
| DNM1 | 2.19 | 10.93 | 49.76 |
| NAGA | 2.2 | 2.98 | 13.64 |
| RHOBTB1 | 2.21 | 11.75 | 54.32 |
| PGP | 2.23 | 4.51 | 21.13 |
| PCTP | 2.24 | 7.31 | 34.54 |
| SPARC | 2.25 | 926.15 | 4,402.49 |
| **PTGFR** | 2.3 | 4.2 | 20.73 |
| VANGL1 | 2.3 | 5.53 | 27.15 |
| IFT122 | 2.34 | 4.98 | 25.15 |
| CDK1 | 2.34 | 26.29 | 132.93 |
| CACYBP | 2.35 | 45.23 | 230.77 |
| DHFR | 2.36 | 4.57 | 23.4 |
| TMEM109 | 2.36 | 15.81 | 81.24 |
| TMEM50B | 2.41 | 6.83 | 36.38 |
| NDE1 | 2.43 | 7.81 | 41.94 |

| | | | |
|---|---|---|---|
| ASB13 | 2.44 | 4.41 | 23.87 |
| ZNF362 | 2.45 | 2.73 | 14.89 |
| DBNDD1 | 2.48 | 2.7 | 15.07 |
| SLC45A4 | 2.55 | 5.16 | 30.13 |
| **GSTM4** | 2.64 | 3.98 | 24.86 |
| SDC1 | 2.66 | 11.63 | 73.32 |
| PTGFRN | 2.68 | 11.35 | 72.97 |
| **STOM** | 2.69 | 21.87 | 141.44 |
| CNP | 2.73 | 20.67 | 137.11 |
| ARRB1 | 2.73 | 11.2 | 74.1 |
| MFI2 | 2.75 | 4.35 | 29.17 |
| PC | 2.76 | 6.31 | 42.79 |
| **PLEKHO2** | 2.77 | 1.97 | 13.46 |
| CACNB3 | 2.77 | 5.04 | 34.4 |
| CAMK2G | 2.81 | 4.8 | 33.75 |
| SLC1A4 | 2.84 | 3.68 | 26.39 |
| **C15orf39** | 2.9 | 3.45 | 25.7 |
| STMN1 | 2.91 | 106.58 | 802.47 |
| AGAP1 | 2.92 | 12.75 | 96.37 |
| GPX8 | 2.94 | 8.86 | 67.79 |
| **AGT** | 2.95 | 2.57 | 19.83 |
| **CRISPLD2** | 2.95 | 4.44 | 34.34 |
| **PAQR8** | 3 | 4.17 | 33.27 |
| SNN | 3.02 | 1.54 | 12.5 |
| CALR | 3.03 | 226.51 | 1,848.86 |
| THEM6 | 3.06 | 2.03 | 16.86 |
| SUFU | 3.07 | 4.36 | 36.66 |
| FGFR3 | 3.08 | 2.92 | 24.72 |
| MCM2 | 3.15 | 10.49 | 93.03 |
| HADH | 3.2 | 5.58 | 51.27 |
| SH3PXD2B | 3.23 | 2.2 | 20.57 |
| IL1R1 | 3.24 | 2.15 | 20.31 |
| SLC43A2 | 3.27 | 1.62 | 15.66 |
| RASSF4 | 3.28 | 2.16 | 20.94 |
| SNX10 | 3.31 | 6.14 | 61.03 |
| PLXDC2 | 3.38 | 3.68 | 38.31 |
| MKI67 | 3.42 | 9.77 | 104.86 |
| ZNF703 | 3.48 | 1.91 | 21.27 |
| VAV3 | 3.62 | 1.91 | 23.44 |
| **YPEL2** | 3.64 | 1.41 | 17.52 |
| BRI3BP | 4.05 | 1.06 | 17.49 |
| **IFIT1** | 4.15 | 3.76 | 66.87 |
| **SLC51A** | 4.21 | 4.36 | 80.71 |
| **SEMA5A** | 4.36 | 1.89 | 38.84 |
| ALDOC | 4.39 | 1.33 | 27.95 |
| RRM2 | 4.69 | 3.07 | 79.18 |
| IGFBP7 | 4.82 | 2.08 | 58.93 |
| IL34 | 4.84 | 1.13 | 32.25 |
| CLDN2 | 7.09 | 0.32 | 43.74 |

Columns indicate the target gene name, $log_2$ fold change (lytic/latent) value, latent iSLK.219 reads and lytic iSLK.219 reads. Bolded genes (n = 21) represent cellular genes that can escape KSHV SOX-mediated host shutoff and are up-regulated during lytic infection, and thus were selected for further analysis.

**Supplementary Table 5.** A list of predicted KSHV targets of miR-31-5p

| Target gene or gene cluster | Gene function | Site type | | | |
|---|---|---|---|---|---|
| | | total | 8mer | 7mer-m8 | 7mer-A1 |
| ORF4/6/7/8/9/10 | ORF4: complement control protein (immune evasion)<br>ORF6: single-stranded DNA-binding protein (viral replication)<br>ORF7: component of terminase that translocates viral genomic DNA into empty capsid during DNA packaging<br>ORF8: envelope glycoprotein (virion egress)<br>ORF9: viral DNA polymerase (viral replication)<br>ORF10: dUTPase (viral replication) | 3 | 1 | 2 | |
| ORF24 | Viral TATA box-binding protein (late gene expression) | 1 | 1 | | |
| ORF61/62 | ORF61: ribonucleotide reductase large subunit (viral DNA synthesis)<br>ORF62: capsid protein (viral capsid assembly) | 1 | | | 1 |
| K15 | Signal transducing membrane protein (virion production) | 1 | | | 1 |

Canonical site types: 8mer site: an exact match to miRNA positions 2-8 from the 5' end (the seed + position 8) with an A opposite position 1; 7mer-m8 site: an exact match to miRNA positions 2-8 from the 5' end (the seed + position 8); 7mer-A1 site: an exact match to miRNA positions 2-7 from the 5' end (the seed) with an A opposite position 1[9].

**Supplementary Table 6.** Oligonucleotides

Primer sequences used for PCR cloning the 3' UTRs of *KHDRBS3*, *SLC51A,* and *NNMT* into psiCHECK-2 vector (with *NotI* and *XhoI* sites)

| Name | Sequence (5' to 3') |
|---|---|
| KHDRBS3 F | TAAGCACTCGAGTTGTACTGTCTGATGTTGTG |
| KHDRBS3 R | TGCTTAGCGGCCGCTTTCTTTCAAAAG |
| SLC51A F | TAAGCACTCGAGGGTGGATGGCTTG |
| SLC51A R | TGCTTAGCGGCCGCCACATTTTCAG |
| NNMT F | TAAGCACTCGAGTGCCTGTGACCTCAATTAAAG |
| NNMT R | TGCTTAGCGGCCGCTTTACATGTAACAATTC |

Mutagenic primer sequences used to generate the mutant 3' UTRs of *KHDRBS3* and *SLC51A* into psiCHECK-2 vector

| Name | Sequence (5' to 3') |
|---|---|
| KHDRBS3 Mut F | CATATGCTAGTTTTTTTTTCAGAACGGTACATGCGTAAAAAGGGA |
| KHDRBS3 Mut R | TCCCTTTTTACGCATGTACCGTTCTGAAAAAAAAACTAGCATATG |
| SLC51A Mut F | GAATAAAATTGCTTTGGAAGAACGGTGGAAGGTGTTTTAAGTTTT |
| SLC51A Mut R | AAAACTTAAAACACCTTCCACCGTTCTTCCAAAGCAATTTTATTC |

Primer sequences used to analyze alternative splicing of ORF50/RTA, ORF57, and K8.1

| Name | Sequence (5' to 3') |
|---|---|
| ORF50 N5_E2 F | GGCTATACAGGACGGGTAATGATA |
| ORF50 N4 F | CTCCAATACCCGGAATTGGGA |
| ORF50 E1/N4_E2 F | GGGTGGCAGGACGGGTAAT |
| ORF50 N3_E2 F | CGTCCGGACGGGTAATGATA |
| ORF50 N4 Intronic R | CCTTGCGGAGTAAGGTTGACT |
| ORF50 E2 R | CAGGACCGCCGAAGCTTCTTA |
| ORF50 Total F | GAGTCCGGCACACTGTACC |
| ORF50 Total R | AAACTGCCTGGGAAGTTAACG |
| ORF57 UnSp F | AGTCCTCGTCTACAACAGACTTT |
| ORF57 UnSp R | GAGGAGGACACAGAGTCTGTAAT |
| ORF57 E1_E2 F | ACGGACAGGGATATCACCGC |
| ORF57 E2 F | GCATCCTAGAGGACTCTGTGTCC |
| ORF57 LE2 R | CGGTTTCTCGACGGCAACT |
| ORF57 E2_E3 R | GGTTTGGCAATCCCAGTACGC |
| ORF57 Total F | TGGACATTATGAAGGGCATCCTA |
| ORF57 Total R | CGGGTTCGGACAATTGCT |
| K8.1 UnSp F | CAATTGGGTAAACCGTCGGTGT |
| K8.1 Short F | CTCCGTCGAGATCATATTCATCT |
| K8.1 Long F | CAGCCTTTTCAGGATCATATTCA |
| K8.1 E2 R | TGCGCGTCTCTTCCTCTAGT |

qPCR primer sequences

| Gene Name | Forward Primer (5' to 3') | Reverse Primer (5' to 3') |
|---|---|---|
| ORF4 | GGCCAGGACAATGAAAAGTG | TCTCCAACTGCAATGTCACC |
| ORF6 | TGGTGGCTGAACTTGGTATG | TGCGTGCCAAACTTCTAGTG |
| ORF7 | TCAGACGACAAAACGCACAC | AGAGCATCAGCCACGTTTTC |
| ORF8 | ACCAACAACCAGGTGGAAAC | GGGCGATAAAAGTGTTCAGG |
| ORF9 | ACGTGCCATTTAGCTTCCAG | AAACGTTCACGCAGACACTG |
| ORF24 | AAGCCTGAGAAACGAACAGC | ACCGTTCCCAAATACAGACG |
| ORF37 | ACGTGGGACTTGGAAAACTG | GAAGGGGCGTTAATTTCCTC |

| | | |
|---|---|---|
| ORF50 (KSHV-encoded) | GAGTCCGGCACACTGTACC | AAACTGCCTGGGAAGTTAACG |
| ORF50 (Dox-inducible and KSHV-encoded) | CGCAATGCGTTACGTTGTTG | GCCCGGACTGTTGAATCG |
| ORF57 | TGGACATTATGAAGGGCATCCTA | CGGGTTCGGACAATTGCT |
| ORF61 | TTTCTTCCTGGCAGTTGGAC | AGAAGCGCTCAATGAACACC |
| ORF62 | CCTTTATCATGGCCACAACC | TGCCTCGTTGGTTTATCTCC |
| K15 | TTCCTGTAACTCCACCTCGTTC | CCACAATGACAACCACCTTG |
| PAN | ATAGGCGACAAAGTGAGGTGGCAT | TAACATTGAAAGAGCGCTCCCAGC |
| LANA N-terminus | GCGCCCTTAACGAGAGGAAGTT | TTCCTTCGCGGTTGTAGATG |

### References


1. Lidenge, S. J., Kossenkov, A. V., Tso, F. Y., Wickramasinghe, J., Privatt, S. R., Ngalamika, O., Ngowi, J. R., Mwaiselage, J., Lieberman, P. M., West, J. T. & Wood, C. Comparative transcriptome analysis of endemic and epidemic Kaposi's sarcoma (KS) lesions and the secondary role of HIV-1 in KS pathogenesis. *PLoS Pathogens*, **16**, e1008681 (2020).
2. Myoung, J. & Ganem, D. Generation of a doxycycline-inducible KSHV producer cell line of endothelial origin: maintenance of tight latency with efficient reactivation upon induction. *Journal of Virological Methods*, **174**, 12-21 (2011).
3. An, F. Q., Folarin, H. M., Compitello, N., Roth, J., Gerson, S. L., McCrae K.R., Fakhari, F. D., Dittmer, D. P. & Renne, R. Long-Term-Infected Telomerase-Immortalized Endothelial Cells: a Model for Kaposi's Sarcoma-Associated Herpesvirus Latency In Vitro and In Vivo. *Journal of Virology,* **80**, 4833-4846 (2006).
4. Love, M. I., Huber, W. & Anders, S. Moderated estimation of fold change and dispersion for RNA-seq data with DESeq2. *Genome Biology*, **15**, 550 (2014).
5. Robinson, M. D., McCarthy, D. J. & Smyth, G. K. edgeR: a Bioconductor package for differential expression analysis of digital gene expression data. *Bioinformatics*, **26**, 139-140 (2010).
6. Langmead, B. & Salzberg, S. L. Fast gapped-read alignment with Bowtie 2. *Nature Methods*, **9**, 357-359 (2012).
7. Li, B. & Dewey, C. N. RSEM: accurate transcript quantification from RNA-Seq data with or without a reference genome. *BMC Bioinformatics*, **12**, 323 (2011).
8. Luo, W., Friedman, M. S., Shedden, K., Hankenson, K. D. & Woolf, P. J. GAGE: generally applicable gene set enrichment for pathway analysis. *BMC Bioinformatics*, **10**, 161 (2009).
9. Agarwal, V., Bell, G. W., Nam, J. W. & Bartel, D. P. Predicting effective microRNA target sites in mammalian mRNAs. *eLife*, **4**, e05005 (2015).
